# Hypoxia-driven oncometabolite L-2HG maintains “stemness”-differentiation balance and facilitates immune suppression in pancreatic cancer

**DOI:** 10.1101/2020.05.08.084244

**Authors:** Vineet K Gupta, Nikita S Sharma, Brittany Durden, Vanessa T Garrido, Kousik Kesh, Dujon Edwards, Dezhen Wang, Ciara Myer, Sanjay K Bhattacharya, Ashok Saluja, Pankaj K Singh, Sulagna Banerjee

**Author notes:** Corresponding Author Address of Correspondence Sulagna Banerjee, PhD, Associate Professor, Department of Surgery, University of Miami, FL, USA 33156.

## Abstract

2-hydroxyglutarate (2-HG) has gained considerable importance in glioma and blood cancers that have mutations in the IDH1/2 gene. In the current study we show for the first time that pancreatic tumors produce 2HG in the absence of IDH1/2 mutation. Our study shows that hypoxic pancreatic tumors that have activated lactate dehydrogenase (LDH) activity, produce the L-isoform of 2HG.

Metabolic mass spectrometric analysis along with chiral derivatization showed that pancreatic cancer cells as well as stromal cells secrete the L-isomeric form of 2-hydroxyglutarate (L-2HG) when exposed to hypoxic environment. Serum analysis of human pancreatic cancer patients also showed similar accumulation of L-2HG. Our results showed that this abnormally accumulated L-2HG regulates self-renewal by increasing expression of genes associated with stemness (Sox-2, CD133) and by decreasing expression of differentiation genes (Pdx-1, HB9, NKX6.1). Further analysis showed that secreted L-2HG mediates cross talk with immune T-cells and hampers their proliferation and migration thereby suppressing the anti-tumor immunity. *In vivo* targeting of LDH enzyme with inhibitor (GSK2837808A) showed decrease in L-2HG as well as subsequent tumor regression and sensitization to immune-checkpoint therapy.

Present study shows for the first time that hypoxia mediated accumulation of L-2HG drives self-renewal in pancreatic cancer by shifting critical balance of gene expression towards stemness and promotes immune suppression by impairing T cell activation in this disease. Additionally, it indicates that targeting LDH can sensitize pancreatic tumors to anti-PD1 therapy by decreasing L-2HG and reverting their immune evasive function.

## Introduction

Metabolic reprogramming is a hallmark of cancer (Hanahan and Weinberg, 2011) that drives tumorigenesis. Over the years it has been observed that both cellular (fibroblasts and immune cells) or acellular (hypoxia) components of the tumor microenvironment are able to modulate tumor cells to undergo adaptive metabolic changes to survive and adapt to oncogenesis associated “stress” in the tumor (Boise and Shanmugam, 2019; Luo and Wang, 2019; McGinn et al., 2017; Nomura et al., 2016; Qian and Rankin, 2019). These altered metabolic pathways in response to the tumor microenvironment often lead to accumulation of metabolites that in turn can influence oncogenic signaling pathways, thus behaving as “oncometabolites”(Janke et al., 2015; Menendez et al., 2016; Nowicki and Gottlieb, 2015).

One such oncometabolite, 2-hydroxyglutarate, or 2HG, has gained considerable interest in cancers such as AML and glioma. In these cancers, a gain of function mutation in the isocitrate dehydrogenase (IDH1/2) enzyme results in accumulation of 2HG (Dang et al., 2009). 2HG, the reduced form of α−KG, is naturally present in two isomeric forms, D-2HG and L-2HG. In normal cells, 2HG is a minor by-product of metabolism. Its levels are maintained at a basal level in the cells by the activity of a conserved family of proteins, the L/D-2-hydroxyglutarate dehydrogenases (L2HGDH and D2HGDH) (Aghili et al., 2009; Linster et al., 2013), which convert 2HG to α−KG. Accumulation of this metabolite in AML and glioma inhibits both DNA (TETs) and histone (KDMs) demethylase, thereby hypermethylating gene promoters and regulating the transcriptional programming of genes involved in proliferation and growth (Chowdhury et al., 2011; Figueroa et al., 2010; Turcan et al., 2012; Xu et al., 2011). 2HG has been shown to be a strong oncogenic molecule that promotes cytokine independence and alters differentiation in hematopoietic cells, providing compelling evidence that a small molecule is sufficient to transform cells (Sciacovelli and Frezza, 2016). Further, studies have shown that 2HG in IDH1/2 mutated gliomas can suppress CD8+ infiltration(Kohanbash et al., 2017; Richardson et al., 2019), indicating that oncometabolites not only affect intracellular epigenetic changes but they can also affect tumor microenvironment.

Interestingly, while most of the studies on 2HG were on cancers with IDH1/2 mutations (for example, AML and glioma), there is recent evidence that under hypoxia there is accumulation of L-isoform of 2HG as a physiological response to oxygen limitation (Intlekofer et al., 2015; Intlekofer et al., 2017; Nadtochiy et al., 2016). Though hypoxia-induced production of L-2HG occurs independently of HIF1, it has been shown to be a result of hypoxic response (Intlekofer et al., 2015; Intlekofer et al., 2017). Accumulation of L-2HG decreases glycolysis and mitochondrial respiration by inhibiting regeneration of NAD^+^(Oldham et al., 2015). Furthermore, hypoxia-induced L-2HG promotes the same chromatin condensation as observed by the D-2HG produced by the IDH1/mutant cancers (Intlekofer et al., 2015; Kats et al., 2014) consistent with the well-established association between hypoxic niches and stem cell populations (Simon and Keith, 2008). Recent cell free studies by Intlekofer, et. Al. (Intlekofer et al., 2017) demonstrated that under hypoxia or acidic pH, a consequence of lactate accumulation in hypoxic condition, LDH and MDH enzymes produce the L-2HG in malignant cells. As observed with glioma cells, hypoxia induced L-2HG was reported to accumulate in CD8+ T-cells and affect their function(Tyrakis et al., 2016).

Pancreatic adenocarcinoma is among the most challenging solid tumors and is the 3^rd^ leading cause of cancer related death in United States alone(Siegel et al., 2019). Increased rate of tumor recurrence and resistance to most therapy is among the primary reasons for poor survival and prognosis in this disease. The robust fibro-inflammatory stroma in these tumors compress the blood vessels and render the tumor extremely hypoxic (Dauer et al., 2017; Feig et al., 2012). This not only prevents effective drug delivery in the tumor, but also triggers a survival adaptation in a population of tumor cells. Previous results from our laboratory show that under hypoxia, there is an enrichment of cells with increased self-renewal, that further contributes to therapy resistance in this disease (McGinn et al., 2017; Nomura et al., 2016). In addition, the pancreatic tumor microenvironment is extremely immune suppressive (Das et al., 2020; Gasser et al., 2017; Rao et al., 2018; Zhang et al., 2020). While heterogeneity of the tumors is one of the major causes for this, the interaction between the stromal components, the extracellular matrix, as well as the metabolic milieu of the microenvironment contributes to this phenomenon. Our recently published studies show that modulating metabolic pathways in the tumor can enhance efficacy of immune checkpoint inhibitor therapy in pancreatic tumors(Ho and Jaffee, 2020; Sharma et al., 2020). While the role of tumor microenvironment in metabolic reprogramming in pancreatic tumors has been acknowledged recently, there has been minimal focus on oncometabolites such as 2HG in this disease.

In the current study, we show that even though pancreatic cancers have no IDH1/2 mutations, they accumulate 2HG in the tumor tissue. Further analysis revealed that this is predominantly the L-isoform of 2HG, that is produced because of the hypoxic microenvironment in this cancer. Our studies also demonstrated that pancreatic stellate cells, the precursor to cancer associated fibroblasts in tumors, produce significantly higher L-2HG compared to the tumor epithelial cells. Our current studies additionally show that L-2HG mediates the balance between “stemness” and differentiation in pancreatic cancer by regulating histone methylation. Our results further show that inhibiting accumulation of L-2HG using a LDH inhibitor facilitated CD8+ infiltration in the pancreatic tumor and sensitized them to anti-PD1 therapy.

## Material & Methods

### Cells and reagents

Human pancreatic cancer cell lines MIA PaCa-2 (purchased from ATCC) and S2-VP10 (Gift from Dr. Masato Yamamoto’s lab, University of Minnesota) were used for present study. MIA PaCa-2 cells were cultured in DMEM (Gibco) containing 10% fetal bovine serum (FBS) and 1% penicillin/streptomycin antibiotic (Gibco) solution. S2-VP10 cells were cultured in RPMI 1640 containing 10% fetal bovine serum and 1% penicillin/streptomycin antibiotic (Gibco) solution. The primary mouse KPC cell line was isolated from the tumor of a 5- to 6-month-old genetically engineered mouse model of KRAS^G12D^p53^R172H^Pdx-1-Cre (KPC) mice using protocol from our previous study (Sangwan et al., 2015). Mouse cancer associated fibroblasts (CAFs) were isolated from tumor bearing KPC mice using the protocol described by Sharon et al.(Sharon et al., 2013). The purity of isolated fibroblasts was verified by flow cytometry using fibroblast surface protein (FSP)+ and CK19-population as marker for fibroblasts. Full grown monolayer cultures of all the cell lines were trypsinized for 5 min (0.25% trypsin-EDTA), harvested and passaged several times for expansion. Cells were cultured in a humidified atmosphere with 5% CO_2_ and at 37° C and were passaged every 72 hours. For hypoxia, cells were cultured at 37° C in hypoxic incubator with 1% O_2_, 5% CO_2_ and 94% N_2_ for 48 hours. All the established cell lines were used from passages 5–20. The RRID of all reagents used in this study can be found in Supplementary Data.

### Quantification of 2-Hydroxyglutarate enantiomers

2HG D/L isomer analysis was done at the Comprehensive Metabolomics Core at University of Michigan according to their established protocol. Follow up identification of L-2HG was done according to the protocol mentioned in (Intlekofer et al., 2015; Intlekofer et al., 2017).

### Colony formation assay

Human pancreatic cancer cells were cultured in normoxic or hypoxic environment for 48 hours. To see the effect of L-2HG on colony formation cells were cultured in the presence of octyl L-2HG at mentioned concentration for 48 hrs. After treatment cells were counted and plated at a density of 1000 cells per well. Colonies were counted after 4 days of plating. The results are presented as no. of colonies from 1000 cells per well.

### iTRAQ proteomics analysis

Cells cultured in normoxia, hypoxia and in the presence of octyl L-2HG for 48hrs were used for iTRAQ MS/MS analysis. Cells were centrifuged at 300g at and resuspended in 30⍰μL of dissolution buffer (0.5⍰M triethylammoniumbicarbonate (TEAB), pH 8.5). Lysed samples were denatured with 2% SDS and reduced with 100⍰mM tris-(2-carboxyethyl) phosphine (TCEP). Samples were incubated for 1⍰hr at 60° C, and then 1⍰μL of iodoacetamide solution (84⍰mM) was added and incubated at 30° C in the dark. Promega Sequencing Grade trypsin was added to each sample, and then incubated at 37° C overnight. 8Plex iTRAQ reagents were prepared the following day by adding 50⍰μL isopropanol to each reagent. Each reagent was then combined in one tube. Sample-reagent mixtures were incubated for 2⍰hr at room temperature, and then the reactions were quenched with water, and combined into one tube. Sample was then vacuum dried and washed three times with water. Dried sample subsequently used for MS/MS analysis. Analysis of identified proteome was done using Ingenuity Pathway Analysis and String DB (Szklarczyk et al., 2019).

### Quantitative Real-time PCR

Quantitative real-time PCR for stemness and differentiation genes was performed using primers purchased from Qiagen (QuantiTect primer assay). RNA was isolated from cells according to the manufacturer’s instructions using Trizol (Invitrogen). The High Capacity cDNA Reverse Transcription Kit (Life Technologies) was used to prepare cDNA from 2 µg of RNA. Subsequently, real-time PCR was performed using the SYBR Green from Roche (Lightcycler 480 SYBR Green I Master) as per the manufacturer’s instructions using a Light Cycler II (Roche). All data were normalized to the housekeeping gene 18S (18S QuantiTect Primer Assay; Qiagen).

### Histone methylation analysis

For histone methylation analysis, histone proteins were isolated from cells by using histone extraction kit (Abcam - ab113476) according to the manufacturer’s protocol. Isolated histone protein concentration was measured using the BCA protein estimation assay (Thermo Scientific). Equal amount of protein was then separated on an SDS PAGE and transferred to nitrocellulose membrane. Blots were probed with antibodies against various histone methylation.

### ChIP assay

MiaPaCa-2 cells were treated with octyl L-2HG or hypoxic environment for 48 hr. After, cells were collected, and CHIP assay was performed by using ChIP kit (Abcam Cat No; AB500) as per the manufacturer’s instructions. After ChIP, DNA levels were measured by real-time quantitative PCR. Primers used for QPCR are as follows:

hPDX1-ChIP-FW TTTTAAGCCACGGTGAAAATG

hPDX1-ChIP-Rv CTAAGAGGCTAGGCCCAGGT

hCD133-Fwd: TTTCCAAATCCAGGAGCAAC

hCD133-Rev: ACAGGAAGGGAGGGAGTCAT

### T cell proliferation assay

Splenocytes were isolated from the spleen of healthy 6-8 week old C57BL6/J mice as follows; spleens were isolated from euthanized mice under sterile environment and crushed by using syringe plunger in 70µm cell filter. Subsequently, RBCs were lysed using RBC lysis buffer and washed using sterile PBS. Subsequently, T cells were isolated from spleen using MojoSort™ Mouse CD3 T Cell Isolation Kit (Biolegend Cat no; 480024) as per the manufacturers protocol. Cells were labeled with CFSE (ThermoFisher cat no: C34554) as per the manufacturer’s protocol. In short, 2⍰×⍰10^7^ cells (20ml PBS) were incubated with CellTrace CFSE staining solution to make final concentration of 5µM and incubated at 37° C for 20 min, and then washed extensively and were left overnight at 37°C 5% CO2 in culture medium. Some cells were used for flow cytometry to check CFSE labeling by flow cytometry. Next day CFSE labeled cells were incubated with 50 µL Invitrogen Dynabeads mouse T-Activator CD3/CD28 per 1 mL of cells for T cell activation. Same time cells were also treated with 1mM Octyl L-2HG in one group of cells. Before acquisition in the flow cytometer, cells were labeled with mouse CD4-APC cy7 (Caltag), TCRb PE cy7 (Caltag), and L/D dye, to differentiate CD4^+^ and CD8^+^ T cells. Flow cytometry analysis was performed before the beginning of the culture and at day 5 and 6⍰using LSRII (BD).

### T cell migration assay

T cell migration was evaluated using an 5.0-μm pore size 24-well, Transwell plate (Corning) as described previously (Zhang et al., 2006). In short, magnetically sorted T cells were left overnight at 37°C 5% CO2 in culture medium. Next day T cells were treated with either 0mM or 1mM Octyl L-2HG for 48 hrs. Cells cultured in the absence of Octyl L-2HG were used as controls. After 48 hrs T cells were washed once with RPMI 1640 medium and counted. 1000 cells resuspended in 100 µl of culture media were placed in the top chamber of the Transwell chamber. Bottom chamber of Transwell plate received chemokine in 500 μL of T cell culture medium. After 90 min incubation at 37° C in a 5% CO2 atmosphere, the top chamber was removed, and the number of T cells that had migrated into the bottom chamber was counted under the microscope.

### Immunohistochemistry

For immunofluorescence, paraffin tissue sections of tumors were deparaffinized in xylenes and hydrated gradually in ethanol. Antigen retrieval was done by steaming the slides in 1X Reveal Decloaker (Biocare Medical) to minimize background staining. Background sniper solution (Biocare Medical) was used for blocking as well as antibody dilution. Primary antibodies for CD133, PDX-1 (Cell signaling) and H3K9me3 (cell signaling) were diluted as per manufacturer’s instruction and incubated overnight at 4°C. Subsequently, secondary antibody conjugated with fluorophore were used. Slides were counterstained with DAPI and visualized in a Leica fluorescent microscope. Tissue samples were also incubated with respective isotype controls and did not show any nonspecific staining. TUNEL assay was performed using TUNEL Assay Kit - HRP-DAB (Abcam-ab206386) according to the manufacturer’s protocol.

### Animal experiment

All animal experiments were performed according to the University of Miami Animal Care Committee guidelines. Female C57BL/6 mice (Jackson Laboratory) between the ages of 4 and 6 weeks were used for orthotopic implantation of KPC and CAF cells. 1000 KPC and 9000 CAFs suspended in 100% Corning Matrigel GFR Basement Membrane Matrix were implanted orthotopically in tail end of mouse pancreas. Tumors were allowed to grow for 10 days, after which mice were randomized and treatment was started. One group of mice received GSK2837808A (LDHA Inhibitor) at a dose of 6 mg/kg/QD orally while another group received vehicle control. After 4 weeks of treatment mice were euthanized and pancreatic tumors were harvested, dimensions were noted, and subsequently used for flow cytometry or immunohistochemistry. To see the effect of CD8 cells in GSK2837808A (LDHA Inhibitor) mediated tumor reduction, CD8-KO mice (B6.129S2-Cd8atm1Mak/J females) were purchased from Jackson Laboratory. 1000 KPC and 9000 CAFs were implanted orthotopically in tail end of pancreas and subsequently treated with above mentioned drug regimen. For evaluating the combinatorial effect of LDHA inhibition and immunotherapy on tumor growth, after tumor implantation mice received anti PD-1 ab (10mg/Kg BW), anti CTLA4 ab (10mg/Kg BW) alone or in combination with GSK2837808A (6 mg/Kg/QD). Immunotherapy was given intraperitoneally on day 15, 17 and 19 post LDHA inhibitor treatment and vehicle control group received isotype control antibody at the same time.

### Flow-cytometry for tumor infiltrated immune cells

Tumor samples harvested from mice were placed in RPMI until they were ready to be processed. Tumors were minced into tiny pieces and then digested with Collagenase IV at 37° C for 2 hours. After the tissue digestion, they were passed through a 70μm nylon filter. The tissue was then spun at 500g for 5 mins. RBCs in tumor were lysed by RBC lysis buffer. After RBC lysis cells were resuspended in 1ml of flow buffer (0.5% Bovine Serum Albumin,2mM EDTA, 1% Penicillin-Streptomycin in 500mL PBS) and stained for cell surface marker at 4 degree for 15 mins. The tubes were then washed and spun at 500g for 5 mins. 100μl of cytofix - cytoperm buffer was added to the pellet and the cells were allowed to fix for an hour at room temperature. After 1 hour, the cells were then washed with 1 ml of FACS buffer and then centrifuged at 500g for 5 mins. The supernatant was then discarded, and the cells were stained with the intercellular antibody for 40 mins in the dark. After 40 mins of staining, the cells were washed with 1ml of FACS buffer and then spun at 500g for 5 mins. Cells were then resuspended in 200μl of FACS buffer and were acquired by using BD LSRII at flow core facility at Uni of Miami. The antibodies used for immune cell analysis were all from Biolegend-CD3 PE Dazzle (catalog 100348), CD4 PE Cy7 (catalog 100528), CD49b FITC (Catalog 108906), CD25 AF-700 (catalog 102024), TCRγ/δ BV 510 (catalog 118131), FoxP3 PE (Catalog 126404), IL-4 BV-711 (catalog 504133), IL-17 BV-421 (catalog 506926), TNF-a BV-650 (Catalog 506333), and IL10 APC/Cy7 (catalog 505036). Data was acquired and analyzed with FACSDiVa software (BD Biosciences) and FlowJo Software.

### Statistical Analysis

All the *in vitro* experiments were performed independently at least three times and values are expressed as mean ± standard error of mean (SEM). The significance of the difference between any two samples was analyzed by student t-test using GraphPad Prism software and values of p<0.05 were considered statistically significant. For experiments with more than two groups, 1-way ANOVA test was performed for statistical analysis.

## Results

### Pancreatic cancer cells and stromal cells produce L-2 HG when exposed to hypoxic microenvironment due to increased LDH/MDH activity

Mutations in Isocitrate Dehydrogenase (IDH) gene results in accumulation of D-isoform of 2-hydroxyglutarate (D-2HG), an oncometabolite responsible for hypermethylation of DNA and histones, thereby affecting gene transcription. Recent research showed that hypoxia induces production of L-isoform of 2HG (L-2HG) as a physiological response to oxygen limitation. Analysis of mutations in the TCGA database (www.cbioportal.org) showed that pancreatic tumors lacked IDH mutations even though they had elevated expression of IDH1 and IDH2 (**Figure 1A**). However, serum samples analysis from pancreatic cancer patients revealed an accumulation of 2HG (**Figure 1B**). Further analysis of the enantiomers showed that the serum 2HG was predominantly the L-isoform (**Figure 1C**). Pancreatic tumors are notoriously hypoxic. To see if hypoxia was responsible for the accumulation of L-2HG, we subjected pancreatic cancer cell lines (MIA-PaCa2 and KPC001) to 1% oxygen. Analysis of the conditioned media showed an increase in L-2HG in pancreatic cancer under hypoxia in both cell lines (**Figure 1D**). Similarly, upon analysis of the cell lysates, we observed that under hypoxia both MIA-PaCa2 and KPC001 produced more L-2HG (**Figure 1E**). Further, upon comparing D- to L-isoforms of 2HG (whether from conditioned media or from cell lysate), we observed that pancreatic cancer cells produced significant L-2HG while almost no D-2HG was being produced. As observed with tumor epithelial cells, pancreatic stellate cells when cultured under hypoxia, produced significantly high L-2HG production compared to D-2HG (**Figure 1F**). Interestingly, while pancreatic stellate cells secreted almost 10 times more L-2HG when compared to epithelial cells like MIA-PaCa2, their cell lysates showed no significant different levels of L-2HG under hypoxia (**Figure 1G**). Furthermore, the while pancreatic epithelial cells as well as stellate cells produced L-2HG, there was almost no D-2HG produced by these cells (**Figure 1D-G**).

**Figure 1:**
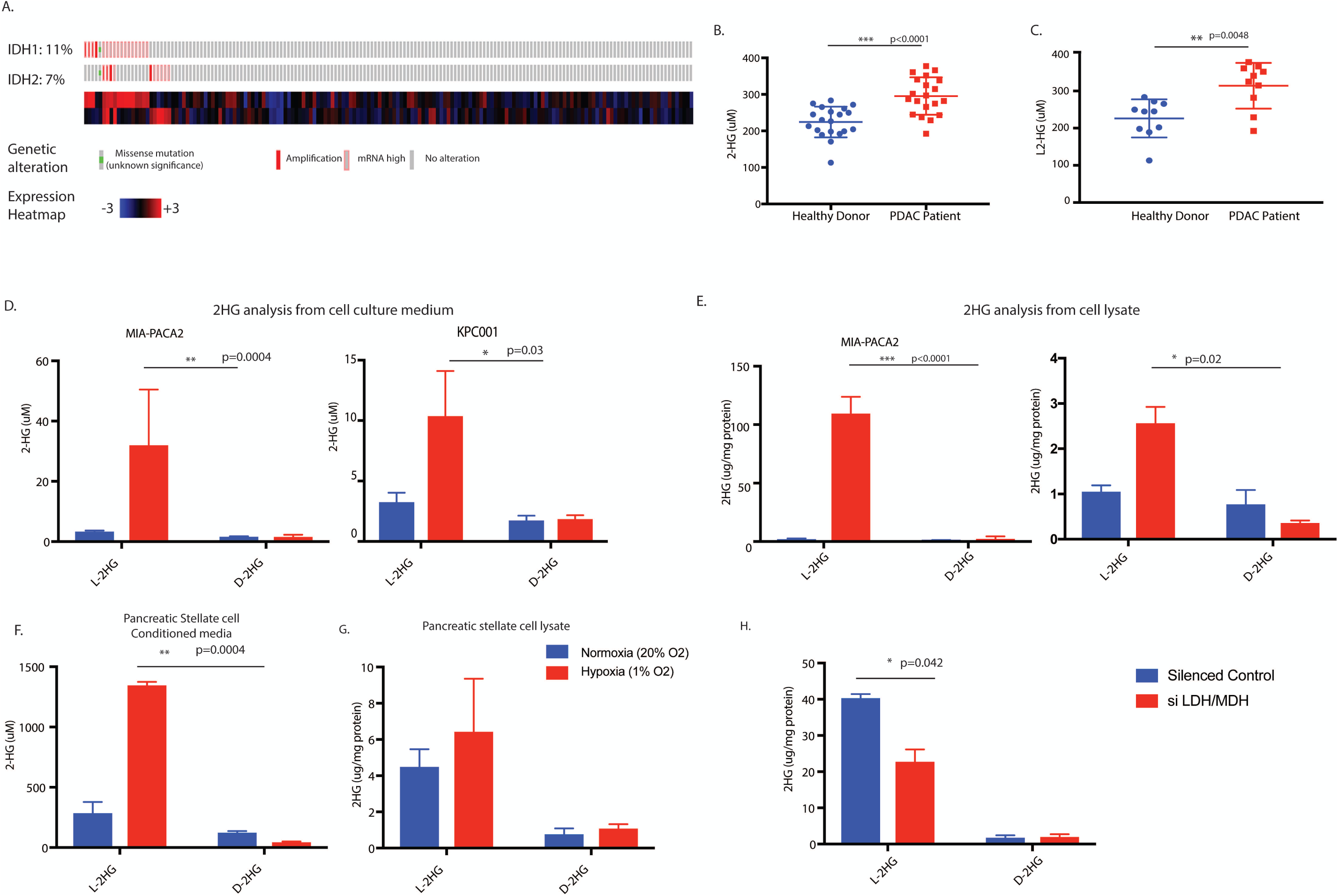
Pancreatic cancer and stromal cells produce L-2HG under hypoxic environment. Mutation analysis of Pancreatic cancer patients in TCGA database showed lack of IDH1/2 mutation (A). Serum sample analysis of pancreatic cancer patients showed accumulation of 2-HG (B). Isomeric analysis showed L-isomeric form predominantly exist in patient serum sample (C). Pancreatic cancer cell lines showed increased L-2HG in condition media (D) and cell lysates (E) when grown in hypoxic environment. Pancreatic stellate cells predominantly secrete L-2HG when grown in hypoxia (F) with no accumulation inside the cells (G). Silencing LDH and MDH enzymes results in decrease in L-2HG when grown under hypoxia (H). * indicate p-value </= 0.05

Since L-isoform of 2HG is produced as a result of LDH and MDH enzyme activity under hypoxia (Intlekofer et al., 2015), we next silenced these enzymes and estimated the 2HG levels in the cells as well as conditioned media. Our results showed that while individual silencing of LDHA, LDHB, LDHC, MDH1 and MDH2 resulted in modest effect on the gene expression, a cocktail of LDH/MDH siRNA significantly decreased the L-2HG levels in these cells (**Figure 1H**).

### L-2HG decreases transcription of differentiation genes

In cancer cells, “stemness” is considered a balance between the ability of the cells to self-renew and suppression of its differentiation potential (Aponte and Caicedo, 2017; O’Brien et al., 2010). In the current study, we studied the expression of genes involved in differentiation following exposure to hypoxia at 1% oxygen. Our results showed that along with increase in the stemness genes as seen previously (McGinn et al., 2017; Nomura et al., 2016), hypoxia also suppressed the expression of genes involved in differentiation (**Supplementary Figure 1A**,**B**) and increased CD133+ tumor initiating population (**Supplementary Figure 1C**). Since our data showed that L-2HG accumulated under hypoxia, we next checked the expression levels of genes involved in differentiation and stemness in pancreatic cancer following treatment with cell-permeable octyl L-2HG. Our results showed that as observed with hypoxia, upon treatment with octyl L-2HG, expression of genes regulating stemness (Sox2, CD133) were increased along with an increase in the CD133+ population (**Figure 2A**) while those involved in differentiation were decreased (**Figure 2B**). Flow cytometry analysis also showed increase in CD133+ cancer stem cells when pancreatic cancer cells were grown in presence L-2HG (**Figure 2C**). To establish the causality that hypoxia induced accumulation of L-2HG was responsible for this, we next silenced the LDH and MDH using siRNA and studied the gene expression of the same panel. Our results showed that a cocktail of LDH/MDH siRNA significantly downregulate expression of stemness genes (**Figure 2C**) and increased the expression of differentiation genes (**Figure 2D**). To further validate this, and study if it was affecting the self-renewal potential of the cancer cells, we performed a colony formation assay at limiting dilution, a surrogate assay for tumorigenicity. Our results showed that like exposure to hypoxia, octyl L-2HG treated cells formed colonies at very low dilutions in both MIA-PACA2 (**Figure 2E**) and well as S2VP10 cells (**Supplementary Figure 2A**,**B**). To study if the “stemness” and differentiation was being induced in epithelial cells as a result of the L-2HG produced by the pancreatic stellate cells (PSC), we next treated the pancreatic cancer cells (MIA-PaCa2) with conditioned media from stellate cells (cultured under hypoxic conditions). Our studies showed that as observed with Octyl L-2HG treatment, conditioned media from PSC also increased expression of stemness genes and suppressed the expression of differentiation genes (**Figure 2F**).

**Figure 2:**
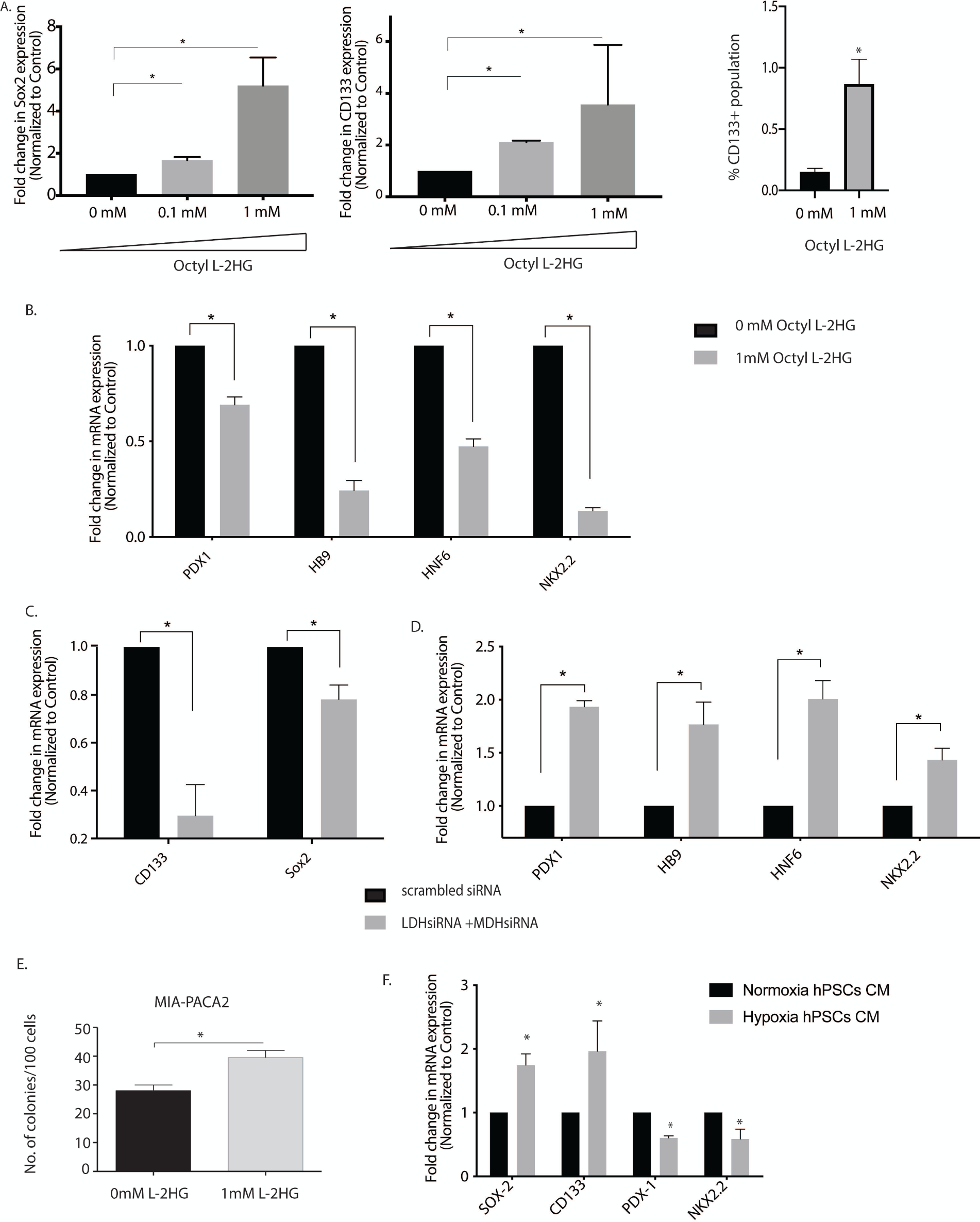
L-2HG promote self-renewal in pancreatic cancer cell by modulating stemness and differentiation gene expression. Octyl L-2HG treatment results in increase in stemness genes expression (A) and decrease in differentiation genes expression (B). Flow cytometry analysis showed increase in CD133+ population upon Octyl L-2HG treatment (C). Silencing LDH and MDH enzymes under hypoxic environment downregulate expression of stemness genes (D) and increased the expression of differentiation genes (E). Pancreatic cancer cells form higher number of colonies when incubated with Octyl L-2HG compared to control (F). Condition media of Pancreatic stellate cells grown under hypoxia upregulate stemness gene expression and downregulate differentiation genes expression (G). * indicate p-value </= 0.05

### Hypoxia and L-2HG affect proteome to upregulate chromatin modification pathways

To study if hypoxia and L-2HG were affecting the same signaling pathway and thus working in series, we next performed an iTRAQ proteomic analysis after growing pancreatic cancer cells under hypoxia as well as after treatment with Octyl L-2HG. The abundance ratio of the identified protein compared to their respective control (normoxia for cells grown in hypoxia and untreated for cells treated with Octyl L-2HG) is shown in **Figure 3A.** We observed that a distinct proteome alteration at hypoxia and after treatment with octyl L-2HG. Using a cut-off score of 1.1 fold in which samples with an abundance ration >1.1 were considered upregulated and samples with an abundance ratio <0.9 were considered downregulated. We next used the sample set for 1mM L-2HG treated pancreatic cancer cells and analyzed them using the STRING database. STRING identifies protein-protein interaction and thus organizes the protein dataset into clusters with similar functions. Our results showed distinct grouping of proteins involved in RNA processing, stress adaptation, histones along with clusters of cellular metabolism and actin reorganization (**Figure 3B**). Similar clustering was also observed in the proteome of samples collected under hypoxia (**Supplementary Figure 3A**). Interestingly, both hypoxia as well as octyl L-2HG treated samples showed same canonical pathways affected in a Ingenuity pathway analysis (**Figure 3C, Supplementary Figure 3B**), indicating that both hypoxia and L-2HG affected the same pathways. Among the top altered pathways were glycolysis, gluconeogenesis and unfolded protein response under both hypoxia as well as octyl L-2HG treatment. While these were expected changes under hypoxia, the observation that these pathways were altered with octyl L-2HG treatment were interesting, indicating that accumulation of L-2HG under hypoxia influenced cellular machinery and stress response pathways as well. In addition, the proteomic analysis also identified components from the EIF2 signaling pathway and Sirtuin signaling pathway under these conditions. Interestingly, according to literature, both of these pathways are involved in chromatin remodeling as well as transcriptional repression in cells (Baird and Wek, 2012; Seto and Yoshida, 2014).

**Figure 3:**
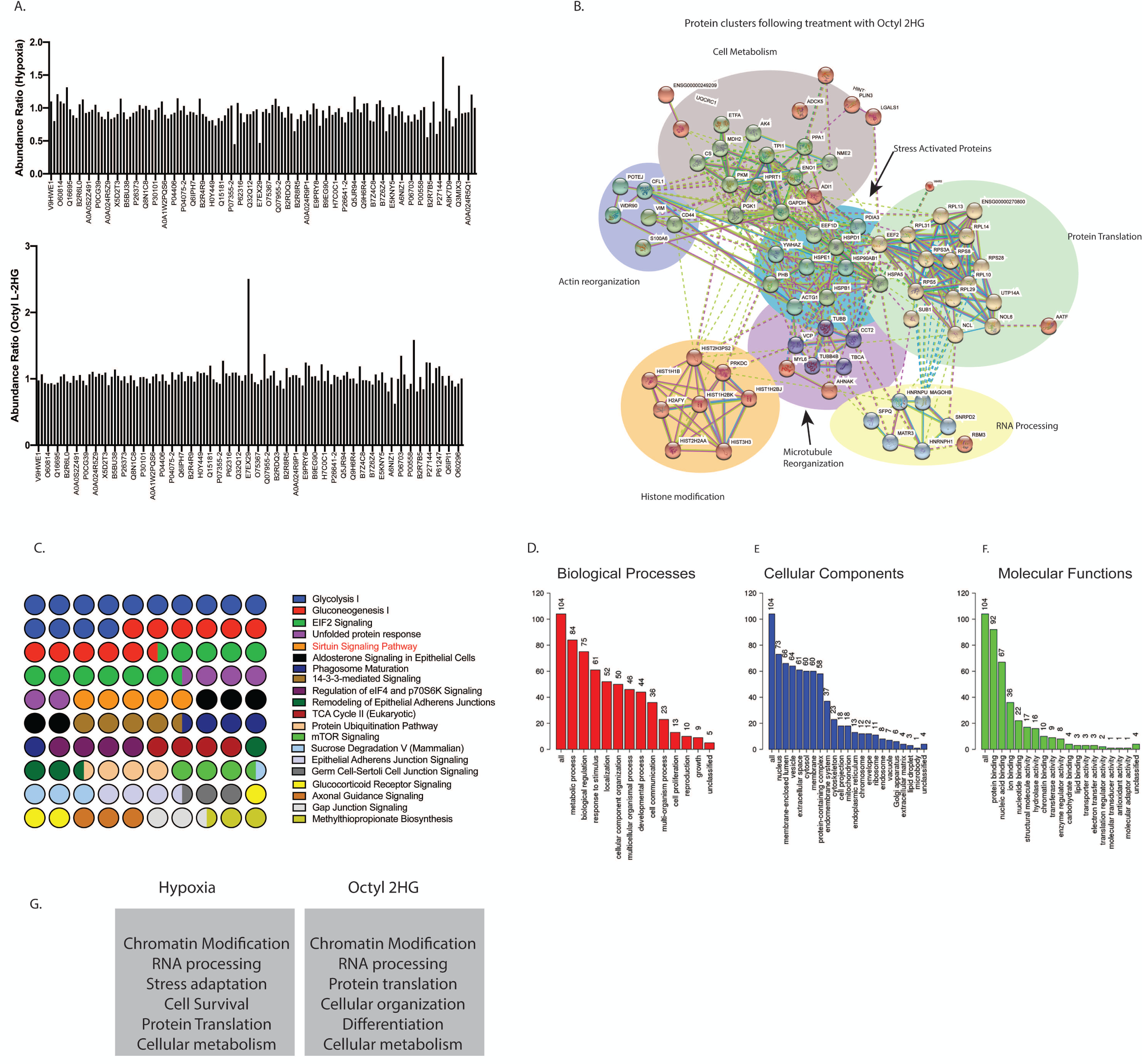
Hypoxia and L-2HG treatment regulate proteome as shown by ITRAQ analysis. The abundance ratio of the identified protein compared to their respective control (normoxia for cells grown in hypoxia and untreated for cells treated with Octyl L-2HG) seen in (A). Analysis of identified proteins through the STRING database showing distinct grouping of proteins involved in RNA processing, stress adaptation, histones along with clusters of cellular metabolism and actin reorganization (B). IPA analysis of canonical pathways represented in the proteome of cells treated with octyl L-2HG (C). The gene ontology analysis showed metabolic processes, response to stimulus and localization among the major biological processes affected by both hypoxia and octyl L-2HG (D),major cellular components (E) and molecular functions (F). Ingenuity Pathway Analysis identified top 5 networks in both the hypoxia as well as octyl L-2HG proteome, that indicated chromatin modifying proteins were enriched upon L-2HG treatment as well as hypoxia (G).

The gene ontology analysis showed metabolic processes, response to stimulus and localization among the major biological processes affected by both hypoxia and octyl L-2HG (**Figure 3D, Supplementary Figure 3C**). Similarly, nuclear proteins were among the major cellular components affected by these treatments (**Figure 3E**). When analyzed for molecular functions, protein and nucleic acid binding seemed to be the major functions under hypoxia and octyl L-2HG treatment. Interestingly, chromatin modification appeared to be among 10% molecular function of proteins under hypoxia and octyl L-2HG treatment (**Figure 3F**). Ingenuity Pathway Analysis also identified top 5 networks in both the hypoxia as well as octyl L-2HG proteome, that indicated chromatin modifying proteins were enriched upon L-2HG treatment as well as hypoxia (**Figure 3G, Supplementary Figure 3D**). The full list of identified peptides is presented in **Supplementary Table 1**.

### Hypoxia induced L-2HG deregulated histone demethylation to suppress pro-differentiation gene and activate stemness genes

Since 2HG is known to regulate the activity of chromatin modifying enzymes, specifically histone methylases, to affect the transcription of genes and as hypoxia was resulting in accumulation of L-2HG, we next studied the global effect on histone methylation status. Our results showed that hypoxia resulted in a global increase in methylation of H3K4, H3K9 and H3K27 (**Supplementary Figure 4 A, B, C**). Similarly, treatment with octyl L-2HG resulted in hypermethylation of H3K4, H3K9 and H3K27 globally (**Figure 4A**). To study the effect of hypermethylation specifically on the stemness and differentiation genes, we chose CD133 as our “stemness” gene and PDX1 as our differentiation gene. We next performed a histone ChiP with H3K4, H3K9 and H3K27 on promoters of both CD133 as well as PDX1 genes with mono/tri methylation specific antibodies. Our results showed that under hypoxia H3K9 was trimethylated in PDX1 (**Figure 4B**), while in CD133 it was mono-methylated (**Figure 4C**). Interestingly, PDX1 was not monomethylated in hypoxia (Supplementary Figure 5A) and CD133 was not trimethylated in hypoxia (**Supplementary Figure 5B**). Mono-methylation of H3K9 is an indicator of gene expression, while trimethylation of H3K9 is associated with repression of gene expression. This data showed that under normoxia, PDX1 expression was “on” while CD133 expression was baseline, while under hypoxia, CD133 expression was turned “on” while PDX1 became inactive. We next validated this observation by treating pancreatic cancer cells with octyl L-2HG. As seen with hypoxia, treatment with octyl L-2HG also showed trimethylation of H3K9 at the PDX1 promoter (**Figure 4D**) and mono-methylation of H3K9 at the CD133 promoter (**Figure 4E**) resulting in PDX1 repression and upregulation of CD133 gene expression compared to untreated observed in Figure 2A, B. Methylation status of H3K27 and H3K4 were not significantly altered in these two genes (**Supplementary Figure 5 C, D**). To further validate that this change in methylation status was due to altered enzyme activity of LDH and MDH, we next silenced LDH/MDH in cells under hypoxia and looked for global methylation. Inhibition of LDH/MDH also showed a decrease in global methylation (**Figure 4F**) as well as CD133 gene repression and PDX1 gene activation observed in Figure 2 C, D.

**Figure 4:**
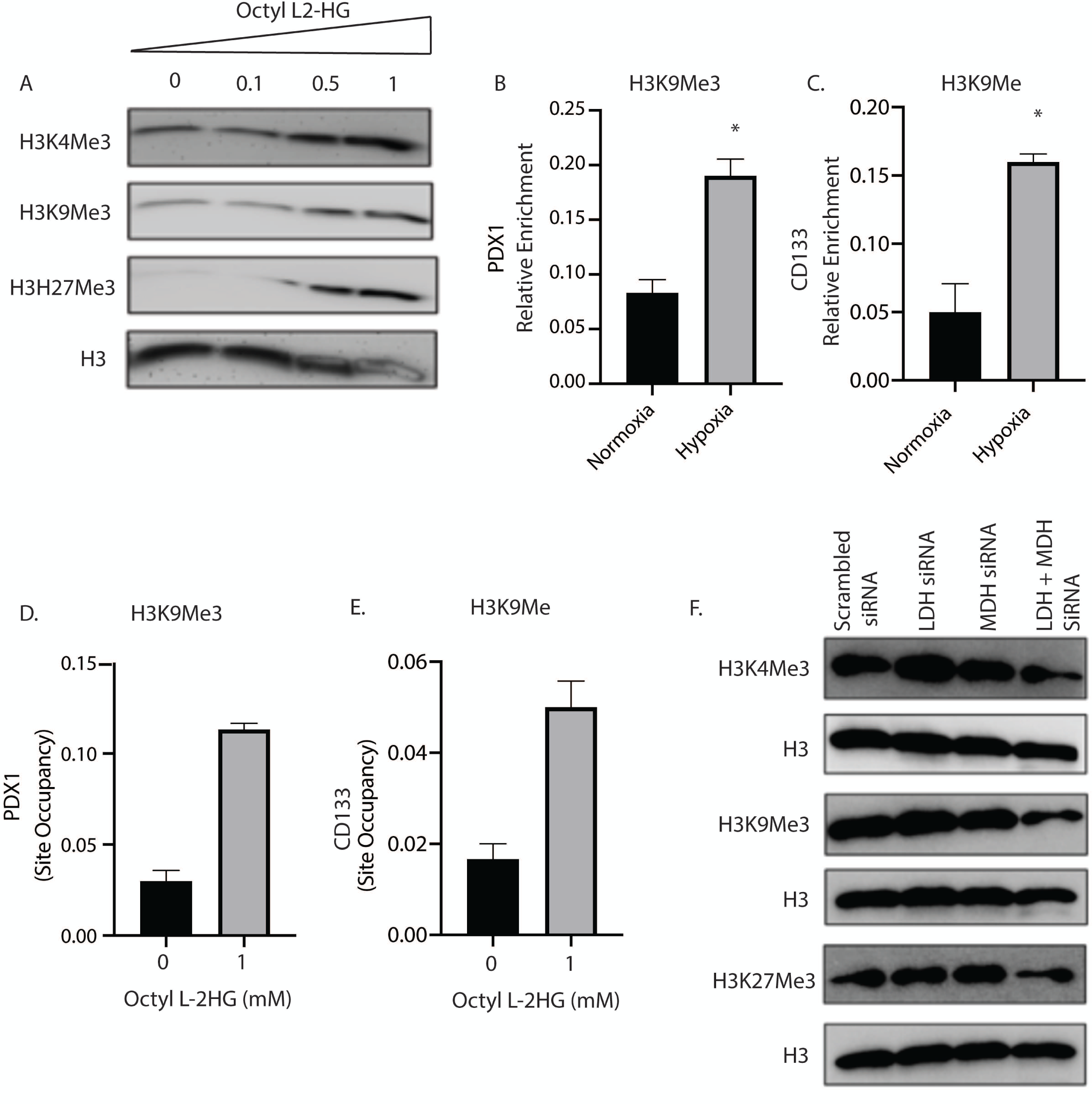
L-2HG drive hypoxia mediated epigenetic changes and regulate stemness and differentiation gene methylation status. Octyl L-2HG treatment results in increase in histone methylation (A). CHIP QPCR analysis showed that in hypoxic cells H3K9 was trimethylated at differentiation gene (PDX-1) promoter (B) and monomethylated in stemness gene (CD133) promoter (C). Similarly, Octyl L-2HG treated cells were trimethylated at PDX-1 promoter (D) and monomethylated at CD133 gene promoter (E). Silencing of LDH and MDH enzyme promote demethylation of histones as shown by western blot analysis under hypoxia (F). * indicate p-value </= 0.05

### LDH inhibition in vivo decreases tumor volume and histone methylation

Pancreatic tumors are known to be hypoxic and thus have an increased LDH activity(Nomura et al., 2016). To study if inhibition of LDH affected the L-2HG in the tumors as well as systemically, we implanted murine pancreatic cancer cell line KPC001 with CAF cells in the pancreas of a C57BL6 mouse. The tumors were treated with a LDH inhibitor GSK2837808A (6 mg/Kg/QD). Following 4 weeks of treatment the animals were sacrificed and a necropsy was performed. Our results showed that there was in significant decrease in the tumor weight (**Figure 5A**) and tumor volume (**Figure 5B**). Additionally, TUNEL staining on the tumor showed increased apoptotic cells in the treated group compared to control (**Figure 5C**). We next analyzed L-2HG levels in the plasma from both treated and untreated group. While L-2HG was significantly decreased in the plasma (**Figure 5D**), we did not observe any significant difference in the tissue level of L-2HG (**Supplementary Figure 5E**). Even though L-2HG levels did not significantly decrease in the tumor tissues, analysis of global histone methylation also showed that inhibition of LDH decreased the total methylation compared to control (**Figure 5E**). Interestingly, upon treatment with LDH inhibitor, there was a significant increase in the expression of differentiation marker Pdx1 (**Figure 5F**) and decrease in the expression of CD133 (**Figure 5G**) consistent with our *in vitro* observation.

**Figure 5:**
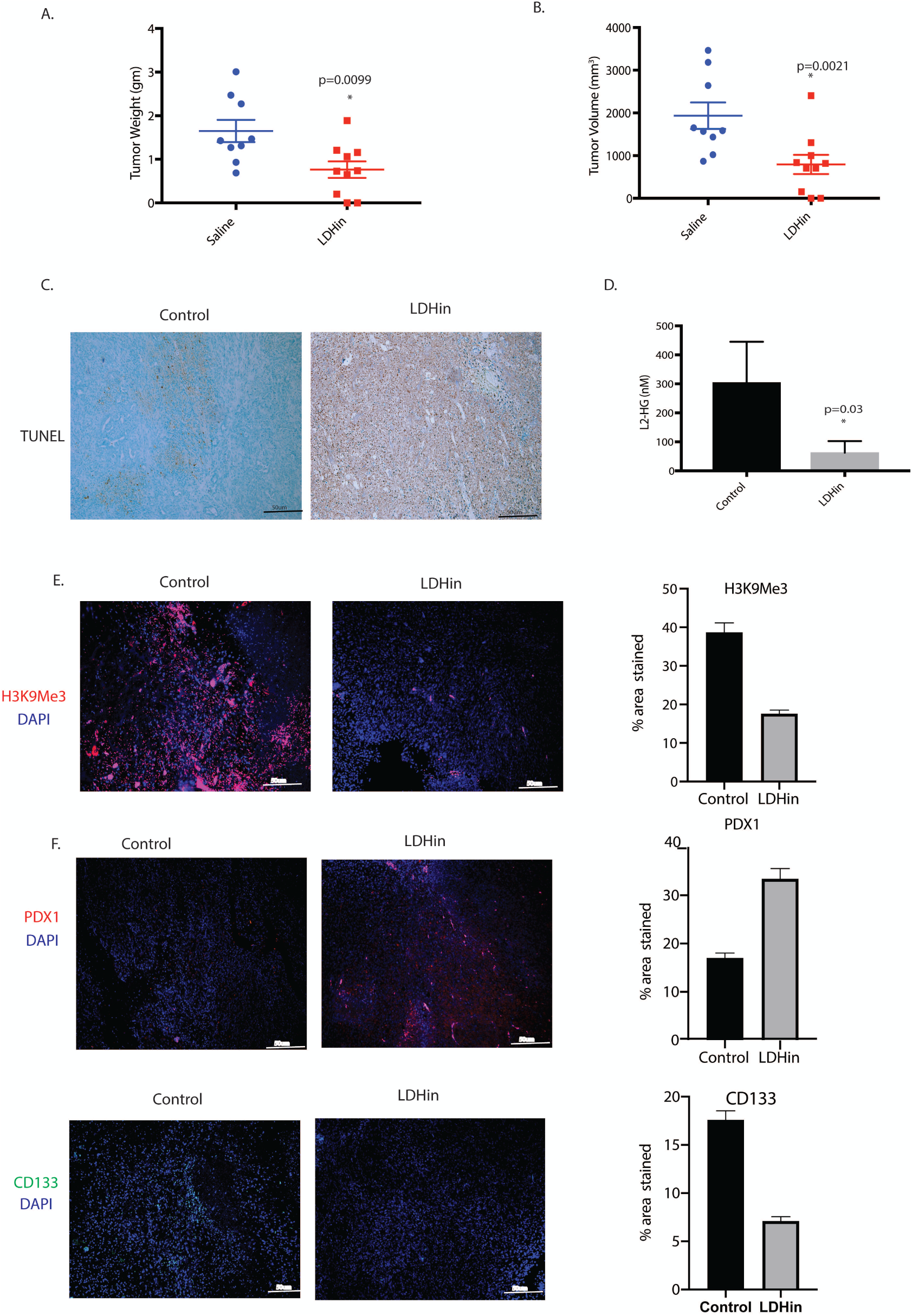
In vivo inhibition of LDH reduces tumor burden and histone methylation in orthotopic mouse pancreatic cancer model. KPC (cancer cells) and CAF (stromal cells) were orthotopically implanted in the pancreas of C57BL/6 mice and subsequently treated with LDH inhibitor GSK2837808A (6 mg/Kg/QD)) for 4 weeks. Treatment with LDH inhibitor GSK2837808A (6 mg/Kg/QD) decreased tumor weight (A) and tumor volume (B). TUNEL staining showed increase in apoptotic cells in LDH inhibitor treated group compared to control (C). Serum analysis showed decrease in L-2HG level in mice treated with LDH inhibitor compared to control (D). Treatment with LDH inhibitor significantly reduced H3K9 methylation in pancreatic tumors compared to control tumor (E). Tumors treated with LDH inhibitor also showed increase in PDX-1 staining (F) and decrease in CD133 staining (G) in pancreatic tumor section as shown by immunohistochemistry. * indicate p-value </= 0.05

### Inhibition of LDH modulated the immune microenvironment and sensitized to immune checkpoint inhibitors

Pancreatic tumors have a notoriously immunosuppressive microenvironment. Particularly, the dearth of CD8+ cytotoxic T-cells in the microenvironment is very apparent(Rao et al., 2018; Sharma et al., 2020). Flow cytometry analysis of the pancreatic tumors treated with LDH inhibitor showed an increased infiltration of CD8+ T-cells (**Figure 6A**). It is reported that the acidic microenvironment in a hypoxic tumor microenvironment can promote immune evasion by excluding CD8+T-cells(Erra Diaz et al., 2018). However, whether hypoxia mediated accumulation of L-2HG also played a role in this CD8+ T-cell exclusion is not known. Interestingly, tumor regressive effect of the LDH inhibitor was lost when pancreatic tumors were implanted orthotopically in the pancreas of CD8-KO mice (**Figure 6B, C**), indicating CD8 cells plays crucial role in tumor progression. To study if accumulated L-2HG as a result of hypoxia and increased LDH activity affected T-cell function, we next investigated the role of L-2HG in T cell migration and expansion using transwell migration and CFSE dilution assays, respectively. Our results showed that octyl- L- 2HG treatment decreased T cell migration compared to control (**Figure 6D**) and abolished their proliferation *in vitro* (**Figure 6E**). This corelated with our *in vivo* work which have shown that L-2HG inhibition results in increased tumor CD8 T cell infiltration and indicate that presence of L2-HG may prevent the expansion of these cells within the tumor. Further analysis also showed that splenocytes, CD4 and CD8 T cell populations are equally affected by L-2HG (**Figure 6E**).

**Figure 6:**
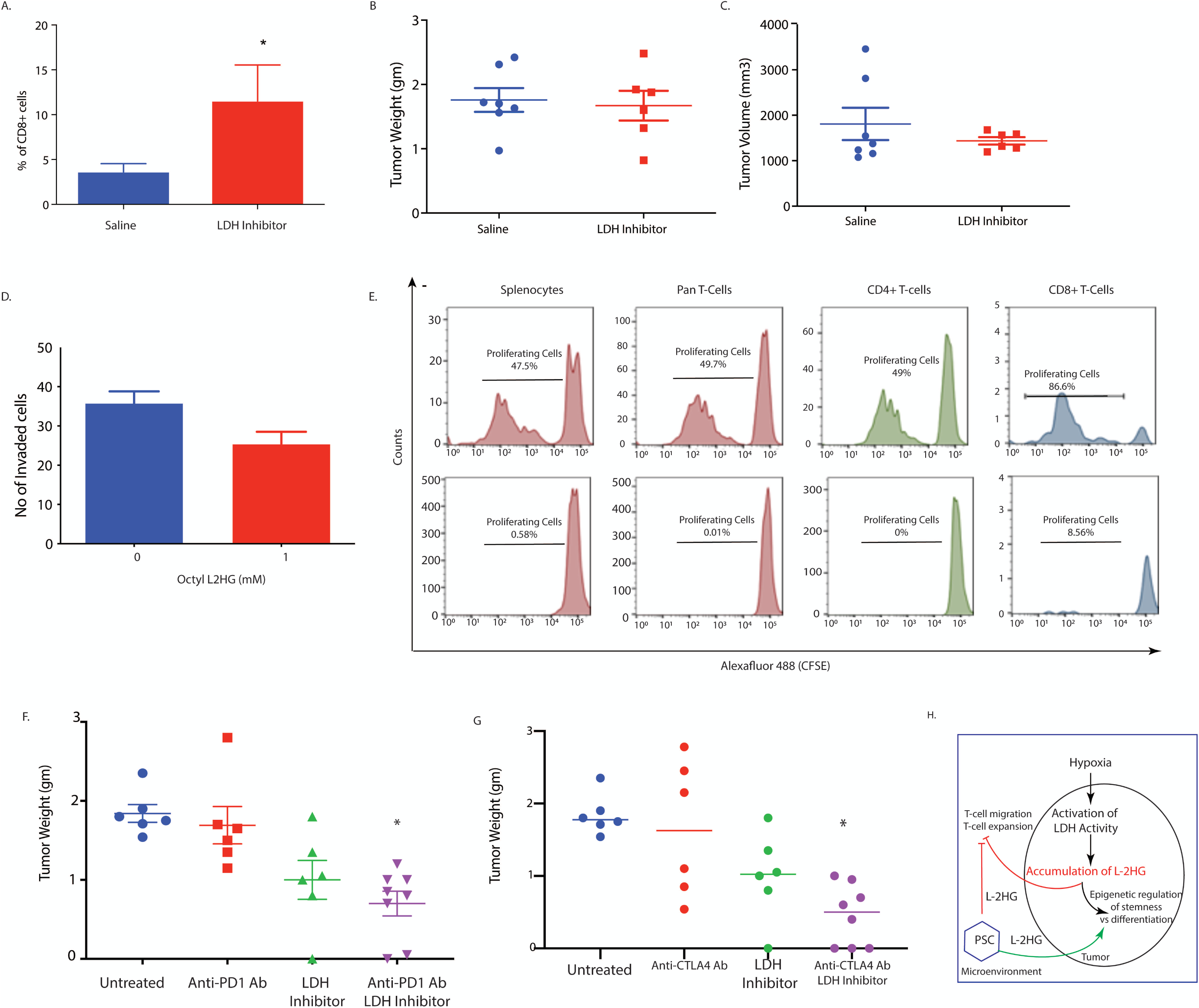
LDH inhibition induce anti-tumor immunity and sensitizes tumors to immunotherapy. In vivo LDH inhibition promote intra-tumoral CD8+ T cell infiltration as shown by flow cytometry (A). Effect of DON on tumor and stroma in syngeneic model was CD8 mediated as pancreatic tumors implanted in CD8KO mice did not show. LDH inhibitor mediated tumor regression was lost in the absence of CD8 T cells as orthotopically implanted pancreatic tumors in CD8 KO mice did not show any change in tumor weight (B) and volume (C) upon in vivo LDH inhibition. Octyl L-2HG treatment reduces T cell migration as shown by transwell migration assay (D). CFSE dilution assay showed significant reduction in proliferation in splenocytes and T cells when in vitro treated with octyl L-2HG (E). Treatment with LDH inhibitor sensitizes pancreatic tumors to anti PD-1 (F) and anti CTLA-4 (G) immune checkpoint therapy in orthotopic syngeneic mouse model of pancreatic cancer. Schematic figure showing how hypoxia driven L-2HG accumulation regulate self-renewal and tumor growth in pancreatic cancer (H). * indicate p-value </= 0.05. NS= Not significant.

Since increased CD8+ cells in the tumor microenvironment typically show enhanced sensitivity to checkpoint inhibitor therapy, we next treated orthotopically implanted pancreatic tumors (KPC:CAF in a ratio of 1:9) of C57/Bl6 mice and performed a combination treatment with LDH inhibitor and anti-PD1 antibody and LDH inhibitor with anti-CTLA4 antibody. Our results showed that effect of both checkpoint inhibitors was enhanced in the presence of LDH inhibition (**Figure 6F, G**).

## Discussion

Oncometabolites like 2HG has been implicated in epigenetic regulation of transcriptional machinery in AML and glioma. Produced by mutant IDH1/2 genes in these cancers, accumulated D-2HG inhibits the activity of enzymes like DNA and histone demethylases that are dependent on α−KG. This leads to a hypermethylated state of DNA and histones that can affect the transcriptional regulation of genes. 2HG exists as two enantiomers D- and L. While the D-2HG is produced as a consequence of IDH1 and IDH2 mutations in AML and glioma, recent studies have shown that low oxygen concentration and acidic microenvironment can lead to accumulation of L-2HG. Our current study showed that in pancreatic tumors that lacked IDH1/2 mutation, there was increased L-2HG accumulation, while there was almost no D-2HG in these samples (Figure 1). Further, we observed that majority of L-2HG in pancreatic tumors was being produced by the pancreatic stellate cells (PSC).

It is well known that microenvironmental component like hypoxia drive “stemness” in pancreatic tumors. We observed that as under hypoxia, treatment with cell permeable octyl L-2HG decreased expression of differentiation genes like PDX1 and increased expression of “stemness” genes like Sox2 and CD133 (Figure 2) in pancreatic cancer cells. Inhibition of LDH/MDH that were responsible for accumulation of L-2HG led to reversal of this observation.

Our study also indicated that the PSC secreted L-2HG acted in a paracrine manner to affect the “stemness-differentiation balance in the pancreatic cancer cells (Figure 2). To study how hypoxia mediated L-2HG affected the global proteome of the pancreatic cancer cells, we performed proteomic analysis of pancreatic cancer cells under hypoxia and after treatment with octyl L-2HG using Isobaric Tags for Relative and Absolute Quantitation or ITRAQ. Interestingly, as observed by our experimental observation, a significant number of proteins involved in RNA processing and chromatin remodeling were observed to be deregulated (Figure 3).

Our study revealed that under hypoxia, activated PSCs produced L-2HG which in turn was used by the tumor epithelial cells to reprogram their epigenetic machinery to regulate the balance between differentiation and de-differentiation or stemness. This is the first report of L-2HG mediated regulation of epigenetic programming in pancreatic cancer. To demonstrate this phenomenon, we focused on two proteins: CD133, that has been associated with all “stemness” properties in pancreatic cancer including tumor initiation, resistance to therapy, increased metastasis and increased self-renewal markers (Banerjee et al., 2014; Bao et al., 2014; Nomura et al., 2015); and Pdx1, a well-known gene that regulates differentiation in pancreatic cancer. Our results showed that H3K9 was mono-methylated in PDX1 under normal oxygen concentration but became trimethylated under hypoxia (Figure 3B). In case of CD133, that was mono-methylated under hypoxia, became trimethylated under normoxia (Figure 4C). Mono-methylation of H3K9 is an indicator of gene expression, while trimethylation of H3K9 is associated with inactivity or repression. This data showed that under normoxia, PDX1 expression was “on” while CD133 expression was baseline, while under hypoxia expression was turned “on” while PDX1 became inactive. Further, we determined that this was indeed due to L-2HG accumulation under hypoxia, since similar observation was made upon treatment with octyl L-2HG, that showed a global hyper-methylation of H3K4, H3K9 and H3K27 (Figure 4D) CD133 gene activity and PDX1 repression compared to untreated (Figure 4E, F). As a proof of principle, we silenced LDH and MDH genes that are responsible for L-2HG synthesis. Our studied confirmed that in the absence of LDH/MDH, that depleted the L-2HG pool under hypoxia, this effect was abrogated.

Since LDH appeared to be the major enzyme that was responsible for synthesis and accumulation of 2HG, we next used a commercial small molecule inhibitor of LDH GSK2837808A to study if inhibition of L-2HG accumulation *in vivo* would affect tumor growth by upsetting the stemness/differentiation balance. Our study showed that treatment with LDH inhibitor decreased tumor volume and induced apoptosis in tumors. Further, treatment with LDH inhibitor also reduced the L-2HG in the serum of the treated animals and decreased global histone modification. Additionally, the expression of CD133 and other stemness genes were downregulated in the treated samples, indicating that LDH inhibitors can be developed as a therapeutic strategy to overcome hypoxia mediated increased self-renewal and tumor recurrence in pancreatic tumors (Figure 5). Interestingly, effect of LDH inhibition was observed to be mediated by CD8, since in a CD8 knock-out mouse we observe a distinct insensitivity of the tumor to LDH inhibition (Figure 6A-C). This also indicated that apart from epigenetic reprogramming of the tumor cells, accumulated 2HG in the hypoxic pancreatic tumor was also affecting its immune microenvironment. To determine this, we assessed the migration and proliferation of T-cells in the presence of octyl L-2HG. Our results revealed that L-2HG prevented infiltration of T-cells in the tumor and inhibited their expansion (Figure 6E, F). This property of the oncometabolite contributed to immune suppression in pancreatic tumors and made them resistant to immune therapy. Upon inhibition of LDH, that also decreased L-2HG, pancreatic tumors could be sensitized to checkpoint inhibitor therapy like anti-PD1 and anti-CTLA4 antibody (Figure 6 F, G). PSCs have been implicated in modulating the microenvironment in pancreatic cancer and thus contribute to their immune evasion(Fu et al., 2018; Martinez-Bosch et al., 2018). However, this is the first study to show that PSC secreted oncometabolite can both remodel the epigenetic machinery in the tumor cells as well as contribute to the immune modulation in pancreatic cancer. The schematic **Figure 6H** demonstrates the summary of our findings.

Role of oncometabolites have not been studied in pancreatic cancer gene regulation. Our study is the first to show that under hypoxia, pancreatic stellate cells (as well as tumor epithelial cells) produce L-2HG. This alters histone methylation status at the promoter region of “stemness” and differentiation genes. In glioma and AML, role of D-2HG (as a result of IDH1/2 mutations) has been extensively studied particularly in context of epigenetic regulation (Dang et al., 2009; Nowicki and Gottlieb, 2015). Recent studies have implicated an immune modulatory role of 2HG in glioma(Bunse et al., 2018). However, they have not been studied in pancreatic cancer. Our observation that inhibition of L-2HG accumulation in pancreatic tumors can alter its immune microenvironment and make it amenable to immune therapy is extremely relevant in context of development of novel therapy regimen. While recent advances have shown that hypoxia and acidic pH can result in accumulation of the previously ignored L2-HG enantiomer, its role in cancers lacking IDH1 mutation has not been explored. In this context, our study on this pancreatic cancer oncometabolite sets the cornerstone that will lead to new mechanistic understanding of 2HG metabolism in the biology of tumor progression.

## Acknowledgement

The authors would like to thank Dr. Lluis Morey, University of Miami, FL for his extensive review and helpful criticism on this manuscript. The authors would also like to acknowledge the Flow Cytometry Shared Resources at the Sylvester Comprehensive Cancer Center for their help.

## References

Aghili, M., Zahedi, F., and Rafiee, E. (2009). Hydroxyglutaric aciduria and malignant brain tumor: a case report and literature review. J Neurooncol 91, 233–236.

Aponte, P.M., and Caicedo, A. (2017). Stemness in Cancer: Stem Cells, Cancer Stem Cells, and Their Microenvironment. Stem Cells Int 2017, 5619472.

Baird, T.D., and Wek, R.C. (2012). Eukaryotic initiation factor 2 phosphorylation and translational control in metabolism. Adv Nutr 3, 307–321.

Banerjee, S., Nomura, A., Sangwan, V., Chugh, R., Dudeja, V., Vickers, S.M., and Saluja, A. (2014). CD133+ tumor initiating cells in a syngenic murine model of pancreatic cancer respond to Minnelide. Clin Cancer Res 20, 2388–2399.

Bao, B., Azmi, A.S., Aboukameel, A., Ahmad, A., Bolling-Fischer, A., Sethi, S., Ali, S., Li, Y., Kong, D., Banerjee, S., et al. (2014). Pancreatic cancer stem-like cells display aggressive behavior mediated via activation of FoxQ1. J Biol Chem 289, 14520–14533.

Boise, L.H., and Shanmugam, M. (2019). Stromal Support of Metabolic Function through Mitochondrial Transfer in Multiple Myeloma. Cancer Res 79, 2102–2103.

Bunse, L., Pusch, S., Bunse, T., Sahm, F., Sanghvi, K., Friedrich, M., Alansary, D., Sonner, J.K., Green, E., Deumelandt, K., et al. (2018). Suppression of antitumor T cell immunity by the oncometabolite (R)-2-hydroxyglutarate. Nat Med 24, 1192–1203.

Chowdhury, R., Yeoh, K.K., Tian, Y.M., Hillringhaus, L., Bagg, E.A., Rose, N.R., Leung, I.K., Li, X.S., Woon, E.C., Yang, M., et al. (2011). The oncometabolite 2-hydroxyglutarate inhibits histone lysine demethylases. EMBO Rep 12, 463–469.

Dang, L., White, D.W., Gross, S., Bennett, B.D., Bittinger, M.A., Driggers, E.M., Fantin, V.R., Jang, H.G., Jin, S., Keenan, M.C., et al. (2009). Cancer-associated IDH1 mutations produce 2-hydroxyglutarate. Nature 462, 739–744.

Das, S., Shapiro, B., Vucic, E.A., Vogt, S., and Bar-Sagi, D. (2020). Tumor Cell-Derived IL1beta Promotes Desmoplasia and Immune Suppression in Pancreatic Cancer. Cancer Res 80, 1088–1101.

Dauer, P., Nomura, A., Saluja, A., and Banerjee, S. (2017). Microenvironment in determining chemo-resistance in pancreatic cancer: Neighborhood matters. Pancreatology 17, 7–12.

Erra Diaz, F., Dantas, E., and Geffner, J. (2018). Unravelling the Interplay between Extracellular Acidosis and Immune Cells. Mediators Inflamm 2018, 1218297.

Feig, C., Gopinathan, A., Neesse, A., Chan, D.S., Cook, N., and Tuveson, D.A. (2012). The pancreas cancer microenvironment. Clin Cancer Res 18, 4266–4276.

Figueroa, M.E., Abdel-Wahab, O., Lu, C., Ward, P.S., Patel, J., Shih, A., Li, Y., Bhagwat, N., Vasanthakumar, A., Fernandez, H.F., et al. (2010). Leukemic IDH1 and IDH2 mutations result in a hypermethylation phenotype, disrupt TET2 function, and impair hematopoietic differentiation. Cancer Cell 18, 553–567.

Fu, Y., Liu, S., Zeng, S., and Shen, H. (2018). The critical roles of activated stellate cells-mediated paracrine signaling, metabolism and onco-immunology in pancreatic ductal adenocarcinoma. Mol Cancer 17, 62.

Gasser, S., Lim, L.H.K., and Cheung, F.S.G. (2017). The role of the tumour microenvironment in immunotherapy. Endocr Relat Cancer 24, T283–T295.

Hanahan, D., and Weinberg, R.A. (2011). Hallmarks of cancer: the next generation. Cell 144, 646–674.

Ho, W.J., and Jaffee, E.M. (2020). Disrupting a converging metabolic target turns up the immunologic-heat in pancreatic tumors. J Clin Invest 130, 71–73.

Intlekofer, A.M., Dematteo, R.G., Venneti, S., Finley, L.W., Lu, C., Judkins, A.R., Rustenburg, A.S., Grinaway, P.B., Chodera, J.D., Cross, J.R., et al. (2015). Hypoxia Induces Production of L-2-Hydroxyglutarate. Cell Metab 22, 304–311.

Intlekofer, A.M., Wang, B., Liu, H., Shah, H., Carmona-Fontaine, C., Rustenburg, A.S., Salah, S., Gunner, M.R., Chodera, J.D., Cross, J.R., et al. (2017). L-2-Hydroxyglutarate production arises from noncanonical enzyme function at acidic pH. Nat Chem Biol 13, 494–500.

Janke, R., Dodson, A.E., and Rine, J. (2015). Metabolism and epigenetics. Annu Rev Cell Dev Biol 31, 473–496.

Kats, L.M., Reschke, M., Taulli, R., Pozdnyakova, O., Burgess, K., Bhargava, P., Straley, K., Karnik, R., Meissner, A., Small, D., et al. (2014). Proto-oncogenic role of mutant IDH2 in leukemia initiation and maintenance. Cell Stem Cell 14, 329–341.

Kohanbash, G., Carrera, D.A., Shrivastav, S., Ahn, B.J., Jahan, N., Mazor, T., Chheda, Z.S., Downey, K.M., Watchmaker, P.B., Beppler, C., et al. (2017). Isocitrate dehydrogenase mutations suppress STAT1 and CD8+ T cell accumulation in gliomas. J Clin Invest 127, 1425–1437.

Linster, C.L., Van Schaftingen, E., and Hanson, A.D. (2013). Metabolite damage and its repair or pre-emption. Nat Chem Biol 9, 72–80.

Luo, W., and Wang, Y. (2019). Hypoxia Mediates Tumor Malignancy and Therapy Resistance. Adv Exp Med Biol 1136, 1–18.

Martinez-Bosch, N., Vinaixa, J., and Navarro, P. (2018). Immune Evasion in Pancreatic Cancer: From Mechanisms to Therapy. Cancers (Basel) 10.

McGinn, O., Gupta, V.K., Dauer, P., Arora, N., Sharma, N., Nomura, A., Dudeja, V., Saluja, A., and Banerjee, S. (2017). Inhibition of hypoxic response decreases stemness and reduces tumorigenic signaling due to impaired assembly of HIF1 transcription complex in pancreatic cancer. Sci Rep 7, 7872.

Menendez, J.A., Corominas-Faja, B., Cuyas, E., Garcia, M.G., Fernandez-Arroyo, S., Fernandez, A.F., Joven, J., Fraga, M.F., and Alarcon, T. (2016). Oncometabolic Nuclear Reprogramming of Cancer Stemness. Stem Cell Reports 6, 273–283.

Nadtochiy, S.M., Schafer, X., Fu, D., Nehrke, K., Munger, J., and Brookes, P.S. (2016). Acidic pH Is a Metabolic Switch for 2-Hydroxyglutarate Generation and Signaling. J Biol Chem 291, 20188–20197.

Nomura, A., Banerjee, S., Chugh, R., Dudeja, V., Yamamoto, M., Vickers, S.M., and Saluja, A.K. (2015). CD133 initiates tumors, induces epithelial-mesenchymal transition and increases metastasis in pancreatic cancer. Oncotarget 6, 8313–8322.

Nomura, A., Dauer, P., Gupta, V., McGinn, O., Arora, N., Majumdar, K., Uhlrich, C., 3rd, Dalluge, J., Dudeja, V., Saluja, A., et al. (2016). Microenvironment mediated alterations to metabolic pathways confer increased chemo-resistance in CD133+ tumor initiating cells. Oncotarget 7, 56324–56337.

Nowicki, S., and Gottlieb, E. (2015). Oncometabolites: tailoring our genes. FEBS J 282, 2796–2805.

O’Brien, C.A., Kreso, A., and Jamieson, C.H. (2010). Cancer stem cells and self-renewal. Clin Cancer Res 16, 3113–3120.

Oldham, W.M., Clish, C.B., Yang, Y., and Loscalzo, J. (2015). Hypoxia-Mediated Increases in L-2-hydroxyglutarate Coordinate the Metabolic Response to Reductive Stress. Cell Metab 22, 291–303.

Qian, J., and Rankin, E.B. (2019). Hypoxia-Induced Phenotypes that Mediate Tumor Heterogeneity. Adv Exp Med Biol 1136, 43–55.

Rao, C.V., Mohammed, A., Asch, A.S., and Janakiram, N.B. (2018). Immunoprevention of Pancreatic Cancer. Curr Med Chem 25, 2576–2584.

Richardson, L.G., Choi, B.D., and Curry, W.T. (2019). (R)-2-hydroxyglutarate drives immune quiescence in the tumor microenvironment of IDH-mutant gliomas. Transl Cancer Res 8, S167–S170.

Sangwan, V., Banerjee, S., Jensen, K.M., Chen, Z., Chugh, R., Dudeja, V., Vickers, S.M., and Saluja, A.K. (2015). Primary and liver metastasis-derived cell lines from KrasG12D; Trp53R172H; Pdx-1 Cre animals undergo apoptosis in response to triptolide. Pancreas 44, 583–589.

Sciacovelli, M., and Frezza, C. (2016). Oncometabolites: Unconventional triggers of oncogenic signalling cascades. Free Radic Biol Med 100, 175–181.

Seto, E., and Yoshida, M. (2014). Erasers of histone acetylation: the histone deacetylase enzymes. Cold Spring Harb Perspect Biol 6, a018713.

Sharma, N.S., Gupta, V.K., Garrido, V.T., Hadad, R., Durden, B.C., Kesh, K., Giri, B., Ferrantella, A., Dudeja, V., Saluja, A., et al. (2020). Targeting tumor-intrinsic hexosamine biosynthesis sensitizes pancreatic cancer to anti-PD1 therapy. J Clin Invest 130, 451–465.

Sharon, Y., Alon, L., Glanz, S., Servais, C., and Erez, N. (2013). Isolation of normal and cancer-associated fibroblasts from fresh tissues by Fluorescence Activated Cell Sorting (FACS). J Vis Exp, e4425.

Siegel, R.L., Miller, K.D., and Jemal, A. (2019). Cancer statistics, 2019. CA Cancer J Clin 69, 7–34.

Simon, M.C., and Keith, B. (2008). The role of oxygen availability in embryonic development and stem cell function. Nat Rev Mol Cell Biol 9, 285–296.

Szklarczyk, D., Gable, A.L., Lyon, D., Junge, A., Wyder, S., Huerta-Cepas, J., Simonovic, M., Doncheva, N.T., Morris, J.H., Bork, P., et al. (2019). STRING v11: protein-protein association networks with increased coverage, supporting functional discovery in genome-wide experimental datasets. Nucleic Acids Res 47, D607–D613.

Turcan, S., Rohle, D., Goenka, A., Walsh, L.A., Fang, F., Yilmaz, E., Campos, C., Fabius, A.W., Lu, C., Ward, P.S., et al. (2012). IDH1 mutation is sufficient to establish the glioma hypermethylator phenotype. Nature 483, 479–483.

Tyrakis, P.A., Palazon, A., Macias, D., Lee, K.L., Phan, A.T., Velica, P., You, J., Chia, G.S., Sim, J., Doedens, A., et al. (2016). S-2-hydroxyglutarate regulates CD8(+) T-lymphocyte fate. Nature 540, 236–241.

Xu, W., Yang, H., Liu, Y., Yang, Y., Wang, P., Kim, S.H., Ito, S., Yang, C., Wang, P., Xiao, M.T., et al. (2011). Oncometabolite 2-hydroxyglutarate is a competitive inhibitor of alpha-ketoglutarate-dependent dioxygenases. Cancer Cell 19, 17–30.

Zhang, T., Somasundaram, R., Berencsi, K., Caputo, L., Gimotty, P., Rani, P., Guerry, D., Swoboda, R., and Herlyn, D. (2006). Migration of cytotoxic T lymphocytes toward melanoma cells in three-dimensional organotypic culture is dependent on CCL2 and CCR4. Eur J Immunol 36, 457–467.

Zhang, Y., Lazarus, J., Steele, N.G., Yan, W., Lee, H.J., Nwosu, Z.C., Halbrook, C.J., Menjivar, R.E., Kemp, S.B., Sirihorachai, V.R., et al. (2020). Regulatory T-cell Depletion Alters the Tumor Microenvironment and Accelerates Pancreatic Carcinogenesis. Cancer Discov 10, 422–439.

