## Supplementary Figures for "Hypoxia-driven oncometabolite L-2HG maintains “stemness”-differentiation balance and facilitates immune suppression in pancreatic cancer"

| Checked | Master | Accession |
|---|---|---|
| TRUE | Master Protein | V9HWE1 |
| TRUE | Master Protein | P63261 |
| TRUE | Master Protein | Q6FI13 |
| TRUE | Master Protein | O60814 |
| TRUE | Master Protein | P10809 |
| TRUE | Master Protein | P06899 |
| TRUE | Master Protein | Q16695 |
| TRUE | Master Protein | P07437 |
| TRUE | Master Protein | P68371 |
| TRUE | Master Protein | B2R6L0 |
| TRUE | Master Protein | Q5TEC6 |
| TRUE | Master Protein | P07900-2 |
| TRUE | Master Protein | A0A0S2Z491 |
| TRUE | Master Protein | P08238 |
| TRUE | Master Protein | P61604 |
| TRUE | Master Protein | P0CG39 |
| TRUE | Master Protein | M1VPF4 |
| TRUE | Master Protein | P06733 |
| TRUE | Master Protein | A0A024R5Z9 |
| TRUE | Master Protein | B7Z1K5 |
| TRUE | Master Protein | A8K486 |
| TRUE | Master Protein | X5D2T3 |
| TRUE | Master Protein | Q00839 |
| TRUE | Master Protein | P19338 |
| TRUE | Master Protein | B5BU38 |
| TRUE | Master Protein | A0A024QYX3 |
| TRUE | Master Protein | P05787-2 |
| TRUE | Master Protein | P26373 |
| TRUE | Master Protein | A0A024R4K3 |
| TRUE | Master Protein | V9HWB4 |
| TRUE | Master Protein | Q8N1C8 |
| TRUE | Master Protein | B2RA03 |
| TRUE | Master Protein | E5RHG6 |
| TRUE | Master Protein | P30101 |
| TRUE | Master Protein | P23246 |
| TRUE | Master Protein | P16070 |
| TRUE | Master Protein | A0A1W2PQS6 |
| TRUE | Master Protein | B2R4P2 |
| TRUE | Master Protein | P13639 |
| TRUE | Master Protein | P04406 |
| TRUE | Master Protein | Q53F64 |
| TRUE | Master Protein | A8K088 |
| TRUE | Master Protein | P04075-2 |
| TRUE | Master Protein | A0A023T6R1 |
| TRUE | Master Protein | A0A087X0X3 |
| TRUE | Master Protein | Q6IPH7 |
| TRUE | Master Protein | V9HW43 |

| | | |
|---|---|---|
| TRUE | Master Protein | Q13344 |
| TRUE | Master Protein | B2R4R9 |
| TRUE | Master Protein | E9PK25 |
| TRUE | Master Protein | A0A024RBF6 |
| TRUE | Master Protein | H0Y449 |
| TRUE | Master Protein | A0A0A0MTS2 |
| TRUE | Master Protein | Q0D2Q6 |
| TRUE | Master Protein | Q15181 |
| TRUE | Master Protein | Q59G24 |
| TRUE | Master Protein | B2R6J2 |
| TRUE | Master Protein | P07355-2 |
| TRUE | Master Protein | P60174 |
| TRUE | Master Protein | P08195-4 |
| TRUE | Master Protein | P62316 |
| TRUE | Master Protein | Q14978-2 |
| TRUE | Master Protein | Q9BVJ6 |
| TRUE | Master Protein | Q32Q12 |
| TRUE | Master Protein | G3V2S9 |
| TRUE | Master Protein | P23381 |
| TRUE | Master Protein | E7EX29 |
| TRUE | Master Protein | P78371 |
| TRUE | Master Protein | A0A024RBE7 |
| TRUE | Master Protein | O75367 |
| TRUE | Master Protein | Q09666 |
| TRUE | Master Protein | P09382 |
| TRUE | Master Protein | Q07955-2 |
| TRUE | Master Protein | O75369-8 |
| TRUE | Master Protein | M0R0R2 |
| TRUE | Master Protein | B2RDQ3 |
| TRUE | Master Protein | Q9NY61 |
| TRUE | Master Protein | B2RD27 |
| TRUE | Master Protein | B2R8R5 |
| TRUE | Master Protein | B4DE59 |
| TRUE | Master Protein | Q15084-2 |
| TRUE | Master Protein | A0A024R9P1 |
| TRUE | Master Protein | Q9BV57 |
| TRUE | Master Protein | P16401 |
| TRUE | Master Protein | E9PRY8 |
| TRUE | Master Protein | B7Z268 |
| TRUE | Master Protein | P13804 |
| TRUE | Master Protein | B9EG90 |
| TRUE | Master Protein | Q15424-3 |
| TRUE | Master Protein | Q59F66 |
| TRUE | Master Protein | H7C0C1 |
| TRUE | Master Protein | A8K9K6 |
| TRUE | Master Protein | A0A0B4J2C3 |
| TRUE | Master Protein | P26641-2 |
| TRUE | Master Protein | O60664 |

| | | |
|---|---|---|
| TRUE | Master Protein | Q59FR8 |
| TRUE | Master Protein | Q5JR94 |
| TRUE | Master Protein | P49773 |
| TRUE | Master Protein | Q59GM9 |
| TRUE | Master Protein | Q9H6R4 |
| TRUE | Master Protein | G8JLB6 |
| TRUE | Master Protein | Q96CT7 |
| TRUE | Master Protein | B7Z4C8 |
| TRUE | Master Protein | A8MXP9 |
| TRUE | Master Protein | P27824-2 |
| TRUE | Master Protein | B7Z6Z4 |
| TRUE | Master Protein | V9HW80 |
| TRUE | Master Protein | A4D198 |
| TRUE | Master Protein | E5KNY5 |
| TRUE | Master Protein | O75390 |
| TRUE | Master Protein | Q9NUJ1 |
| TRUE | Master Protein | A6NIZ1 |
| TRUE | Master Protein | B4DYN5 |
| TRUE | Master Protein | Q5TZA2 |
| TRUE | Master Protein | P06703 |
| TRUE | Master Protein | P31930 |
| TRUE | Master Protein | P00492 |
| TRUE | Master Protein | P00558 |
| TRUE | Master Protein | O43852-3 |
| TRUE | Master Protein | A8K401 |
| TRUE | Master Protein | B2R7B5 |
| TRUE | Master Protein | B4DF19 |
| TRUE | Master Protein | Q7Z7F4 |
| TRUE | Master Protein | P27144 |
| TRUE | Master Protein | Q92994-4 |
| TRUE | Master Protein | A0A126LAY7 |
| TRUE | Master Protein | A0A075B6Z2 |
| TRUE | Master Protein | A0A024R8B6 |
| TRUE | Master Protein | Q17R60 |
| TRUE | Master Protein | Q658U4 |
| TRUE | Master Protein | A8K7D9 |
| TRUE | Master Protein | P61247 |
| TRUE | Master Protein | A0A218KGR2 |
| TRUE | Master Protein | P78527 |
| TRUE | Master Protein | J9R021 |
| TRUE | Master Protein | Q3MIX3 |
| TRUE | Master Protein | Q6IPI1 |
| TRUE | Master Protein | Q9ULV4-3 |
| TRUE | Master Protein | A0A024R5Q1 |
| TRUE | Master Protein | O60296 |
| TRUE | Master Protein | H0YHV6 |

| Description |
|---|
| Epididymis luminal protein 113 OS=Homo sapiens GN=HEL113 PE=2 SV=1 |
| Actin, cytoplasmic 2 OS=Homo sapiens GN=ACTG1 PE=1 SV=1 |
| Histone H2A type 2-A OS=Homo sapiens GN=HIST2H2AA3 PE=1 SV=3 |
| Histone H2B type 1-K OS=Homo sapiens GN=HIST1H2BK PE=1 SV=3 |
| 60 kDa heat shock protein, mitochondrial OS=Homo sapiens GN=HSPD1 PE=1 SV=2 |
| Histone H2B type 1-J OS=Homo sapiens GN=HIST1H2BJ PE=1 SV=3 |
| Histone H3.1t OS=Homo sapiens GN=HIST3H3 PE=1 SV=3 |
| Tubulin beta chain OS=Homo sapiens GN=TUBB PE=1 SV=2 |
| Tubulin beta-4B chain OS=Homo sapiens GN=TUBB4B PE=1 SV=1 |
| Tubulin beta chain OS=Homo sapiens PE=2 SV=1 |
| Histone H3 OS=Homo sapiens GN=HIST2H3PS2 PE=1 SV=1 |
| Isoform 2 of Heat shock protein HSP 90-alpha OS=Homo sapiens GN=HSP90AA1 |
| Nucleophosmin isoform 2 (Fragment) OS=Homo sapiens GN=NPM1 PE=2 SV=1 |
| Heat shock protein HSP 90-beta OS=Homo sapiens GN=HSP90AB1 PE=1 SV=4 |
| 10 kDa heat shock protein, mitochondrial OS=Homo sapiens GN=HSPE1 PE=1 SV=2 |
| POTE ankyrin domain family member J OS=Homo sapiens GN=POTEJ PE=3 SV=1 |
| Tyrosine-protein kinase receptor OS=Homo sapiens GN=TPM3-ROS1 PE=2 SV=1 |
| Alpha-enolase OS=Homo sapiens GN=ENO1 PE=1 SV=2 |
| Pyruvate kinase OS=Homo sapiens GN=PKM2 PE=3 SV=1 |
| Tubulin alpha chain OS=Homo sapiens PE=2 SV=1 |
| Peptidyl-prolyl cis-trans isomerase OS=Homo sapiens PE=2 SV=1 |
| Ribosomal protein L10 isoform A (Fragment) OS=Homo sapiens GN=RPL10 PE=2 SV=1 |
| Heterogeneous nuclear ribonucleoprotein U OS=Homo sapiens GN=HNRNPU PE=1 SV=6 |
| Nucleolin OS=Homo sapiens GN=NCL PE=1 SV=3 |
| Annexin OS=Homo sapiens GN=ANXA1 PE=2 SV=1 |
| RNA binding motif (RNP1, RRM) protein 3, isoform CRA_c OS=Homo sapiens GN=RBM3 PE=4 SV=1 |
| Isoform 2 of Keratin, type II cytoskeletal 8 OS=Homo sapiens GN=KRT8 |
| 60S ribosomal protein L13 OS=Homo sapiens GN=RPL13 PE=1 SV=4 |
| Malate dehydrogenase OS=Homo sapiens GN=MDH2 PE=2 SV=1 |
| Epididymis secretory sperm binding protein Li 89n OS=Homo sapiens GN=HEL-S-89n PE=2 SV=1 |
| HSPA9 protein (Fragment) OS=Homo sapiens GN=HSPA9 PE=2 SV=1 |
| cDNA, FLJ94640, highly similar to Homo sapiens keratin 18 (KRT18), mRNA OS=Homo sapiens PE=2 SV=1 |
| Tubulin-specific chaperone A OS=Homo sapiens GN=TBCA PE=1 SV=2 |
| Protein disulfide-isomerase A3 OS=Homo sapiens GN=PDIA3 PE=1 SV=4 |
| Splicing factor, proline- and glutamine-rich OS=Homo sapiens GN=SFPQ PE=1 SV=2 |
| CD44 antigen OS=Homo sapiens GN=CD44 PE=1 SV=3 |
| RPS10-NUDT3 readthrough OS=Homo sapiens GN=RPS10-NUDT3 PE=4 SV=1 |
| cDNA, FLJ92164, highly similar to Homo sapiens peroxiredoxin 1 (PRDX1), mRNA OS=Homo sapiens PE=2 SV=1 |
| Elongation factor 2 OS=Homo sapiens GN=EEF2 PE=1 SV=4 |
| Glyceraldehyde-3-phosphate dehydrogenase OS=Homo sapiens GN=GAPDH PE=1 SV=3 |
| Heterogeneous nuclear ribonucleoprotein AB isoform a variant (Fragment) OS=Homo sapiens PE=2 SV=1 |
| cDNA FLJ78614, highly similar to Homo sapiens eukaryotic translation initiation factor 4A, isoform 1 (EIF4A1), mRNA OS |
| Isoform 2 of Fructose-bisphosphate aldolase A OS=Homo sapiens GN=ALDOA |
| Mago nashi protein OS=Homo sapiens GN=FLJ10292 PE=2 SV=1 |
| Heterogeneous nuclear ribonucleoprotein M OS=Homo sapiens GN=HNRNPM PE=1 SV=1 |
| RPL14 protein OS=Homo sapiens GN=RPL14 PE=1 SV=1 |
| Epididymis secretory protein Li 102 OS=Homo sapiens GN=HEL-S-102 PE=2 SV=1 |

| |
|---|
| Fus-like protein (Fragment) OS=Homo sapiens PE=2 SV=1 |
| HCG26477 OS=Homo sapiens GN=RPS28 PE=2 SV=1 |
| Cofilin-1 OS=Homo sapiens GN=CFL1 PE=1 SV=1 |
| HCG26523, isoform CRA_a OS=Homo sapiens GN=hCG_26523 PE=4 SV=1 |
| Nuclease-sensitive element-binding protein 1 (Fragment) OS=Homo sapiens GN=YBX1 PE=1 SV=1 |
| Glucose-6-phosphate isomerase (Fragment) OS=Homo sapiens GN=GPI PE=1 SV=1 |
| Phosphoglycerate mutase OS=Homo sapiens GN=PGAM1 PE=2 SV=1 |
| Inorganic pyrophosphatase OS=Homo sapiens GN=PPA1 PE=1 SV=2 |
| Activated RNA polymerase II transcription cofactor 4 variant (Fragment) OS=Homo sapiens PE=2 SV=1 |
| cDNA, FLJ92973, highly similar to Homo sapiens villin 2 (ezrin) (VIL2), mRNA OS=Homo sapiens PE=2 SV=1 |
| Isoform 2 of Annexin A2 OS=Homo sapiens GN=ANXA2 |
| Triosephosphate isomerase OS=Homo sapiens GN=TPI1 PE=1 SV=3 |
| Isoform 4 of 4F2 cell-surface antigen heavy chain OS=Homo sapiens GN=SLC3A2 |
| Small nuclear ribonucleoprotein Sm D2 OS=Homo sapiens GN=SNRPD2 PE=1 SV=1 |
| Isoform Beta of Nucleolar and coiled-body phosphoprotein 1 OS=Homo sapiens GN=NOLC1 |
| U3 small nucleolar RNA-associated protein 14 homolog A OS=Homo sapiens GN=UTP14A PE=1 SV=1 |
| Nucleoside diphosphate kinase OS=Homo sapiens GN=NME1-NME2 PE=1 SV=1 |
| SRA stem-loop-interacting RNA-binding protein, mitochondrial OS=Homo sapiens GN=SLIRP PE=1 SV=1 |
| Tryptophan--tRNA ligase, cytoplasmic OS=Homo sapiens GN=WARS PE=1 SV=2 |
| 14-3-3 protein zeta/delta (Fragment) OS=Homo sapiens GN=YWHAZ PE=1 SV=1 |
| T-complex protein 1 subunit beta OS=Homo sapiens GN=CCT2 PE=1 SV=4 |
| Thymopoietin, isoform CRA_c OS=Homo sapiens GN=TMPO PE=4 SV=1 |
| Core histone macro-H2A.1 OS=Homo sapiens GN=H2AFY PE=1 SV=4 |
| Neuroblast differentiation-associated protein AHNAK OS=Homo sapiens GN=AHNAK PE=1 SV=2 |
| Galectin-1 OS=Homo sapiens GN=LGALS1 PE=1 SV=2 |
| Isoform ASF-2 of Serine/arginine-rich splicing factor 1 OS=Homo sapiens GN=SRSF1 |
| Isoform 8 of Filamin-B OS=Homo sapiens GN=FLNB |
| 40S ribosomal protein S5 OS=Homo sapiens GN=RPS5 PE=1 SV=1 |
| cDNA, FLJ96718, highly similar to Homo sapiens splicing factor, arginine/serine-rich 10 (transformer 2 homolog, Drosop |
| Protein AATF OS=Homo sapiens GN=AATF PE=1 SV=1 |
| cDNA, FLJ96428, highly similar to Homo sapiens proteasome (prosome, macropain) 26S subunit, non-ATPase, 7 (Mov34 |
| cDNA, FLJ94025, highly similar to Homo sapiens tripartite motif-containing 28 (TRIM28), mRNA OS=Homo sapiens PE=2 |
| cDNA FLJ60424, highly similar to Junction plakoglobin OS=Homo sapiens PE=2 SV=1 |
| Isoform 2 of Protein disulfide-isomerase A6 OS=Homo sapiens GN=PDIA6 |
| HCG2026745, isoform CRA_a OS=Homo sapiens GN=hCG_2026745 PE=4 SV=1 |
| 1,2-dihydroxy-3-keto-5-methylthiopentene dioxygenase OS=Homo sapiens GN=ADI1 PE=1 SV=1 |
| Histone H1.5 OS=Homo sapiens GN=HIST1H1B PE=1 SV=3 |
| Elongation factor 1-delta OS=Homo sapiens GN=EEF1D PE=1 SV=1 |
| Single-stranded DNA binding protein 1, isoform CRA_c OS=Homo sapiens GN=SSBP1 PE=2 SV=1 |
| Electron transfer flavoprotein subunit alpha, mitochondrial OS=Homo sapiens GN=ETFA PE=1 SV=1 |
| Topoisomerase (DNA) I OS=Homo sapiens GN=TOP1 PE=2 SV=1 |
| Isoform 3 of Scaffold attachment factor B1 OS=Homo sapiens GN=SAFB |
| DEAD box polypeptide 17 isoform p82 variant (Fragment) OS=Homo sapiens PE=2 SV=1 |
| Uncharacterized protein (Fragment) OS=Homo sapiens PE=4 SV=2 |
| cDNA FLJ76962, highly similar to Homo sapiens nucleolar protein 5A (56kDa with KKE/D repeat) (NOL5A), mRNA OS=Hor |
| Translationally-controlled tumor protein OS=Homo sapiens GN=TPT1 PE=1 SV=1 |
| Isoform 2 of Elongation factor 1-gamma OS=Homo sapiens GN=EEF1G |
| Perilipin-3 OS=Homo sapiens GN=PLIN3 PE=1 SV=3 |

| |
|---|
| Galectin (Fragment) OS=Homo sapiens PE=2 SV=1 |
| 40S ribosomal protein S8 OS=Homo sapiens GN=RPS8 PE=2 SV=1 |
| Histidine triad nucleotide-binding protein 1 OS=Homo sapiens GN=HINT1 PE=1 SV=2 |
| Alpha-1,4 glucan phosphorylase (Fragment) OS=Homo sapiens PE=2 SV=1 |
| Nucleolar protein 6 OS=Homo sapiens GN=NOL6 PE=1 SV=2 |
| Heterogeneous nuclear ribonucleoprotein H OS=Homo sapiens GN=HNRNPH1 PE=1 SV=1 |
| Coiled-coil domain-containing protein 124 OS=Homo sapiens GN=CCDC124 PE=1 SV=1 |
| 60S ribosomal protein L31 OS=Homo sapiens GN=RPL31 PE=1 SV=1 |
| Matrin-3 OS=Homo sapiens GN=MATR3 PE=1 SV=1 |
| Isoform 2 of Calnexin OS=Homo sapiens GN=CANX |
| Myosin light polypeptide 6 OS=Homo sapiens GN=MYL6 PE=1 SV=1 |
| Epididymis luminal protein 220 OS=Homo sapiens GN=HEL-S-70 PE=1 SV=1 |
| Similar to mKIAA0038 protein OS=Homo sapiens GN=LOC392647 PE=4 SV=1 |
| Leucine-rich PPR-motif containing OS=Homo sapiens GN=LRPPRC PE=4 SV=1 |
| Citrate synthase, mitochondrial OS=Homo sapiens GN=CS PE=1 SV=2 |
| Mycophenolic acid acyl-glucuronide esterase, mitochondrial OS=Homo sapiens GN=ABHD10 PE=1 SV=1 |
| Ras-related protein Rap-1b-like protein OS=Homo sapiens PE=2 SV=1 |
| Succinate dehydrogenase [ubiquinone] flavoprotein subunit, mitochondrial OS=Homo sapiens PE=2 SV=1 |
| Rootletin OS=Homo sapiens GN=CROCC PE=1 SV=1 |
| Protein S100-A6 OS=Homo sapiens GN=S100A6 PE=1 SV=1 |
| Cytochrome b-c1 complex subunit 1, mitochondrial OS=Homo sapiens GN=UQCRC1 PE=1 SV=3 |
| Hypoxanthine-guanine phosphoribosyltransferase OS=Homo sapiens GN=HPRT1 PE=1 SV=2 |
| Phosphoglycerate kinase 1 OS=Homo sapiens GN=PGK1 PE=1 SV=3 |
| Isoform 3 of Calumenin OS=Homo sapiens GN=CALU |
| Prohibitin, isoform CRA_a OS=Homo sapiens GN=PHB PE=2 SV=1 |
| cDNA, FLJ93365, highly similar to Homo sapiens KH domain containing, RNA binding, signal transduction associated 1 ( |
| cDNA FLJ50099, highly similar to Endothelin-converting enzyme 2 (EC 3.4.24.71) OS=Homo sapiens PE=2 SV=1 |
| Prestin OS=Homo sapiens GN=PRES PE=2 SV=1 |
| Adenylate kinase 4, mitochondrial OS=Homo sapiens GN=AK4 PE=1 SV=1 |
| Isoform 4 of Transcription factor IIIB 90 kDa subunit OS=Homo sapiens GN=BRF1 |
| U40 OS=Homo sapiens GN=U40 PE=3 SV=1 |
| T-cell receptor alpha joining 56 (Fragment) OS=Homo sapiens GN=TRAJ56 PE=4 SV=1 |
| Nucleoporin 214kDa, isoform CRA_b OS=Homo sapiens GN=NUP214 PE=4 SV=1 |
| Interphotoreceptor matrix proteoglycan 1 OS=Homo sapiens GN=IMPG1 PE=1 SV=2 |
| Peptidylprolyl isomerase OS=Homo sapiens GN=DKFZp666D193 PE=2 SV=1 |
| Importin subunit alpha OS=Homo sapiens PE=2 SV=1 |
| 40S ribosomal protein S3a OS=Homo sapiens GN=RPS3A PE=1 SV=2 |
| Amyloid beta A4 protein isoform a OS=Homo sapiens GN=APP PE=4 SV=1 |
| DNA-dependent protein kinase catalytic subunit OS=Homo sapiens GN=PRKDC PE=1 SV=3 |
| Eukaryotic translation initiation factor 3 subunit A OS=Homo sapiens GN=eIF3a PE=2 SV=1 |
| Uncharacterized aarF domain-containing protein kinase 5 OS=Homo sapiens GN=ADCK5 PE=1 SV=2 |
| 60S ribosomal protein L29 OS=Homo sapiens GN=RPL29 PE=2 SV=1 |
| Isoform 3 of Coronin-1C OS=Homo sapiens GN=CORO1C |
| HCG2003792, isoform CRA_b OS=Homo sapiens GN=hCG_2003792 PE=4 SV=1 |
| Trafficking kinesin-binding protein 2 OS=Homo sapiens GN=TRAK2 PE=1 SV=2 |
| WD repeat-containing protein 90 (Fragment) OS=Homo sapiens GN=WDR90 PE=4 SV=1 |

| Coverage | # Peptides | # PSMs | # Unique Peptides | # Protein Groups | # AAs |
|---|---|---|---|---|---|
| 65.2360515 | 29 | 189 | 8 | 1 | 466 |
| 39.2 | 16 | 118 | 3 | 1 | 375 |
| 61.53846154 | 10 | 60 | 5 | 1 | 130 |
| 74.6031746 | 15 | 94 | 0 | 1 | 126 |
| 27.39965096 | 17 | 51 | 17 | 1 | 573 |
| 74.6031746 | 14 | 86 | 1 | 1 | 126 |
| 58.08823529 | 10 | 99 | 1 | 1 | 136 |
| 28.15315315 | 10 | 67 | 1 | 1 | 444 |
| 28.08988764 | 10 | 63 | 2 | 1 | 445 |
| 22.02247191 | 7 | 50 | 1 | 1 | 445 |
| 32.35294118 | 7 | 83 | 1 | 1 | 136 |
| 6.557377049 | 7 | 21 | 2 | 1 | 854 |
| 7.823129252 | 3 | 28 | 3 | 1 | 294 |
| 8.977900552 | 8 | 24 | 3 | 1 | 724 |
| 89.21568627 | 10 | 39 | 10 | 1 | 102 |
| 4.238921002 | 3 | 29 | 1 | 1 | 1038 |
| 6.896551724 | 4 | 19 | 4 | 1 | 725 |
| 27.64976959 | 10 | 23 | 10 | 1 | 434 |
| 17.70244821 | 5 | 12 | 5 | 1 | 531 |
| 11.75337187 | 4 | 10 | 4 | 1 | 519 |
| 35.75757576 | 6 | 19 | 6 | 1 | 165 |
| 10.28037383 | 3 | 15 | 3 | 1 | 214 |
| 6.666666667 | 5 | 15 | 2 | 1 | 825 |
| 12.25352113 | 5 | 9 | 5 | 1 | 710 |
| 20.23121387 | 5 | 20 | 5 | 1 | 346 |
| 21.01910828 | 2 | 10 | 2 | 1 | 157 |
| 8.806262231 | 4 | 11 | 4 | 1 | 511 |
| 7.582938389 | 2 | 5 | 2 | 1 | 211 |
| 16.86390533 | 3 | 5 | 3 | 1 | 338 |
| 8.868501529 | 3 | 8 | 3 | 1 | 654 |
| 6.461086637 | 4 | 12 | 4 | 1 | 681 |
| 8.139534884 | 3 | 8 | 2 | 1 | 430 |
| 8.396946565 | 1 | 5 | 1 | 1 | 131 |
| 10.69306931 | 3 | 5 | 3 | 1 | 505 |
| 3.96039604 | 3 | 7 | 1 | 1 | 707 |
| 1.617250674 | 1 | 10 | 1 | 1 | 742 |
| 5.498281787 | 2 | 4 | 2 | 1 | 291 |
| 19.59798995 | 4 | 8 | 4 | 1 | 199 |
| 5.827505828 | 4 | 8 | 4 | 1 | 858 |
| 19.40298507 | 5 | 15 | 5 | 1 | 335 |
| 15.36144578 | 4 | 6 | 4 | 1 | 332 |
| 9.852216749 | 2 | 4 | 2 | 1 | 406 |
| 3.827751196 | 1 | 3 | 1 | 1 | 418 |
| 11.48648649 | 2 | 4 | 2 | 1 | 148 |
| 6.02739726 | 3 | 5 | 3 | 1 | 730 |
| 10.90909091 | 1 | 3 | 1 | 1 | 220 |
| 11.2195122 | 2 | 5 | 2 | 1 | 205 |

| | | | | | |
|---|---|---|---|---|---|
| 4.356060606 | 1 | 2 | 1 | 1 | 528 |
| 27.53623188 | 1 | 2 | 1 | 1 | 69 |
| 12.25490196 | 2 | 3 | 2 | 1 | 204 |
| 6.896551724 | 1 | 15 | 1 | 1 | 145 |
| 2.406417112 | 2 | 4 | 2 | 1 | 374 |
| 4.886561955 | 2 | 2 | 2 | 1 | 573 |
| 13.77952756 | 2 | 11 | 2 | 1 | 254 |
| 5.190311419 | 1 | 2 | 1 | 1 | 289 |
| 33.58208955 | 2 | 2 | 2 | 1 | 134 |
| 2.04778157 | 1 | 2 | 1 | 1 | 586 |
| 7.843137255 | 2 | 3 | 2 | 1 | 357 |
| 4.195804196 | 1 | 6 | 1 | 1 | 286 |
| 2.571860817 | 2 | 2 | 2 | 1 | 661 |
| 16.94915254 | 1 | 2 | 1 | 1 | 118 |
| 1.551480959 | 1 | 2 | 1 | 1 | 709 |
| 1.815823606 | 1 | 2 | 1 | 1 | 771 |
| 18.15068493 | 2 | 2 | 2 | 1 | 292 |
| 17.74193548 | 1 | 3 | 1 | 1 | 124 |
| 7.218683652 | 2 | 4 | 2 | 1 | 471 |
| 5.691056911 | 1 | 4 | 1 | 1 | 246 |
| 3.925233645 | 1 | 1 | 1 | 1 | 535 |
| 7.04845815 | 3 | 5 | 3 | 1 | 454 |
| 7.795698925 | 1 | 1 | 1 | 1 | 372 |
| 3.616298812 | 3 | 3 | 3 | 1 | 5890 |
| 7.407407407 | 1 | 2 | 1 | 1 | 135 |
| 13.35616438 | 2 | 3 | 2 | 1 | 292 |
| 0.493733384 | 1 | 1 | 1 | 1 | 2633 |
| 3.555555556 | 1 | 4 | 1 | 1 | 225 |
| 3.472222222 | 1 | 3 | 1 | 1 | 288 |
| 2.678571429 | 1 | 2 | 1 | 1 | 560 |
| 5.24691358 | 1 | 1 | 1 | 1 | 324 |
| 2.035928144 | 1 | 1 | 1 | 1 | 835 |
| 5.861456483 | 3 | 8 | 2 | 1 | 563 |
| 5.081300813 | 1 | 1 | 1 | 1 | 492 |
| 5.245901639 | 1 | 2 | 1 | 1 | 305 |
| 11.73184358 | 1 | 1 | 1 | 1 | 179 |
| 10.61946903 | 2 | 3 | 2 | 1 | 226 |
| 4.017216643 | 1 | 2 | 1 | 1 | 697 |
| 6.289308176 | 1 | 2 | 1 | 1 | 159 |
| 6.906906907 | 1 | 1 | 1 | 1 | 333 |
| 3.529411765 | 2 | 2 | 2 | 1 | 765 |
| 1.85387132 | 1 | 2 | 1 | 1 | 917 |
| 1.899592944 | 1 | 3 | 1 | 1 | 737 |
| 5.785123967 | 1 | 1 | 1 | 1 | 242 |
| 2.02020202 | 1 | 2 | 1 | 1 | 594 |
| 9.644670051 | 1 | 4 | 1 | 1 | 197 |
| 2.05338809 | 1 | 2 | 1 | 1 | 487 |
| 7.603686636 | 1 | 3 | 1 | 1 | 434 |

| | | | | | |
|---|---|---|---|---|---|
| 9.302325581 | 1 | 1 | 1 | 1 | 258 |
| 7.692307692 | 1 | 1 | 1 | 1 | 208 |
| 19.04761905 | 1 | 1 | 1 | 1 | 126 |
| 1.61849711 | 1 | 1 | 1 | 1 | 865 |
| 2.705061082 | 1 | 1 | 1 | 1 | 1146 |
| 2.118644068 | 1 | 1 | 1 | 1 | 472 |
| 10.31390135 | 1 | 2 | 1 | 1 | 223 |
| 6.923076923 | 1 | 1 | 1 | 1 | 130 |
| 2.346368715 | 1 | 2 | 1 | 1 | 895 |
| 2.870813397 | 1 | 1 | 1 | 1 | 627 |
| 6.722689076 | 1 | 1 | 1 | 1 | 238 |
| 1.11662531 | 1 | 2 | 1 | 1 | 806 |
| 3.401360544 | 1 | 2 | 1 | 1 | 294 |
| 1.004304161 | 1 | 2 | 1 | 1 | 1394 |
| 2.360515021 | 1 | 1 | 1 | 1 | 466 |
| 7.189542484 | 1 | 13 | 1 | 1 | 306 |
| 7.608695652 | 1 | 2 | 1 | 1 | 184 |
| 2.095808383 | 1 | 1 | 1 | 1 | 668 |
| 0.743678731 | 1 | 1 | 1 | 1 | 2017 |
| 8.888888889 | 1 | 1 | 1 | 1 | 90 |
| 2.083333333 | 1 | 1 | 1 | 1 | 480 |
| 6.422018349 | 1 | 1 | 1 | 1 | 218 |
| 2.637889688 | 1 | 2 | 1 | 1 | 417 |
| 2.476780186 | 1 | 1 | 1 | 1 | 323 |
| 4.044117647 | 1 | 1 | 1 | 1 | 272 |
| 1.805869074 | 1 | 1 | 1 | 1 | 443 |
| 3.366058906 | 1 | 1 | 1 | 1 | 713 |
| 1.340482574 | 1 | 1 | 1 | 1 | 746 |
| 5.381165919 | 1 | 1 | 1 | 1 | 223 |
| 12.39669421 | 1 | 1 | 1 | 1 | 242 |
| 1.377410468 | 1 | 1 | 1 | 1 | 726 |
| 38.0952381 | 1 | 1 | 1 | 1 | 21 |
| 1.220084467 | 1 | 23 | 1 | 1 | 2131 |
| 2.132998745 | 1 | 2 | 1 | 1 | 797 |
| 2.920962199 | 1 | 1 | 1 | 1 | 582 |
| 1.701323251 | 1 | 1 | 1 | 1 | 529 |
| 4.545454545 | 1 | 3 | 1 | 1 | 264 |
| 1.298701299 | 1 | 2 | 1 | 1 | 770 |
| 0.436046512 | 1 | 2 | 1 | 1 | 4128 |
| 0.795947902 | 1 | 1 | 1 | 1 | 1382 |
| 4.137931034 | 1 | 1 | 1 | 1 | 580 |
| 7.453416149 | 1 | 4 | 1 | 1 | 161 |
| 1.328273245 | 1 | 1 | 1 | 1 | 527 |
| 13.95348837 | 1 | 2 | 1 | 1 | 129 |
| 2.735229759 | 1 | 1 | 1 | 1 | 914 |
| 18.26086957 | 1 | 1 | 1 | 1 | 115 |

| MW [kDa] | calc. pI | Found in Sample: 113 | Found in Sample: 114 |
|---|---|---|---|
| 53.6 | 5.12 | High | High |
| 41.8 | 5.48 | High | High |
| 14.1 | 10.9 | High | High |
| 13.9 | 10.32 | Not Found | Not Found |
| 61 | 5.87 | High | High |
| 13.9 | 10.32 | High | High |
| 15.5 | 11.12 | High | High |
| 49.6 | 4.89 | High | High |
| 49.8 | 4.89 | High | High |
| 49.9 | 4.89 | High | High |
| 15.4 | 11.27 | High | High |
| 98.1 | 5.16 | High | High |
| 32.6 | 4.78 | High | High |
| 83.2 | 5.03 | High | High |
| 10.9 | 8.92 | High | High |
| 117.3 | 5.97 | High | High |
| 82.4 | 4.97 | High | High |
| 47.1 | 7.39 | High | High |
| 58 | 7.71 | High | High |
| 57.7 | 5.07 | High | High |
| 18 | 6.9 | High | High |
| 24.6 | 10.08 | High | High |
| 90.5 | 6 | High | High |
| 76.6 | 4.7 | High | High |
| 38.7 | 7.02 | High | High |
| 17.2 | 8.91 | High | High |
| 56.6 | 5.43 | High | High |
| 24.2 | 11.65 | High | High |
| 35.5 | 8.68 | High | High |
| 72.3 | 5.16 | High | High |
| 73.8 | 6.37 | High | High |
| 48 | 5.38 | High | High |
| 15.1 | 5.16 | High | High |
| 56.7 | 6.35 | High | High |
| 76.1 | 9.44 | High | High |
| 81.5 | 5.33 | High | High |
| 33.1 | 9.29 | High | High |
| 22.2 | 8.38 | High | High |
| 95.3 | 6.83 | High | High |
| 36 | 8.46 | High | High |
| 36 | 7.42 | High | High |
| 46.1 | 5.48 | High | High |
| 45.2 | 8.25 | High | High |
| 17.3 | 6.39 | High | High |
| 77.5 | 8.78 | High | High |
| 23.8 | 10.93 | High | High |
| 22.8 | 6.4 | High | High |

| | | | |
|---|---|---|---|
| 53.3 | 9.42 | High | High |
| 7.8 | 10.7 | High | High |
| 22.7 | 8.34 | High | High |
| 17.3 | 10.2 | High | High |
| 42 | 10.43 | High | High |
| 64.8 | 9.04 | High | High |
| 28.8 | 7.18 | High | High |
| 32.6 | 5.86 | High | High |
| 15.1 | 9.38 | High | High |
| 69.4 | 6.27 | High | High |
| 40.4 | 8.37 | High | High |
| 30.8 | 5.92 | High | High |
| 71.1 | 4.97 | High | High |
| 13.5 | 9.91 | High | High |
| 74.7 | 9.47 | High | High |
| 87.9 | 7.87 | High | High |
| 32.6 | 8.48 | High | High |
| 13.9 | 11.09 | High | High |
| 53.1 | 6.23 | High | High |
| 28 | 4.92 | High | High |
| 57.5 | 6.46 | High | High |
| 50.6 | 9.38 | High | High |
| 39.6 | 9.79 | High | High |
| 628.7 | 6.15 | High | High |
| 14.7 | 5.5 | High | High |
| 32 | 5.91 | High | High |
| 281.5 | 5.71 | High | High |
| 25.3 | 9.76 | High | High |
| 33.7 | 11.25 | High | High |
| 63.1 | 4.94 | High | High |
| 37 | 6.77 | High | High |
| 88.5 | 5.77 | High | High |
| 62.6 | 5.17 | High | High |
| 53.9 | 5.33 | High | High |
| 34.1 | 11.81 | High | High |
| 21.5 | 5.68 | High | High |
| 22.6 | 10.92 | High | High |
| 76.5 | 7.05 | High | High |
| 18.5 | 10.1 | High | High |
| 35.1 | 8.38 | High | High |
| 90.6 | 9.31 | High | High |
| 102.8 | 5.49 | High | High |
| 81 | 7.93 | High | High |
| 26.8 | 9.45 | High | High |
| 65.9 | 9.23 | High | High |
| 22.6 | 5.24 | High | High |
| 56.1 | 7.72 | High | High |
| 47 | 5.44 | High | High |

| | | | |
|---|---|---|---|
| 27.1 | 8.41 | High | High |
| 24.2 | 10.32 | High | High |
| 13.8 | 6.95 | High | High |
| 98.8 | 6.96 | High | High |
| 127.5 | 7.64 | High | High |
| 51.2 | 6.8 | High | High |
| 25.8 | 9.54 | High | High |
| 15.1 | 10.37 | High | High |
| 99.9 | 6.04 | High | High |
| 71.5 | 4.7 | High | High |
| 26.7 | 5.08 | High | High |
| 89.3 | 5.26 | High | High |
| 32.6 | 8.87 | High | High |
| 157.8 | 6.13 | High | High |
| 51.7 | 8.32 | High | High |
| 33.9 | 8.57 | High | High |
| 20.9 | 5.48 | High | High |
| 72.6 | 7.49 | High | High |
| 228.4 | 5.5 | High | High |
| 10.2 | 5.48 | High | High |
| 52.6 | 6.37 | High | High |
| 24.6 | 6.68 | High | High |
| 44.6 | 8.1 | High | High |
| 38 | 4.63 | High | High |
| 29.8 | 5.76 | High | High |
| 48.2 | 8.66 | Medium | Medium |
| 80.6 | 5.52 | High | High |
| 81.4 | 6.29 | Medium | Medium |
| 25.3 | 8.4 | High | High |
| 25.9 | 7.37 | Medium | Medium |
| 82.8 | 5.64 | Not Found | Medium |
| 2.2 | 10.29 | Not Found | Not Found |
| 218 | 8.18 | High | High |
| 89.3 | 4.89 | Not Found | Not Found |
| 64.1 | 5.55 | High | High |
| 57.8 | 5.4 | Medium | Medium |
| 29.9 | 9.73 | Not Found | Not Found |
| 86.9 | 4.83 | High | High |
| 468.8 | 7.12 | Not Found | Not Found |
| 166.4 | 6.79 | High | High |
| 65.9 | 8.97 | Medium | Medium |
| 17.9 | 11.66 | High | High |
| 58.9 | 7.75 | High | High |
| 14.7 | 4.93 | High | High |
| 101.4 | 5.24 | High | High |
| 12.8 | 9.5 | Medium | Medium |

| Found in Sample: 115 | Found in Sample: 116 | Found in Sample: 117 |
|---|---|---|
| High | High | High |
| High | High | High |
| High | High | High |
| Not Found | Not Found | Not Found |
| High | High | High |
| High | High | High |
| High | High | High |
| High | High | High |
| High | High | High |
| High | High | High |
| High | High | High |
| High | High | High |
| High | High | High |
| High | High | High |
| High | High | High |
| High | High | High |
| High | High | High |
| High | High | High |
| High | High | High |
| High | High | High |
| High | High | High |
| High | High | High |
| High | High | High |
| High | High | High |
| High | High | High |
| High | High | High |
| High | High | High |
| High | High | High |
| High | High | High |
| High | High | High |
| High | High | High |
| High | High | High |
| High | High | High |
| High | High | High |
| High | High | High |
| High | High | High |
| High | High | High |
| High | High | High |
| High | High | High |
| High | High | High |
| High | High | High |
| High | High | High |
| High | High | High |
| High | High | High |
| High | High | High |
| High | High | High |
| High | High | High |

| | | |
|---|---|---|
| High | High | High |
| High | High | High |
| High | High | High |
| High | High | High |
| High | High | High |
| High | High | High |
| High | High | High |
| High | High | High |
| High | High | High |
| High | High | High |
| High | High | High |
| High | High | High |
| High | High | High |
| High | High | High |
| High | High | High |
| High | High | High |
| High | High | High |
| High | High | High |
| High | High | High |
| High | High | High |
| High | High | High |
| High | High | High |
| High | High | High |
| High | High | High |
| High | High | High |
| High | High | High |
| High | High | High |
| High | High | High |
| High | High | High |
| High | High | High |
| High | High | High |
| High | High | High |
| High | High | High |
| High | High | High |
| High | High | High |
| High | High | High |
| High | High | High |
| High | High | High |
| High | High | High |
| High | High | High |
| High | High | High |
| High | High | High |
| High | High | High |
| High | High | High |
| High | High | High |
| High | High | High |
| High | High | High |
| High | High | High |

| | | |
|---|---|---|
| High | High | High |
| High | High | High |
| High | High | High |
| High | High | High |
| High | High | High |
| High | High | High |
| High | High | High |
| High | High | High |
| High | High | High |
| High | High | High |
| High | High | High |
| High | High | High |
| High | High | High |
| High | High | High |
| High | High | High |
| High | High | High |
| High | High | High |
| High | High | High |
| High | High | High |
| High | High | High |
| High | High | High |
| High | High | High |
| High | High | High |
| High | High | High |
| High | High | High |
| Medium | Medium | Medium |
| High | High | High |
| Medium | Medium | Medium |
| High | High | High |
| Medium | Medium | Medium |
| Not Found | Medium | Not Found |
| Not Found | Not Found | Not Found |
| High | High | Medium |
| Not Found | Not Found | Not Found |
| High | High | High |
| Medium | Medium | Medium |
| Medium | Not Found | Not Found |
| High | High | High |
| Medium | Not Found | Not Found |
| High | High | High |
| Medium | Medium | Medium |
| High | High | High |
| High | High | High |
| High | High | High |
| High | High | High |
| Medium | Medium | Medium |

| Found in Sample: 118 | Found in Sample: 119 | Found in Sample: 121 | Modificatio |
|---|---|---|---|
| High | High | High | |
| High | High | High | |
| High | High | High | |
| Not Found | Not Found | Not Found | |
| High | High | High | |
| High | High | High | |
| High | High | High | |
| High | High | High | |
| High | High | High | |
| High | High | High | |
| High | High | High | |
| High | High | High | |
| High | High | High | |
| High | High | High | |
| High | High | High | |
| High | High | High | |
| High | High | High | |
| High | High | High | |
| High | High | High | |
| High | High | High | |
| High | High | High | |
| High | High | High | |
| High | High | High | |
| High | High | High | |
| High | High | High | |
| High | High | High | |
| High | High | High | |
| High | High | High | |
| High | High | High | |
| High | High | High | |
| High | High | High | |
| High | High | High | |
| High | High | High | |
| High | High | High | |
| High | High | High | |
| High | High | High | |
| High | High | High | |
| High | High | High | |
| High | High | High | |
| High | High | High | |
| High | High | High | |
| High | High | High | |
| High | High | High | |
| High | High | High | |
| High | High | High | |
| High | High | High | |
| High | High | High | |

| | | |
|---|---|---|
| High | High | High |
| High | High | High |
| High | High | High |
| High | High | High |
| High | High | High |
| High | High | High |
| High | High | High |
| High | High | High |
| High | High | High |
| High | High | High |
| High | High | High |
| High | High | High |
| High | High | High |
| High | High | High |
| High | High | High |
| High | High | High |
| High | High | High |
| High | High | High |
| High | High | High |
| High | High | High |
| High | High | High |
| High | High | High |
| High | High | High |
| High | High | High |
| High | High | High |
| High | High | High |
| High | High | High |
| High | High | High |
| High | High | High |
| High | High | High |
| High | High | High |
| High | High | High |
| High | High | High |
| High | High | High |
| High | High | High |
| High | High | High |
| High | High | High |
| High | High | High |
| High | High | High |
| High | High | High |
| High | High | High |
| High | High | High |
| High | High | High |
| High | High | High |
| High | High | High |
| High | High | High |
| High | High | High |
| High | High | High |

| | | |
|---|---|---|
| High | High | High |
| High | High | High |
| High | High | High |
| High | High | High |
| High | High | High |
| High | High | High |
| High | High | High |
| High | High | High |
| High | High | High |
| High | High | High |
| High | High | High |
| High | High | High |
| High | High | High |
| High | High | High |
| High | High | High |
| High | High | High |
| High | High | High |
| High | High | High |
| High | High | High |
| High | High | High |
| High | High | High |
| High | High | High |
| High | High | High |
| High | High | High |
| High | High | High |
| Medium | Medium | Medium |
| High | High | High |
| Medium | Medium | Medium |
| High | High | High |
| Medium | Medium | Medium |
| Not Found | Not Found | Not Found |
| Not Found | Not Found | Not Found |
| High | High | High |
| Not Found | Not Found | Not Found |
| High | High | High |
| Medium | Medium | Medium |
| Not Found | Not Found | Not Found |
| High | High | High |
| Not Found | Not Found | Not Found |
| High | High | High |
| Medium | Medium | Medium |
| High | High | High |
| High | High | High |
| High | High | High |
| High | High | High |
| Medium | Medium | Medium |

| Abundance Ratio: 114/113 | Abundance Ratio: 113/114 | Abundance Ratio: 116/115 |
|---|---|---|
| 1.093 | 0.915 | 1.107 |
| 0.726 | 1.378 | 0.88 |
| 1.278 | 0.782 | 1.142 |
| 1.124 | 0.89 | 1.076 |
| 1.028 | 0.973 | 1.115 |
| 1.345 | 0.744 | 1.286 |
| 1.047 | 0.955 | 0.915 |
| 0.84 | 1.19 | 0.94 |
| 0.855 | 1.17 | 0.845 |
| 0.964 | 1.037 | 1.147 |
| 1.241 | 0.806 | 1.022 |
| 0.928 | 1.078 | 0.919 |
| 0.942 | 1.062 | 0.938 |
| 0.999 | 1.001 | 0.948 |
| 0.978 | 1.022 | 1.122 |
| 0.97 | 1.031 | 0.905 |
| 0.89 | 1.123 | 0.847 |
| 0.839 | 1.192 | 0.821 |
| 0.937 | 1.068 | 0.954 |
| 0.817 | 1.224 | 0.85 |
| 0.865 | 1.156 | 0.848 |
| 0.904 | 1.106 | 0.915 |
| 1.08 | 0.926 | 1.206 |
| 0.963 | 1.039 | 0.909 |
| 0.865 | 1.157 | 0.796 |
| 0.85 | 1.176 | 0.887 |
| 0.945 | 1.058 | 0.91 |
| 0.951 | 1.051 | 0.947 |
| 1.006 | 0.994 | 1.084 |
| 0.93 | 1.076 | 1.05 |
| 1.046 | 0.956 | 1.019 |
| 0.968 | 1.033 | 0.976 |
| 0.786 | 1.273 | 0.848 |
| 0.914 | 1.095 | 1.035 |
| 1.02 | 0.981 | 0.99 |
| 1.062 | 0.941 | 1.14 |
| 0.941 | 1.063 | 0.85 |
| 0.847 | 1.18 | 0.834 |
| 0.868 | 1.153 | 0.937 |
| 0.933 | 1.072 | 0.925 |
| 1.171 | 0.854 | 1.124 |
| 0.982 | 1.018 | 0.947 |
| 1.054 | 0.949 | 1.011 |
| 0.911 | 1.098 | 0.878 |
| 0.924 | 1.082 | 0.969 |
| 0.986 | 1.014 | 0.882 |
| 0.905 | 1.105 | 0.885 |

| | | |
|---|---|---|
| 1.159 | 0.863 | 1.128 |
| 0.882 | 1.134 | 0.938 |
| 0.844 | 1.184 | 0.897 |
| 0.814 | 1.228 | 0.793 |
| 0.808 | 1.237 | 0.822 |
| 0.687 | 1.456 | 0.783 |
| 0.82 | 1.22 | 0.871 |
| 0.784 | 1.275 | 0.824 |
| 0.862 | 1.159 | 0.919 |
| 0.99 | 1.01 | 1 |
| 1.09 | 0.918 | 0.987 |
| 0.403 | 2.48 | 0.499 |
| 1.066 | 0.938 | 1.088 |
| 0.998 | 1.002 | 0.843 |
| 0.666 | 1.5 | 0.902 |
| 0.969 | 1.032 | 0.845 |
| 0.889 | 1.125 | 0.928 |
| 0.976 | 1.024 | 0.924 |
| 1.016 | 0.984 | 0.944 |
| 0.445 | 2.248 | 0.496 |
| 0.9 | 1.112 | 0.982 |
| 1.004 | 0.996 | 0.966 |
| 1.121 | 0.892 | 0.725 |
| 1.074 | 0.931 | 1.022 |
| 1.136 | 0.88 | 1.147 |
| 1.016 | 0.984 | 1.041 |
| 0.863 | 1.159 | 1.015 |
| 0.859 | 1.164 | 0.847 |
| 1.014 | 0.987 | 1.027 |
| 0.855 | 1.17 | 0.976 |
| 0.645 | 1.549 | 0.652 |
| 0.795 | 1.258 | 0.868 |
| 0.866 | 1.155 | 0.944 |
| 0.815 | 1.227 | 0.792 |
| 0.891 | 1.123 | 1.039 |
| 0.891 | 1.122 | 0.914 |
| 1.094 | 0.914 | 1.048 |
| 1.012 | 0.989 | 0.963 |
| 0.723 | 1.384 | 0.736 |
| 0.97 | 1.031 | 0.957 |
| 0.829 | 1.206 | 0.863 |
| 1.08 | 0.926 | 0.983 |
| 0.929 | 1.076 | 0.862 |
| 0.865 | 1.157 | 1.011 |
| 1.017 | 0.984 | 0.976 |
| 0.905 | 1.105 | 0.822 |
| 0.75 | 1.333 | 0.811 |
| 1.002 | 0.998 | 0.878 |

| | | |
|---|---|---|
| 0.918 | 1.09 | 0.95 |
| 1.236 | 0.809 | 1.11 |
| 0.947 | 1.056 | 1.013 |
| 0.988 | 1.012 | 0.881 |
| 1.303 | 0.767 | 0.833 |
| 1.056 | 0.947 | 1.092 |
| 0.691 | 1.446 | 0.932 |
| 0.768 | 1.302 | 0.82 |
| 1.031 | 0.97 | 1.092 |
| 1.034 | 0.967 | 1.196 |
| 1.044 | 0.958 | 0.983 |
| 0.566 | 1.768 | 0.728 |
| 0.855 | 1.169 | 0.952 |
| 1.02 | 0.98 | 1.002 |
| 0.863 | 1.159 | 0.898 |
| 1.365 | 0.733 | 0.719 |
| 0.974 | 1.026 | 0.924 |
| 0.756 | 1.322 | 0.807 |
| 0.9 | 1.111 | 0.874 |
| 0.693 | 1.444 | 0.895 |
| 0.878 | 1.139 | 0.929 |
| 0.84 | 1.191 | 0.803 |
| 0.997 | 1.003 | 0.936 |
| 0.928 | 1.077 | 1.1 |
| 0.556 | 1.798 | 0.558 |
| 0.699 | 1.431 | 0.852 |
| 1.125 | 0.889 | 1.074 |
| 0.603 | 1.659 | 0.61 |
| 0.841 | 1.189 | 0.749 |
| 100 | 0.01 | 100 |
| 2.673 | 0.374 | 0.889 |
| 0.987 | 1.013 | 0.993 |
| 0.885 | 1.13 | 1.032 |
| | | 0.01 |
| 0.729 | 1.372 | 0.71 |
| 0.749 | 1.335 | 0.943 |
| 1.093 | 0.915 | 1.586 |
| 0.9 | 1.111 | 0.934 |
| 0.926 | 1.08 | 0.932 |
| 0.91 | 1.099 | 0.962 |
| 1.217 | 0.821 | 1.192 |
| 1.016 | 0.984 | 0.991 |

| Abundance Ratio: 115/116 | Abundance Ratio: 118 /117 | Abundance Ratio: 119 /117 |
|---|---|---|
| 0.904 | 1.507 | 0.978 |
| 1.137 | 0.943 | 1.069 |
| 0.875 | 0.963 | 0.939 |
| 0.929 | 0.847 | 0.926 |
| 0.897 | 0.922 | 0.936 |
| 0.778 | 0.964 | 0.915 |
| 1.093 | 0.694 | 0.944 |
| 1.063 | 0.965 | 1.037 |
| 1.183 | 0.932 | 1.069 |
| 0.872 | 0.788 | 0.962 |
| 0.978 | 0.877 | 0.941 |
| 1.088 | 1.013 | 1.045 |
| 1.066 | 1.019 | 0.979 |
| 1.055 | 1.018 | 1.059 |
| 0.891 | 0.922 | 0.973 |
| 1.106 | 0.841 | 0.906 |
| 1.18 | 1.006 | 1.07 |
| 1.218 | 0.981 | 1.107 |
| 1.048 | 0.958 | 1.031 |
| 1.176 | 1.008 | 1.079 |
| 1.179 | 0.986 | 1.059 |
| 1.093 | 0.893 | 1.096 |
| 0.829 | 0.909 | 0.906 |
| 1.101 | 0.97 | 1.034 |
| 1.256 | 1.021 | 1.142 |
| 1.128 | 1.005 | 1.045 |
| 1.098 | 1.179 | 0.977 |
| 1.056 | 1.005 | 1.04 |
| 0.923 | 0.919 | 0.937 |
| 0.952 | 0.927 | 1.053 |
| 0.981 | 0.975 | 0.981 |
| 1.024 | 1.029 | 0.889 |
| 1.18 | 0.978 | 1.025 |
| 0.966 | 0.94 | 1.029 |
| 1.01 | 0.982 | 1.004 |
| 0.877 | 0.974 | 0.932 |
| 1.177 | 0.986 | 1.036 |
| 1.199 | 0.951 | 1.075 |
| 1.068 | 0.967 | 0.965 |
| 1.081 | 0.965 | 1.067 |
| 0.89 | 0.942 | 0.97 |
| 1.056 | 1.19 | 0.915 |
| 0.989 | 1.011 | 1.101 |
| 1.139 | 0.871 | 0.988 |
| 1.031 | 1.001 | 1.065 |
| 1.134 | 0.935 | 1.104 |
| 1.13 | 0.989 | 0.965 |

| | | |
|---|---|---|
| 0.886 | 0.918 | 1.043 |
| 1.066 | 1.006 | 1.04 |
| 1.115 | 0.915 | 0.958 |
| 1.262 | 1.048 | 1.11 |
| 1.217 | 1.025 | 1.086 |
| 1.278 | 0.939 | 1.093 |
| 1.148 | 1 | 1.055 |
| 1.213 | 1.171 | 1.2 |
| 1.088 | 0.951 | 0.994 |
| 1 | 0.915 | 0.948 |
| 1.013 | 0.877 | 0.926 |
| 2.005 | 1.897 | 1.273 |
| 0.919 | 0.963 | 1.082 |
| 1.186 | 0.97 | 1.044 |
| 1.109 | 1.048 | 1.094 |
| 1.184 | 0.932 | 1.096 |
| 1.077 | 0.998 | 1.05 |
| 1.082 | 0.928 | 0.972 |
| 1.059 | 0.981 | 1.056 |
| 2.017 | 2.808 | 2.507 |
| 1.018 | 0.826 | 1.077 |
| 1.035 | 0.986 | 0.951 |
| 1.378 | 0.906 | 0.921 |
| 0.978 | 0.974 | 0.984 |
| 0.872 | 1.1 | 1.375 |
| 0.961 | 0.971 | 1.006 |
| 0.985 | 1.195 | 1.052 |
| 1.181 | 1.394 | 1.122 |
| 0.974 | 1.019 | 0.898 |
| 1.025 | 0.922 | 0.977 |
| 1.535 | 0.899 | 0.852 |
| 1.152 | 1.166 | 1.163 |
| 1.06 | 1.101 | 1.025 |
| 1.262 | 1 | 1.056 |
| 0.963 | 1.009 | 1.057 |
| 1.095 | 0.894 | 1.015 |
| 0.954 | 1.054 | 0.902 |
| 1.039 | 0.919 | 1.246 |
| 1.359 | 1.157 | 1.2 |
| 1.045 | 0.934 | 1.035 |
| 1.159 | 1.017 | 1.041 |
| 1.018 | 1.03 | 0.975 |
| 1.16 | 1.082 | 1.22 |
| 0.989 | 0.859 | 0.967 |
| 1.025 | 0.978 | 1.124 |
| 1.216 | 0.874 | 0.919 |
| 1.233 | 1.087 | 1.058 |
| 1.139 | 0.909 | 1.032 |

| | | |
|---|---|---|
| 1.053 | 0.917 | 0.986 |
| 0.901 | 0.93 | 1.052 |
| 0.987 | 0.985 | 1.105 |
| 1.136 | 0.974 | 1.002 |
| 1.2 | 1.082 | 1.008 |
| 0.916 | 0.859 | 0.929 |
| 1.073 | 1.114 | 1.189 |
| 1.22 | 0.816 | 0.982 |
| 0.916 | 0.908 | 0.979 |
| 0.836 | 0.999 | 0.966 |
| 1.018 | 1.073 | 0.915 |
| 1.373 | 1.021 | 1.067 |
| 1.05 | 1.014 | 0.978 |
| 0.998 | 0.958 | 1.098 |
| 1.114 | 0.871 | 0.94 |
| 1.391 | 0.946 | 0.81 |
| 1.082 | 1.008 | 1.018 |
| 1.239 | 0.91 | 0.626 |
| 1.144 | 1.039 | 0.999 |
| 1.118 | 1.261 | 1.349 |
| 1.076 | 0.755 | 1.066 |
| 1.245 | 0.889 | 0.849 |
| 1.068 | 0.95 | 0.925 |
| 0.909 | 1.434 | 1.586 |
| 1.792 | 0.956 | 0.81 |
| 1.174 | 0.986 | 1.029 |
| 0.931 | 0.896 | 0.844 |
| 1.639 | 1.164 | 1.248 |
| 1.336 | 1.014 | 1.244 |
| 0.01 | | |
| 1.125 | 100 | 100 |
| 1.007 | 0.911 | 0.935 |
| 0.969 | 0.981 | 1.164 |
| 100 | | |
| 1.408 | 1.067 | 1.189 |
| 1.061 | 1.044 | 1.043 |
| 0.631 | 0.967 | 0.898 |
| 1.071 | 0.982 | 1.047 |
| 1.073 | 0.988 | 0.973 |
| 1.04 | 0.825 | 0.883 |
| 0.839 | 0.826 | 0.929 |
| 1.009 | 0.843 | 1.006 |

| Abundance Ratio: 121 /117 | oundance Ratio: (F1, 116) / (F1, 11 | oundance Ratio: (F1, 117) / (F1, 11 |
|---|---|---|
| 1.068 | 0.62 | 0.664 |
| 1.139 | 1.151 | 1.06 |
| 0.872 | 1.128 | 1.039 |
| 0.865 | 1.246 | 1.18 |
| 0.895 | 1.359 | 1.085 |
| 0.942 | 1.118 | 1.038 |
| 0.76 | 1.093 | 1.44 |
| 0.948 | 0.952 | 1.036 |
| 1.044 | 0.941 | 1.073 |
| 0.976 | 0.902 | 1.268 |
| 1.019 | 1.206 | 1.14 |
| 1.03 | 0.964 | 0.987 |
| 0.998 | 1.147 | 0.981 |
| 1.138 | 0.918 | 0.982 |
| 0.921 | 1.472 | 1.085 |
| 0.853 | 1.093 | 1.189 |
| 1.09 | 0.959 | 0.994 |
| 1.161 | 0.957 | 1.019 |
| 1.012 | 0.973 | 1.044 |
| 1.044 | 0.878 | 0.993 |
| 1.122 | 1.015 | 1.014 |
| 1.057 | 0.787 | 1.12 |
| 0.781 | 1.244 | 1.101 |
| 1.055 | 0.985 | 1.031 |
| 1.287 | 0.883 | 0.979 |
| 0.913 | 0.913 | 0.995 |
| 0.915 | 0.771 | 0.848 |
| 1.145 | 1.002 | 0.995 |
| 0.962 | 1.261 | 1.088 |
| 1.07 | 1.162 | 1.079 |
| 0.955 | 1.113 | 1.026 |
| 0.896 | 0.826 | 0.971 |
| 1.034 | 0.968 | 1.022 |
| 0.987 | 1.28 | 1.064 |
| 1.018 | 1.116 | 1.018 |
| 0.948 | 0.959 | 1.027 |
| 1.103 | 0.94 | 1.014 |
| 1.138 | 0.954 | 1.051 |
| 1.024 | 0.985 | 1.034 |
| 0.992 | 0.951 | 1.036 |
| 0.879 | 0.981 | 1.061 |
| 0.877 | 0.752 | 0.84 |
| 1.081 | 0.939 | 0.989 |
| 0.923 | 1.024 | 1.149 |
| 1.074 | 1.118 | 0.999 |
| 1.079 | 1.011 | 1.07 |
| 1.027 | 0.976 | 1.012 |

| | | |
|---|---|---|
| 0.942 | 1.297 | 1.09 |
| 1.076 | 1.022 | 0.995 |
| 1.062 | 1.219 | 1.093 |
| 1.26 | 0.815 | 0.954 |
| 1.099 | 0.891 | 0.976 |
| 1.281 | 1.16 | 1.065 |
| 1.063 | 1.05 | 1 |
| 1.277 | 0.844 | 0.854 |
| 1.008 | 0.957 | 1.051 |
| 0.914 | 1.065 | 1.093 |
| 0.844 | 1.004 | 1.14 |
| 2.438 | 0.917 | 0.527 |
| 1.02 | 0.993 | 1.038 |
| 1.009 | 0.963 | 1.031 |
| 0.876 | 1.066 | 0.955 |
| 0.832 | 0.881 | 1.073 |
| 1.055 | 0.949 | 1.002 |
| 0.87 | 1.038 | 1.077 |
| 1.28 | 1.008 | 1.019 |
| 0.963 | 0.651 | 0.356 |
| 1.04 | 1.174 | 1.21 |
| 0.965 | 0.998 | 1.014 |
| 0.592 | 0.814 | 1.103 |
| 0.941 | 1.001 | 1.026 |
| 1.349 | 1.867 | 0.909 |
| 0.931 | 1.221 | 1.029 |
| 1.133 | 0.842 | 0.837 |
| 2.33 | 0.783 | 0.717 |
| 0.823 | 0.88 | 0.981 |
| 0.858 | 1.071 | 1.084 |
| 1.003 | 0.76 | 1.112 |
| 1.042 | 1.009 | 0.857 |
| 1.025 | 0.941 | 0.908 |
| 1.008 | 0.933 | 1 |
| 1.049 | 1.004 | 0.991 |
| 0.954 | 1.046 | 1.119 |
| 0.806 | 0.785 | 0.949 |
| 1.208 | 1.172 | 1.089 |
| 1.378 | 1.008 | 0.864 |
| 1.072 | 1.22 | 1.071 |
| 1.042 | 1.063 | 0.983 |
| 1.099 | 0.973 | 0.971 |
| 1.161 | 1.345 | 0.924 |
| 1.029 | 0.999 | 1.164 |
| 1.158 | 1.191 | 1.022 |
| 0.935 | 0.968 | 1.145 |
| 1.09 | 0.785 | 0.92 |
| 0.806 | 1.071 | 1.101 |

| | | |
|---|---|---|
| 1.046 | 0.931 | 1.091 |
| 0.898 | 0.866 | 1.075 |
| 0.99 | 0.993 | 1.015 |
| 0.954 | 0.956 | 1.027 |
| 0.754 | 1.015 | 0.924 |
| 0.825 | 1.199 | 1.165 |
| 1.166 | 0.978 | 0.898 |
| 1.05 | 0.875 | 1.225 |
| 0.888 | 1.182 | 1.101 |
| 0.959 | 1.21 | 1.001 |
| 1.12 | 0.902 | 0.932 |
| 1.22 | 1 | 0.979 |
| 0.925 | 0.828 | 0.986 |
| 1.063 | 1.186 | 1.043 |
| 0.985 | 1.053 | 1.148 |
| 0.992 | 1.012 | 1.057 |
| 0.946 | 1.01 | 0.992 |
| 1.107 | 1.056 | 1.099 |
| 1.074 | 0.993 | 0.962 |
| 1.724 | 0.908 | 0.793 |
| 1.134 | 1.243 | 1.324 |
| 1.038 | 0.927 | 1.124 |
| 0.995 | 1.137 | 1.052 |
| 1.313 | 1.17 | 0.697 |
| 0.777 | 0.749 | 1.046 |
| 0.862 | 0.952 | 1.014 |
| 0.897 | 1.073 | 1.116 |
| 1.722 | 0.907 | 0.859 |
| 1.214 | 1.129 | 0.986 |
| | 100 | |
| 100 | 1.134 | 0.01 |
| 1.028 | 1.198 | 1.098 |
| 1.027 | 1.1 | 1.02 |
| 1.033 | 1.137 | 0.937 |
| 1.02 | 1 | 0.958 |
| 0.872 | 1.418 | 1.034 |
| 1.007 | 0.812 | 1.019 |
| 0.934 | 1.205 | 1.012 |
| 0.844 | 0.968 | 1.212 |
| 0.88 | 0.912 | 1.211 |
| 0.883 | 1.218 | 1.186 |

| oundance Ratio: (F1, 119) / (F1, 11 | oundance Ratio: (F1, 121) / (F1, 11 | oundance Ratio: (F1, 113) / (F1, 11 |
|---|---|---|
| 0.649 | 0.709 | 0.877 |
| 1.133 | 1.207 | 1.167 |
| 0.975 | 0.905 | 0.972 |
| 1.093 | 1.021 | 1.144 |
| 1.016 | 0.971 | 1.272 |
| 0.949 | 0.977 | 1.011 |
| 1.36 | 1.094 | 0.797 |
| 1.074 | 0.982 | 0.937 |
| 1.147 | 1.12 | 0.888 |
| 1.22 | 1.238 | 0.923 |
| 1.073 | 1.162 | 1.18 |
| 1.032 | 1.017 | 0.934 |
| 0.96 | 0.979 | 1.246 |
| 1.041 | 1.118 | 0.846 |
| 1.056 | 0.999 | 1.364 |
| 1.077 | 1.014 | 0.986 |
| 1.064 | 1.084 | 0.966 |
| 1.128 | 1.183 | 0.862 |
| 1.076 | 1.057 | 0.88 |
| 1.071 | 1.036 | 0.864 |
| 1.073 | 1.137 | 0.904 |
| 1.227 | 1.183 | 0.643 |
| 0.997 | 0.86 | 1.235 |
| 1.065 | 1.088 | 0.892 |
| 1.118 | 1.26 | 0.802 |
| 1.04 | 0.909 | 0.893 |
| 0.829 | 0.776 | 0.973 |
| 1.035 | 1.139 | 1.019 |
| 1.019 | 1.046 | 1.207 |
| 1.136 | 1.154 | 1.065 |
| 1.006 | 0.979 | 1.124 |
| 0.863 | 0.87 | 1 |
| 1.047 | 1.057 | 1.161 |
| 1.094 | 1.05 | 1.173 |
| 1.022 | 1.037 | 1.069 |
| 0.957 | 0.973 | 0.935 |
| 1.05 | 1.118 | 0.874 |
| 1.13 | 1.196 | 0.899 |
| 0.997 | 1.059 | 1.016 |
| 1.105 | 1.028 | 0.792 |
| 1.03 | 0.933 | 0.817 |
| 0.769 | 0.737 | 0.916 |
| 1.089 | 1.069 | 0.805 |
| 1.134 | 1.06 | 1.073 |
| 1.064 | 1.073 | 1.148 |
| 1.181 | 1.154 | 0.818 |
| 0.976 | 1.039 | 1.112 |

| | | |
|---|---|---|
| 1.136 | 1.027 | 0.965 |
| 1.034 | 1.07 | 0.979 |
| 1.046 | 1.16 | 1.169 |
| 1.059 | 1.202 | 0.927 |
| 1.06 | 1.072 | 1.03 |
| 1.164 | 1.364 | 1.433 |
| 1.056 | 1.063 | 1.136 |
| 1.025 | 1.091 | 0.837 |
| 1.045 | 1.059 | 0.902 |
| 1.037 | 0.999 | 1.083 |
| 1.056 | 0.962 | 0.997 |
| 0.671 | 1.285 | 3.032 |
| 1.123 | 1.059 | 0.862 |
| 1.077 | 1.04 | 0.982 |
| 1.044 | 0.836 | 1.575 |
| 1.176 | 0.893 | 0.977 |
| 1.052 | 1.057 | 0.859 |
| 1.047 | 0.937 | 1.024 |
| 1.076 | 1.304 | 0.902 |
| 0.893 | 0.343 | 1.455 |
| 1.303 | 1.259 | 0.925 |
| 0.965 | 0.979 | 1.048 |
| 1.016 | 0.653 | 0.733 |
| 1.01 | 0.965 | 0.927 |
| 1.25 | 1.227 | 1.353 |
| 1.036 | 0.959 | 1.17 |
| 0.881 | 0.949 | 1.018 |
| 0.805 | 1.671 | 1.471 |
| 0.881 | 0.807 | 1.089 |
| 1.059 | 0.93 | 1.353 |
| 0.948 | 1.116 | 1.603 |
| 0.997 | 0.893 | 1.245 |
| 0.931 | 0.931 | 1.045 |
| 1.057 | 1.008 | 1.078 |
| 1.047 | 1.04 | 1.027 |
| 1.136 | 1.068 | 0.926 |
| 0.856 | 0.765 | 1.029 |
| 1.356 | 1.314 | 0.833 |
| 1.037 | 1.191 | 1.409 |
| 1.108 | 1.148 | 1.069 |
| 1.024 | 1.024 | 1.205 |
| 0.947 | 1.068 | 1.077 |
| 1.128 | 1.073 | 1.059 |
| 1.125 | 1.198 | 0.848 |
| 1.149 | 1.184 | 1.158 |
| 1.052 | 1.07 | 0.983 |
| 0.973 | 1.003 | 0.918 |
| 1.136 | 0.887 | 0.903 |

| | | |
|---|---|---|
| 1.075 | 1.142 | 0.776 |
| 1.131 | 0.965 | 0.61 |
| 1.122 | 1.005 | 0.988 |
| 1.029 | 0.979 | 1.06 |
| 0.931 | 0.697 | 0.504 |
| 1.082 | 0.961 | 1.059 |
| 1.068 | 1.047 | 1.354 |
| 1.203 | 1.287 | 0.834 |
| 1.078 | 0.978 | 1.066 |
| 0.967 | 0.96 | 1.054 |
| 0.853 | 1.045 | 0.934 |
| 1.044 | 1.194 | 1.574 |
| 0.965 | 0.913 | 1.06 |
| 1.145 | 1.109 | 1.066 |
| 1.08 | 1.131 | 1.028 |
| 0.856 | 1.048 | 0.916 |
| 1.01 | 0.939 | 0.968 |
| 0.688 | 1.217 | 1.839 |
| 0.962 | 1.034 | 1.104 |
| 1.07 | 1.367 | 0.945 |
| 1.411 | 1.501 | 0.983 |
| 0.955 | 1.167 | 1.317 |
| 0.974 | 1.047 | 1.105 |
| 1.106 | 0.916 | 0.954 |
| 0.847 | 0.813 | 1.507 |
| 1.044 | 0.874 | 1.082 |
| 0.942 | 1.001 | 0.987 |
| 1.072 | 1.479 | 1.237 |
| 1.227 | 1.197 | 1.263 |
| 0.519 | 1.038 | 1.276 |
| 1.027 | 1.13 | 1.169 |
| 1.187 | 1.047 | 0.921 |
| 1.115 | 0.969 | 1.38 |
| 0.998 | 0.977 | 1.091 |
| 0.929 | 0.902 | 1.087 |
| 1.067 | 1.026 | 0.815 |
| 0.985 | 0.945 | 1.306 |
| 1.07 | 1.023 | 1.028 |
| 1.126 | 1.065 | 0.694 |
| 1.193 | 1.047 | 1.005 |

| oundance Ratio: (F1, 114) / (F1, 11 | oundance Ratio: (F1, 115) / (F1, 11 | oundance Ratio: (F1, 116) / (F1, 11 |
|---|---|---|
| 0.959 | 0.864 | 0.956 |
| 0.847 | 1.154 | 1.016 |
| 1.242 | 1.013 | 1.157 |
| 1.286 | 1.06 | 1.14 |
| 1.307 | 1.2 | 1.338 |
| 1.36 | 0.916 | 1.178 |
| 0.834 | 0.878 | 0.804 |
| 0.788 | 0.943 | 0.886 |
| 0.759 | 0.971 | 0.821 |
| 0.89 | 0.644 | 0.739 |
| 1.465 | 1.1 | 1.124 |
| 0.867 | 1.016 | 0.934 |
| 1.173 | 1.274 | 1.195 |
| 0.845 | 0.93 | 0.882 |
| 1.334 | 1.242 | 1.394 |
| 0.957 | 1.122 | 1.015 |
| 0.86 | 1.063 | 0.901 |
| 0.724 | 1.033 | 0.848 |
| 0.824 | 0.947 | 0.904 |
| 0.706 | 0.965 | 0.821 |
| 0.782 | 1.115 | 0.946 |
| 0.581 | 0.701 | 0.641 |
| 1.333 | 1.034 | 1.248 |
| 0.858 | 1.018 | 0.925 |
| 0.694 | 0.992 | 0.79 |
| 0.759 | 0.99 | 0.877 |
| 0.919 | 1.022 | 0.93 |
| 0.969 | 1.023 | 0.969 |
| 1.214 | 1.142 | 1.238 |
| 0.99 | 0.974 | 1.023 |
| 1.176 | 1.085 | 1.106 |
| 0.968 | 0.98 | 0.957 |
| 0.913 | 1.09 | 0.924 |
| 1.072 | 1.131 | 1.17 |
| 1.09 | 1.103 | 1.092 |
| 0.994 | 0.879 | 1.002 |
| 0.822 | 1.053 | 0.895 |
| 0.762 | 1.012 | 0.844 |
| 0.882 | 1.054 | 0.988 |
| 0.739 | 0.931 | 0.861 |
| 0.956 | 0.848 | 0.953 |
| 0.9 | 1.033 | 0.979 |
| 0.849 | 0.853 | 0.862 |
| 0.977 | 1.029 | 0.903 |
| 1.061 | 1.084 | 1.051 |
| 0.807 | 0.971 | 0.856 |
| 1.007 | 1.13 | 1 |

| | | |
|---|---|---|
| 1.119 | 1.011 | 1.141 |
| 0.864 | 1.054 | 0.988 |
| 0.987 | 1.299 | 1.165 |
| 0.755 | 0.971 | 0.77 |
| 0.833 | 1.023 | 0.84 |
| 0.984 | 1.273 | 0.996 |
| 0.931 | 1.143 | 0.995 |
| 0.656 | 1 | 0.824 |
| 0.778 | 0.997 | 0.916 |
| 1.072 | 1.028 | 1.028 |
| 1.087 | 0.964 | 0.951 |
| 1.222 | 2.742 | 1.367 |
| 0.919 | 0.813 | 0.884 |
| 0.98 | 1.061 | 0.894 |
| 1.05 | 1.132 | 1.02 |
| 0.947 | 0.887 | 0.749 |
| 0.764 | 0.972 | 0.902 |
| 0.999 | 1.072 | 0.991 |
| 0.917 | 0.992 | 0.936 |
| 0.647 | 1.471 | 0.729 |
| 0.832 | 0.917 | 0.901 |
| 1.052 | 1.071 | 1.035 |
| 0.821 | 1.104 | 0.801 |
| 0.995 | 0.97 | 0.991 |
| 1.537 | 1.303 | 1.494 |
| 1.188 | 1.133 | 1.179 |
| 0.878 | 0.942 | 0.956 |
| 1.264 | 1.15 | 0.973 |
| 1.103 | 0.973 | 1 |
| 1.157 | 1.037 | 1.012 |
| 1.034 | 1.231 | 0.802 |
| 0.99 | 1.167 | 1.012 |
| 0.905 | 1.072 | 1.011 |
| 0.878 | 1.114 | 0.883 |
| 0.915 | 0.924 | 0.959 |
| 0.825 | 1.008 | 0.921 |
| 1.125 | 0.875 | 0.917 |
| 0.843 | 0.898 | 0.864 |
| 1.019 | 1.321 | 0.972 |
| 1.037 | 1.151 | 1.101 |
| 0.999 | 1.203 | 1.038 |
| 1.164 | 1.045 | 1.027 |
| 0.984 | 1.384 | 1.193 |
| 0.733 | 0.878 | 0.888 |
| 1.178 | 1.063 | 1.037 |
| 0.89 | 1.119 | 0.92 |
| 0.689 | 0.995 | 0.807 |
| 0.905 | 1.075 | 0.943 |

| | | |
|---|---|---|
| 0.712 | 0.911 | 0.866 |
| 0.753 | 0.69 | 0.766 |
| 0.936 | 0.874 | 0.886 |
| 1.047 | 1.055 | 0.929 |
| 0.657 | 1.308 | 1.09 |
| 1.118 | 1.015 | 1.108 |
| 0.936 | 0.982 | 0.916 |
| 0.641 | 0.887 | 0.727 |
| 1.099 | 1.005 | 1.097 |
| 1.09 | 1.047 | 1.252 |
| 0.975 | 1.076 | 1.057 |
| 0.89 | 1.315 | 0.957 |
| 0.907 | 0.901 | 0.858 |
| 1.087 | 1.034 | 1.036 |
| 0.887 | 1.086 | 0.975 |
| 1.25 | 1.644 | 1.182 |
| 0.943 | 1.082 | 1 |
| 1.391 | 1.901 | 1.534 |
| 0.994 | 1.181 | 1.032 |
| 0.654 | 0.948 | 0.848 |
| 0.863 | 0.948 | 0.88 |
| 1.106 | 1.208 | 0.97 |
| 1.102 | 1.247 | 1.167 |
| 0.886 | 0.962 | 1.058 |
| 0.838 | 1.584 | 0.884 |
| 0.756 | 1.072 | 0.913 |
| 1.111 | 1.061 | 1.139 |
| 0.746 | 1.386 | 0.845 |
| 1.062 | 1.23 | 0.92 |
| 100 | | 100 |
| 3.412 | 2.458 | 2.184 |
| 1.154 | 1.175 | 1.167 |
| 0.815 | 0.898 | 0.927 |
| | 100 | |
| 1.006 | 1.436 | 1.02 |
| 0.817 | 1.063 | 1.002 |
| 1.187 | 0.962 | 1.526 |
| 0.734 | 0.815 | 0.761 |
| 1.209 | 1.313 | 1.224 |
| 0.936 | 0.941 | 0.905 |
| 0.845 | 0.68 | 0.81 |
| 1.022 | 1.03 | 1.021 |

| oundance Ratio: (F1, 117) / (F1, 11 | oundance Ratio: (F1, 118) / (F1, 11 | oundance Ratio: (F1, 121) / (F1, 11 |
|---|---|---|
| 1.023 | 1.541 | 1.092 |
| 0.936 | 0.883 | 1.065 |
| 1.065 | 1.025 | 0.928 |
| 1.08 | 0.915 | 0.934 |
| 1.068 | 0.985 | 0.956 |
| 1.093 | 1.054 | 1.03 |
| 1.059 | 0.735 | 0.804 |
| 0.965 | 0.931 | 0.915 |
| 0.936 | 0.872 | 0.977 |
| 1.039 | 0.819 | 1.014 |
| 1.063 | 0.932 | 1.083 |
| 0.957 | 0.969 | 0.986 |
| 1.022 | 1.042 | 1.02 |
| 0.944 | 0.961 | 1.074 |
| 1.028 | 0.947 | 0.947 |
| 1.104 | 0.928 | 0.941 |
| 0.934 | 0.94 | 1.019 |
| 0.904 | 0.887 | 1.049 |
| 0.97 | 0.929 | 0.982 |
| 0.927 | 0.934 | 0.968 |
| 0.945 | 0.932 | 1.06 |
| 0.912 | 0.815 | 0.964 |
| 1.104 | 1.003 | 0.863 |
| 0.967 | 0.939 | 1.021 |
| 0.876 | 0.894 | 1.127 |
| 0.957 | 0.961 | 0.874 |
| 1.023 | 1.207 | 0.936 |
| 0.962 | 0.967 | 1.101 |
| 1.068 | 0.981 | 1.027 |
| 0.95 | 0.881 | 1.017 |
| 1.02 | 0.994 | 0.973 |
| 1.125 | 1.158 | 1.008 |
| 0.976 | 0.955 | 1.009 |
| 0.972 | 0.914 | 0.96 |
| 0.996 | 0.978 | 1.014 |
| 1.073 | 1.045 | 1.017 |
| 0.965 | 0.952 | 1.065 |
| 0.931 | 0.885 | 1.059 |
| 1.037 | 1.003 | 1.062 |
| 0.938 | 0.905 | 0.93 |
| 1.031 | 0.971 | 0.906 |
| 1.093 | 1.301 | 0.958 |
| 0.908 | 0.919 | 0.982 |
| 1.013 | 0.882 | 0.934 |
| 0.939 | 0.94 | 1.008 |
| 0.906 | 0.847 | 0.978 |
| 1.036 | 1.024 | 1.064 |

| | | |
|---|---|---|
| 0.959 | 0.88 | 0.904 |
| 0.962 | 0.967 | 1.035 |
| 1.044 | 0.956 | 1.108 |
| 0.901 | 0.945 | 1.136 |
| 0.921 | 0.944 | 1.012 |
| 0.915 | 0.859 | 1.172 |
| 0.947 | 0.947 | 1.007 |
| 0.833 | 0.976 | 1.064 |
| 1.006 | 0.957 | 1.014 |
| 1.055 | 0.965 | 0.963 |
| 1.08 | 0.947 | 0.911 |
| 0.786 | 1.491 | 1.916 |
| 0.924 | 0.89 | 0.942 |
| 0.958 | 0.929 | 0.966 |
| 0.914 | 0.958 | 0.8 |
| 0.912 | 0.85 | 0.759 |
| 0.952 | 0.951 | 1.005 |
| 1.029 | 0.955 | 0.895 |
| 0.947 | 0.929 | 1.212 |
| 0.399 | 1.12 | 0.384 |
| 0.928 | 0.767 | 0.966 |
| 1.051 | 1.036 | 1.014 |
| 1.086 | 0.984 | 0.642 |
| 1.016 | 0.99 | 0.956 |
| 0.728 | 0.8 | 0.982 |
| 0.994 | 0.965 | 0.925 |
| 0.951 | 1.136 | 1.077 |
| 0.892 | 1.243 | 2.078 |
| 1.114 | 1.136 | 0.917 |
| 1.024 | 0.944 | 0.878 |
| 1.173 | 1.055 | 1.177 |
| 0.86 | 1.003 | 0.896 |
| 0.976 | 1.074 | 1 |
| 0.947 | 0.946 | 0.954 |
| 0.946 | 0.955 | 0.993 |
| 0.985 | 0.881 | 0.94 |
| 1.109 | 1.169 | 0.893 |
| 0.803 | 0.737 | 0.969 |
| 0.833 | 0.964 | 1.148 |
| 0.966 | 0.902 | 1.036 |
| 0.96 | 0.977 | 1.001 |
| 1.025 | 1.056 | 1.127 |
| 0.82 | 0.887 | 0.952 |
| 1.034 | 0.889 | 1.065 |
| 0.89 | 0.87 | 1.03 |
| 1.088 | 0.95 | 1.017 |
| 0.945 | 1.027 | 1.031 |
| 0.969 | 0.88 | 0.781 |

| | | |
|---|---|---|
| 1.015 | 0.93 | 1.062 |
| 0.951 | 0.885 | 0.854 |
| 0.905 | 0.892 | 0.896 |
| 0.998 | 0.972 | 0.952 |
| 0.992 | 1.074 | 0.748 |
| 1.077 | 0.924 | 0.889 |
| 0.841 | 0.937 | 0.98 |
| 1.019 | 0.831 | 1.07 |
| 1.022 | 0.928 | 0.907 |
| 1.035 | 1.034 | 0.993 |
| 1.093 | 1.172 | 1.224 |
| 0.938 | 0.958 | 1.144 |
| 1.023 | 1.037 | 0.946 |
| 0.911 | 0.873 | 0.969 |
| 1.063 | 0.926 | 1.048 |
| 1.235 | 1.168 | 1.224 |
| 0.982 | 0.99 | 0.929 |
| 1.597 | 1.453 | 1.768 |
| 1.001 | 1.04 | 1.075 |
| 0.741 | 0.934 | 1.278 |
| 0.938 | 0.708 | 1.064 |
| 1.177 | 1.047 | 1.222 |
| 1.081 | 1.027 | 1.075 |
| 0.631 | 0.904 | 0.828 |
| 1.234 | 1.18 | 0.959 |
| 0.972 | 0.958 | 0.838 |
| 1.185 | 1.062 | 1.063 |
| 0.801 | 0.933 | 1.379 |
| 0.804 | 0.815 | 0.976 |
| 0.01 | 1.925 | 1.999 |
| 1.069 | 0.974 | 1.1 |
| 0.859 | 0.843 | 0.882 |
| 0.841 | 0.897 | 0.869 |
| 0.959 | 1.002 | 0.978 |
| 1.113 | 1.076 | 0.971 |
| 0.955 | 0.937 | 0.961 |
| 1.028 | 1.015 | 0.96 |
| 1.132 | 0.934 | 0.956 |
| 1.076 | 0.888 | 0.947 |
| 0.994 | 0.838 | 0.878 |

| bundance Ratio: (F1, 113) / (F1, 12 | bundance Ratio: (F1, 114) / (F1, 12 | bundance Ratio: (F1, 115) / (F1, 12 |
|---|---|---|
| 0.803 | 0.878 | 0.791 |
| 1.095 | 0.795 | 1.084 |
| 1.047 | 1.339 | 1.091 |
| 1.225 | 1.377 | 1.135 |
| 1.331 | 1.368 | 1.256 |
| 0.982 | 1.321 | 0.89 |
| 0.991 | 1.038 | 1.092 |
| 1.025 | 0.861 | 1.03 |
| 0.909 | 0.777 | 0.995 |
| 0.91 | 0.877 | 0.635 |
| 1.089 | 1.352 | 1.015 |
| 0.948 | 0.879 | 1.031 |
| 1.222 | 1.151 | 1.249 |
| 0.787 | 0.787 | 0.866 |
| 1.441 | 1.409 | 1.312 |
| 1.048 | 1.016 | 1.192 |
| 0.949 | 0.845 | 1.043 |
| 0.822 | 0.69 | 0.985 |
| 0.896 | 0.839 | 0.964 |
| 0.893 | 0.729 | 0.998 |
| 0.853 | 0.737 | 1.052 |
| 0.666 | 0.603 | 0.727 |
| 1.431 | 1.545 | 1.199 |
| 0.873 | 0.841 | 0.997 |
| 0.712 | 0.616 | 0.88 |
| 1.022 | 0.868 | 1.132 |
| 1.039 | 0.982 | 1.092 |
| 0.925 | 0.88 | 0.93 |
| 1.176 | 1.182 | 1.112 |
| 1.047 | 0.974 | 0.959 |
| 1.155 | 1.208 | 1.115 |
| 0.992 | 0.96 | 0.972 |
| 1.151 | 0.904 | 1.08 |
| 1.222 | 1.116 | 1.178 |
| 1.054 | 1.074 | 1.088 |
| 0.919 | 0.977 | 0.864 |
| 0.821 | 0.772 | 0.989 |
| 0.849 | 0.719 | 0.956 |
| 0.957 | 0.83 | 0.993 |
| 0.852 | 0.795 | 1.001 |
| 0.901 | 1.055 | 0.935 |
| 0.956 | 0.939 | 1.078 |
| 0.82 | 0.864 | 0.868 |
| 1.148 | 1.046 | 1.101 |
| 1.138 | 1.052 | 1.075 |
| 0.837 | 0.826 | 0.993 |
| 1.045 | 0.946 | 1.062 |

| | | |
|---|---|---|
| 1.068 | 1.238 | 1.119 |
| 0.946 | 0.835 | 1.018 |
| 1.055 | 0.891 | 1.172 |
| 0.816 | 0.664 | 0.855 |
| 1.018 | 0.823 | 1.011 |
| 1.222 | 0.839 | 1.086 |
| 1.128 | 0.925 | 1.134 |
| 0.786 | 0.617 | 0.939 |
| 0.889 | 0.767 | 0.983 |
| 1.124 | 1.112 | 1.067 |
| 1.095 | 1.193 | 1.058 |
| 1.582 | 0.638 | 1.431 |
| 0.915 | 0.976 | 0.862 |
| 1.017 | 1.015 | 1.098 |
| 1.968 | 1.312 | 1.414 |
| 1.288 | 1.248 | 1.168 |
| 0.855 | 0.76 | 0.968 |
| 1.144 | 1.117 | 1.198 |
| 0.744 | 0.757 | 0.819 |
| 3.789 | 1.686 | 3.829 |
| 0.958 | 0.862 | 0.95 |
| 1.033 | 1.037 | 1.056 |
| 1.141 | 1.278 | 1.72 |
| 0.969 | 1.041 | 1.015 |
| 1.379 | 1.566 | 1.327 |
| 1.264 | 1.284 | 1.224 |
| 0.945 | 0.815 | 0.874 |
| 0.708 | 0.608 | 0.553 |
| 1.187 | 1.204 | 1.062 |
| 1.541 | 1.317 | 1.181 |
| 1.361 | 0.879 | 1.046 |
| 1.39 | 1.104 | 1.302 |
| 1.045 | 0.905 | 1.072 |
| 1.13 | 0.92 | 1.168 |
| 1.034 | 0.921 | 0.93 |
| 0.985 | 0.878 | 1.072 |
| 1.152 | 1.259 | 0.98 |
| 0.86 | 0.87 | 0.926 |
| 1.227 | 0.887 | 1.15 |
| 1.032 | 1.001 | 1.111 |
| 1.204 | 0.998 | 1.202 |
| 0.956 | 1.033 | 0.927 |
| 1.112 | 1.034 | 1.454 |
| 0.796 | 0.689 | 0.824 |
| 1.124 | 1.143 | 1.032 |
| 0.966 | 0.875 | 1.1 |
| 0.891 | 0.669 | 0.965 |
| 1.156 | 1.158 | 1.376 |

| | | |
|---|---|---|
| 0.731 | 0.67 | 0.858 |
| 0.714 | 0.882 | 0.808 |
| 1.103 | 1.044 | 0.975 |
| 1.114 | 1.1 | 1.108 |
| 0.673 | 0.878 | 1.748 |
| 1.192 | 1.258 | 1.142 |
| 1.381 | 0.955 | 1.002 |
| 0.779 | 0.599 | 0.829 |
| 1.175 | 1.211 | 1.107 |
| 1.062 | 1.098 | 1.054 |
| 0.763 | 0.797 | 0.879 |
| 1.376 | 0.778 | 1.149 |
| 1.12 | 0.958 | 0.953 |
| 1.1 | 1.123 | 1.068 |
| 0.981 | 0.846 | 1.037 |
| 0.748 | 1.021 | 1.343 |
| 1.041 | 1.014 | 1.164 |
| 1.04 | 0.787 | 1.075 |
| 1.027 | 0.925 | 1.099 |
| 0.739 | 0.512 | 0.742 |
| 0.925 | 0.812 | 0.891 |
| 1.078 | 0.905 | 0.989 |
| 1.028 | 1.025 | 1.16 |
| 1.152 | 1.069 | 1.162 |
| 1.571 | 0.874 | 1.652 |
| 1.291 | 0.902 | 1.279 |
| 0.929 | 1.044 | 0.998 |
| 0.897 | 0.541 | 1.005 |
| 1.294 | 1.089 | 1.26 |
| | 100 | |
| 0.639 | 1.707 | 1.23 |
| 1.063 | 1.049 | 1.068 |
| 1.043 | 0.923 | 1.018 |
| | | 100 |
| 1.588 | 1.157 | 1.652 |
| 1.115 | 0.835 | 1.086 |
| 1.119 | 1.223 | 0.991 |
| 0.847 | 0.763 | 0.848 |
| 1.361 | 1.26 | 1.368 |
| 1.076 | 0.979 | 0.984 |
| 0.734 | 0.893 | 0.718 |
| 1.145 | 1.164 | 1.174 |

| bundance Ratio: (F1, 116) / (F1, 12 | bundance Ratio: (F1, 117) / (F1, 12 | bundance Ratio: (F1, 118) / (F1, 12 |
|---|---|---|
| 0.876 | 0.937 | 1.411 |
| 0.953 | 0.878 | 0.829 |
| 1.246 | 1.147 | 1.105 |
| 1.221 | 1.156 | 0.98 |
| 1.4 | 1.118 | 1.03 |
| 1.144 | 1.062 | 1.024 |
| 0.999 | 1.317 | 0.914 |
| 0.969 | 1.055 | 1.018 |
| 0.841 | 0.958 | 0.893 |
| 0.728 | 1.025 | 0.808 |
| 1.037 | 0.981 | 0.86 |
| 0.947 | 0.971 | 0.983 |
| 1.172 | 1.002 | 1.021 |
| 0.821 | 0.879 | 0.894 |
| 1.473 | 1.086 | 1.001 |
| 1.078 | 1.173 | 0.986 |
| 0.884 | 0.917 | 0.922 |
| 0.809 | 0.861 | 0.845 |
| 0.92 | 0.988 | 0.946 |
| 0.848 | 0.958 | 0.965 |
| 0.892 | 0.891 | 0.879 |
| 0.665 | 0.946 | 0.845 |
| 1.446 | 1.28 | 1.163 |
| 0.906 | 0.948 | 0.919 |
| 0.701 | 0.777 | 0.794 |
| 1.004 | 1.095 | 1.1 |
| 0.994 | 1.093 | 1.289 |
| 0.88 | 0.873 | 0.878 |
| 1.206 | 1.04 | 0.956 |
| 1.007 | 0.935 | 0.866 |
| 1.136 | 1.048 | 1.021 |
| 0.949 | 1.116 | 1.149 |
| 0.916 | 0.967 | 0.946 |
| 1.219 | 1.013 | 0.952 |
| 1.077 | 0.982 | 0.964 |
| 0.985 | 1.055 | 1.027 |
| 0.841 | 0.907 | 0.894 |
| 0.797 | 0.879 | 0.836 |
| 0.93 | 0.976 | 0.944 |
| 0.925 | 1.008 | 0.973 |
| 1.051 | 1.137 | 1.072 |
| 1.021 | 1.14 | 1.357 |
| 0.878 | 0.925 | 0.935 |
| 0.967 | 1.084 | 0.944 |
| 1.042 | 0.931 | 0.932 |
| 0.876 | 0.927 | 0.867 |
| 0.94 | 0.974 | 0.963 |

| | | |
|---|---|---|
| 1.262 | 1.061 | 0.974 |
| 0.955 | 0.93 | 0.935 |
| 1.051 | 0.942 | 0.862 |
| 0.678 | 0.794 | 0.832 |
| 0.831 | 0.91 | 0.932 |
| 0.85 | 0.781 | 0.733 |
| 0.988 | 0.941 | 0.941 |
| 0.774 | 0.783 | 0.917 |
| 0.903 | 0.993 | 0.944 |
| 1.066 | 1.095 | 1.001 |
| 1.044 | 1.185 | 1.04 |
| 0.714 | 0.41 | 0.778 |
| 0.938 | 0.981 | 0.945 |
| 0.926 | 0.991 | 0.962 |
| 1.275 | 1.142 | 1.196 |
| 0.987 | 1.202 | 1.12 |
| 0.898 | 0.948 | 0.946 |
| 1.107 | 1.149 | 1.067 |
| 0.773 | 0.781 | 0.767 |
| 1.898 | 1.038 | 2.915 |
| 0.932 | 0.961 | 0.794 |
| 1.02 | 1.036 | 1.022 |
| 1.247 | 1.69 | 1.532 |
| 1.037 | 1.063 | 1.036 |
| 1.522 | 0.741 | 0.815 |
| 1.274 | 1.074 | 1.043 |
| 0.887 | 0.882 | 1.054 |
| 0.469 | 0.429 | 0.598 |
| 1.091 | 1.215 | 1.239 |
| 1.152 | 1.166 | 1.075 |
| 0.682 | 0.997 | 0.896 |
| 1.13 | 0.96 | 1.119 |
| 1.011 | 0.976 | 1.074 |
| 0.925 | 0.992 | 0.992 |
| 0.966 | 0.953 | 0.962 |
| 0.979 | 1.048 | 0.937 |
| 1.027 | 1.241 | 1.308 |
| 0.892 | 0.828 | 0.761 |
| 0.846 | 0.726 | 0.84 |
| 1.062 | 0.933 | 0.871 |
| 1.038 | 0.96 | 0.976 |
| 0.911 | 0.91 | 0.937 |
| 1.253 | 0.861 | 0.932 |
| 0.834 | 0.971 | 0.835 |
| 1.006 | 0.863 | 0.845 |
| 0.904 | 1.07 | 0.934 |
| 0.783 | 0.917 | 0.997 |
| 1.207 | 1.24 | 1.127 |

| | | |
|---|---|---|
| 0.815 | 0.956 | 0.876 |
| 0.897 | 1.114 | 1.036 |
| 0.989 | 1.01 | 0.995 |
| 0.976 | 1.049 | 1.021 |
| 1.457 | 1.326 | 1.435 |
| 1.247 | 1.212 | 1.04 |
| 0.934 | 0.858 | 0.956 |
| 0.68 | 0.952 | 0.777 |
| 1.209 | 1.126 | 1.022 |
| 1.261 | 1.043 | 1.042 |
| 0.864 | 0.893 | 0.957 |
| 0.837 | 0.82 | 0.837 |
| 0.907 | 1.081 | 1.096 |
| 1.07 | 0.941 | 0.901 |
| 0.931 | 1.015 | 0.884 |
| 0.965 | 1.008 | 0.954 |
| 1.076 | 1.057 | 1.065 |
| 0.868 | 0.904 | 0.822 |
| 0.96 | 0.931 | 0.967 |
| 0.664 | 0.58 | 0.731 |
| 0.828 | 0.882 | 0.666 |
| 0.794 | 0.964 | 0.857 |
| 1.086 | 1.005 | 0.955 |
| 1.277 | 0.761 | 1.092 |
| 0.921 | 1.287 | 1.23 |
| 1.089 | 1.16 | 1.144 |
| 1.071 | 1.114 | 0.999 |
| 0.613 | 0.581 | 0.676 |
| 0.943 | 0.824 | 0.835 |
| 100 | | |
| 1.093 | 0.01 | 0.963 |
| 1.061 | 0.972 | 0.885 |
| 1.05 | 0.974 | 0.955 |
| 1.174 | 0.968 | 1.032 |
| 1.024 | 0.98 | 1.024 |
| 1.572 | 1.146 | 1.109 |
| 0.792 | 0.993 | 0.975 |
| 1.275 | 1.071 | 1.058 |
| 0.947 | 1.185 | 0.978 |
| 0.856 | 1.137 | 0.939 |
| 1.163 | 1.132 | 0.955 |

| oundance Ratio: (F1, 119) / (F1, 12 | oundances (Scaled): F1: 113, Contr | oundances (Scaled): F1: 114, Samp |
|---|---|---|
| 0.916 | 84.4 | 92.3 |
| 0.939 | 115.7 | 84 |
| 1.078 | 92.5 | 118.3 |
| 1.071 | 106.9 | 120.2 |
| 1.046 | 111.5 | 114.6 |
| 0.971 | 93.6 | 125.9 |
| 1.243 | 92.3 | 96.6 |
| 1.093 | 101.8 | 85.5 |
| 1.024 | 98.3 | 84.1 |
| 0.986 | 104.4 | 100.7 |
| 0.923 | 105.5 | 131 |
| 1.014 | 97.5 | 90.5 |
| 0.981 | 111.1 | 104.6 |
| 0.931 | 90.4 | 90.4 |
| 1.056 | 117.9 | 115.3 |
| 1.062 | 98 | 95 |
| 0.982 | 100.6 | 89.6 |
| 0.953 | 94.4 | 79.2 |
| 1.018 | 94.7 | 88.6 |
| 1.034 | 96.2 | 78.6 |
| 0.944 | 94.1 | 81.4 |
| 1.037 | 82.2 | 74.3 |
| 1.159 | 112 | 120.9 |
| 0.98 | 93.6 | 90.1 |
| 0.887 | 89.5 | 77.3 |
| 1.144 | 97.7 | 83 |
| 1.068 | 97.2 | 91.8 |
| 0.908 | 101.8 | 96.8 |
| 0.974 | 108.8 | 109.4 |
| 0.984 | 107.8 | 100.3 |
| 1.027 | 106 | 111 |
| 0.992 | 97.6 | 94.5 |
| 0.991 | 115.7 | 90.9 |
| 1.042 | 111.8 | 102.2 |
| 0.986 | 102.5 | 104.5 |
| 0.983 | 94.2 | 100 |
| 0.939 | 91.7 | 86.2 |
| 0.944 | 97.3 | 82.4 |
| 0.942 | 101.1 | 87.7 |
| 1.075 | 89.3 | 83.3 |
| 1.103 | 87.3 | 102.3 |
| 1.044 | 89.6 | 88 |
| 1.018 | 89.8 | 94.6 |
| 1.07 | 109.9 | 100.1 |
| 0.992 | 111.6 | 103.1 |
| 1.023 | 91.1 | 89.9 |
| 0.94 | 106.2 | 96.2 |

| | | |
|---|---|---|
| 1.106 | 96.8 | 112.2 |
| 0.967 | 99.8 | 88 |
| 0.902 | 107.2 | 90.5 |
| 0.881 | 100.1 | 81.5 |
| 0.988 | 108.4 | 87.6 |
| 0.853 | 132.8 | 91.2 |
| 0.993 | 112.1 | 91.9 |
| 0.94 | 93.1 | 73 |
| 0.986 | 95.3 | 82.2 |
| 1.038 | 105.7 | 104.6 |
| 1.098 | 100.5 | 109.6 |
| 0.522 | 178.9 | 72.1 |
| 1.061 | 95.3 | 101.7 |
| 1.035 | 101.2 | 100.9 |
| 1.249 | 149.1 | 99.4 |
| 1.318 | 110.4 | 106.9 |
| 0.995 | 92.8 | 82.5 |
| 1.117 | 102.8 | 100.4 |
| 0.825 | 92.1 | 93.6 |
| 2.604 | 161.6 | 71.9 |
| 1.035 | 102.3 | 92 |
| 0.986 | 100.9 | 101.3 |
| 1.557 | 81.7 | 91.6 |
| 1.046 | 94.5 | 101.4 |
| 1.019 | 117.7 | 133.7 |
| 1.081 | 109.4 | 111.1 |
| 0.928 | 102.3 | 88.3 |
| 0.481 | 116.9 | 100.4 |
| 1.091 | 104.5 | 105.9 |
| 1.139 | 128.8 | 110.1 |
| 0.849 | 141.3 | 91.2 |
| 1.116 | 121.9 | 96.9 |
| 1 | 103.5 | 89.6 |
| 1.048 | 110.6 | 90.1 |
| 1.007 | 106.5 | 94.8 |
| 1.064 | 98.9 | 88.2 |
| 1.119 | 101.4 | 110.9 |
| 1.032 | 96 | 97.1 |
| 0.871 | 130.1 | 94 |
| 0.965 | 103.5 | 100.4 |
| 0.999 | 115 | 95.3 |
| 0.887 | 101.1 | 109.3 |
| 1.05 | 102.3 | 95.1 |
| 0.939 | 92.5 | 80 |
| 0.97 | 112.6 | 114.5 |
| 0.983 | 98.7 | 89.4 |
| 0.97 | 99.1 | 74.4 |
| 1.28 | 96.9 | 97.1 |

| | | |
|---|---|---|
| 0.942 | 85.4 | 78.3 |
| 1.171 | 74.9 | 92.6 |
| 1.116 | 107.2 | 101.5 |
| 1.05 | 105.8 | 104.5 |
| 1.336 | 54.7 | 71.2 |
| 1.125 | 103.4 | 109.2 |
| 1.02 | 136.3 | 94.2 |
| 0.934 | 95.2 | 73.1 |
| 1.102 | 105 | 108.2 |
| 1.008 | 99.2 | 102.5 |
| 0.817 | 87.6 | 91.5 |
| 0.874 | 143.5 | 81.2 |
| 1.057 | 109.7 | 93.8 |
| 1.032 | 106.9 | 109.1 |
| 0.954 | 102.6 | 88.5 |
| 0.817 | 76.2 | 104 |
| 1.076 | 98.1 | 95.5 |
| 0.566 | 117.9 | 89.2 |
| 0.931 | 104.8 | 94.4 |
| 0.783 | 102.8 | 71.2 |
| 0.94 | 106.5 | 93.5 |
| 0.819 | 116.5 | 97.8 |
| 0.93 | 100.4 | 100.2 |
| 1.208 | 105.7 | 98.1 |
| 1.043 | 131.2 | 73 |
| 1.193 | 114 | 79.7 |
| 0.94 | 91.8 | 103.2 |
| 0.725 | 118.8 | 71.6 |
| 1.025 | 125.2 | 105.3 |
| | | 367.1 |
| 0.5 | 71.6 | 191.5 |
| 0.909 | 106.2 | 104.8 |
| 1.133 | 103.1 | 91.2 |
| 1.151 | 130.6 | 95.2 |
| 1.022 | 110.3 | 82.6 |
| 1.03 | 97.4 | 106.5 |
| 1.04 | 93.4 | 84.1 |
| 1.042 | 115.4 | 106.8 |
| 1.046 | 105 | 95.6 |
| 1.056 | 80 | 97.4 |
| 1.139 | 103.3 | 105 |

| oundances (Scaled): F1: 115, Samp | oundances (Scaled): F1: 116, Samp | oundances (Scaled): F1: 117, Samp |
|---|---|---|
| 83.1 | 92 | 98.4 |
| 114.5 | 100.7 | 92.8 |
| 96.4 | 110.1 | 101.4 |
| 99.1 | 106.6 | 100.9 |
| 105.2 | 117.3 | 93.7 |
| 84.8 | 109 | 101.2 |
| 101.6 | 93 | 122.6 |
| 102.4 | 96.3 | 104.8 |
| 107.6 | 90.9 | 103.6 |
| 72.9 | 83.6 | 117.6 |
| 98.3 | 100.5 | 95 |
| 106.1 | 97.5 | 99.9 |
| 113.6 | 106.6 | 91.1 |
| 99.4 | 94.3 | 100.9 |
| 107.4 | 120.5 | 88.8 |
| 111.5 | 100.8 | 109.7 |
| 110.7 | 93.8 | 97.3 |
| 113.1 | 92.9 | 98.9 |
| 101.9 | 97.2 | 104.4 |
| 107.5 | 91.4 | 103.2 |
| 116.1 | 98.5 | 98.4 |
| 89.6 | 81.9 | 116.7 |
| 93.8 | 113.2 | 100.1 |
| 106.9 | 97.1 | 101.6 |
| 110.6 | 88.1 | 97.7 |
| 108.3 | 96 | 104.7 |
| 102.1 | 92.9 | 102.2 |
| 102.2 | 96.8 | 96.1 |
| 102.9 | 111.5 | 96.2 |
| 98.7 | 103.6 | 96.2 |
| 102.4 | 104.4 | 96.2 |
| 95.6 | 93.4 | 109.8 |
| 108.6 | 92.1 | 97.2 |
| 107.8 | 111.6 | 92.7 |
| 105.8 | 104.7 | 95.5 |
| 88.5 | 100.9 | 108.1 |
| 110.5 | 93.9 | 101.3 |
| 109.6 | 91.4 | 100.7 |
| 104.9 | 98.2 | 103.1 |
| 104.9 | 97 | 105.7 |
| 90.6 | 101.9 | 110.2 |
| 101.1 | 95.7 | 106.9 |
| 95 | 96.1 | 101.2 |
| 105.4 | 92.5 | 103.7 |
| 105.4 | 102.2 | 91.3 |
| 108.1 | 95.4 | 100.9 |
| 108 | 95.6 | 99 |

| | | |
|---|---|---|
| 101.4 | 114.4 | 96.2 |
| 107.4 | 100.7 | 98 |
| 119.1 | 106.8 | 95.7 |
| 104.9 | 83.2 | 97.4 |
| 107.6 | 88.4 | 96.9 |
| 118 | 92.3 | 84.8 |
| 112.7 | 98.2 | 93.5 |
| 111.2 | 91.7 | 92.7 |
| 105.4 | 96.8 | 106.4 |
| 100.4 | 100.3 | 103 |
| 97.1 | 95.9 | 108.8 |
| 161.8 | 80.7 | 46.4 |
| 89.9 | 97.7 | 102.2 |
| 109.2 | 92.1 | 98.6 |
| 107.2 | 96.6 | 86.5 |
| 100.2 | 84.6 | 103.1 |
| 105 | 97.5 | 102.9 |
| 107.7 | 99.5 | 103.3 |
| 101.3 | 95.6 | 96.7 |
| 163.3 | 81 | 44.3 |
| 101.4 | 99.6 | 102.6 |
| 103.2 | 99.6 | 101.2 |
| 123.2 | 89.4 | 121.1 |
| 98.9 | 101.1 | 103.6 |
| 113.3 | 130 | 63.3 |
| 105.9 | 110.2 | 92.9 |
| 94.7 | 96.1 | 95.6 |
| 91.3 | 77.3 | 70.8 |
| 93.5 | 96 | 107 |
| 98.7 | 96.3 | 97.5 |
| 108.5 | 70.7 | 103.4 |
| 114.2 | 99.1 | 84.2 |
| 106.1 | 100.1 | 96.6 |
| 114.3 | 90.5 | 97.1 |
| 95.7 | 99.4 | 98.1 |
| 107.7 | 98.4 | 105.3 |
| 86.3 | 90.4 | 109.3 |
| 103.4 | 99.5 | 92.4 |
| 121.9 | 89.7 | 76.9 |
| 111.4 | 106.6 | 93.6 |
| 114.8 | 99.1 | 91.7 |
| 98.1 | 96.4 | 96.3 |
| 133.8 | 115.3 | 79.2 |
| 95.7 | 96.8 | 112.8 |
| 103.4 | 100.9 | 86.5 |
| 112.3 | 92.4 | 109.2 |
| 107.4 | 87.1 | 102 |
| 115.3 | 101.2 | 104 |

| | | |
|---|---|---|
| 100.2 | 95.2 | 111.6 |
| 84.8 | 94.1 | 116.9 |
| 94.8 | 96.1 | 98.2 |
| 105.3 | 92.7 | 99.7 |
| 141.9 | 118.3 | 107.7 |
| 99.1 | 108.2 | 105.2 |
| 98.9 | 92.2 | 84.6 |
| 101.2 | 83 | 116.3 |
| 99 | 108 | 100.6 |
| 98.4 | 117.8 | 97.4 |
| 100.9 | 99.1 | 102.5 |
| 119.8 | 87.3 | 85.5 |
| 93.3 | 88.8 | 105.8 |
| 103.7 | 103.9 | 91.4 |
| 108.4 | 97.3 | 106.2 |
| 136.7 | 98.3 | 102.7 |
| 109.6 | 101.4 | 99.5 |
| 121.8 | 98.3 | 102.4 |
| 112.1 | 98 | 95 |
| 103.2 | 92.4 | 80.7 |
| 102.6 | 95.4 | 101.6 |
| 106.8 | 85.8 | 104.1 |
| 113.3 | 106.1 | 98.2 |
| 106.6 | 117.2 | 69.8 |
| 137.9 | 77 | 107.5 |
| 112.9 | 96.2 | 102.4 |
| 98.6 | 105.9 | 110.1 |
| 133.1 | 81.2 | 77 |
| 121.9 | 91.2 | 79.7 |
| | 432.9 | |
| 137.9 | 122.6 | |
| 106.7 | 106 | 97.1 |
| 100.6 | 103.8 | 96.2 |
| 800 | | |
| 136 | 96.6 | 79.6 |
| 107.5 | 101.3 | 97 |
| 86.3 | 136.8 | 99.8 |
| 93.5 | 87.3 | 109.5 |
| 116 | 108.1 | 90.8 |
| 96.1 | 92.4 | 115.7 |
| 78.3 | 93.4 | 124 |
| 105.8 | 104.9 | 102.1 |

| oundances (Scaled): F1: 118, Samp | oundances (Scaled): F1: 119, Samp | oundances (Scaled): F1: 121, Samp |
|---|---|---|
| 148.3 | 96.3 | 105.1 |
| 87.5 | 99.2 | 105.7 |
| 97.6 | 95.2 | 88.4 |
| 85.5 | 93.5 | 87.3 |
| 86.3 | 87.7 | 83.8 |
| 97.6 | 92.6 | 95.3 |
| 85.1 | 115.7 | 93.1 |
| 101.2 | 108.6 | 99.4 |
| 96.6 | 110.8 | 108.2 |
| 92.7 | 113.2 | 114.8 |
| 83.4 | 89.4 | 96.9 |
| 101.2 | 104.4 | 102.9 |
| 92.9 | 89.2 | 90.9 |
| 102.8 | 106.9 | 114.9 |
| 81.9 | 86.4 | 81.8 |
| 92.2 | 99.3 | 93.5 |
| 97.8 | 104.1 | 106.1 |
| 97.1 | 109.5 | 114.9 |
| 99.9 | 107.6 | 105.7 |
| 104 | 111.4 | 107.8 |
| 97 | 104.1 | 110.4 |
| 104.2 | 127.9 | 123.3 |
| 91 | 90.7 | 78.2 |
| 98.5 | 105 | 107.2 |
| 99.7 | 111.5 | 125.7 |
| 105.2 | 109.4 | 95.6 |
| 120.5 | 99.9 | 93.5 |
| 96.5 | 99.9 | 110 |
| 88.4 | 90.1 | 92.5 |
| 89.2 | 101.3 | 102.9 |
| 93.8 | 94.4 | 91.9 |
| 113 | 97.6 | 98.4 |
| 95.1 | 99.6 | 100.6 |
| 87.1 | 95.3 | 91.5 |
| 93.8 | 95.9 | 97.3 |
| 105.2 | 100.7 | 102.4 |
| 99.9 | 104.9 | 111.7 |
| 95.8 | 108.2 | 114.6 |
| 99.8 | 99.5 | 105.7 |
| 102 | 112.7 | 104.9 |
| 103.9 | 106.9 | 96.9 |
| 127.2 | 97.8 | 93.7 |
| 102.4 | 111.4 | 109.5 |
| 90.3 | 102.4 | 95.7 |
| 91.3 | 97.2 | 98 |
| 94.3 | 111.4 | 108.9 |
| 97.9 | 95.5 | 101.7 |

| | | |
|---|---|---|
| 88.2 | 100.3 | 90.6 |
| 98.6 | 101.9 | 105.5 |
| 87.6 | 91.6 | 101.6 |
| 102.1 | 108.1 | 122.7 |
| 99.3 | 105.2 | 106.5 |
| 79.6 | 92.7 | 108.6 |
| 93.5 | 98.7 | 99.4 |
| 108.6 | 111.3 | 118.4 |
| 101.2 | 105.7 | 107.2 |
| 94.2 | 97.7 | 94.1 |
| 95.5 | 100.8 | 91.8 |
| 88 | 59 | 113.1 |
| 98.4 | 110.6 | 104.2 |
| 95.6 | 103 | 99.5 |
| 90.7 | 94.7 | 75.8 |
| 96 | 113 | 85.7 |
| 102.7 | 108 | 108.6 |
| 95.9 | 100.4 | 89.9 |
| 94.9 | 102.1 | 123.7 |
| 124.3 | 111 | 42.6 |
| 84.8 | 110.5 | 106.8 |
| 99.8 | 96.3 | 97.7 |
| 109.8 | 111.6 | 71.7 |
| 101 | 102 | 97.5 |
| 69.6 | 87 | 85.4 |
| 90.3 | 93.5 | 86.5 |
| 114.2 | 100.5 | 108.3 |
| 98.7 | 79.4 | 165 |
| 109 | 96 | 88 |
| 89.9 | 95.2 | 83.6 |
| 93 | 88.1 | 103.8 |
| 98.2 | 97.9 | 87.7 |
| 106.3 | 99 | 99 |
| 97.1 | 102.6 | 97.9 |
| 99 | 103.6 | 102.9 |
| 94.1 | 106.9 | 100.5 |
| 115.2 | 98.6 | 88 |
| 84.9 | 115.2 | 111.6 |
| 89 | 92.3 | 106 |
| 87.3 | 96.8 | 100.3 |
| 93.2 | 95.4 | 95.5 |
| 99.1 | 93.9 | 105.8 |
| 85.7 | 96.6 | 92 |
| 96.9 | 109.1 | 116.1 |
| 84.6 | 97.2 | 100.2 |
| 95.4 | 100.4 | 102.1 |
| 110.9 | 107.9 | 111.2 |
| 94.5 | 107.3 | 83.8 |

| | | |
|---|---|---|
| 102.3 | 110 | 116.8 |
| 108.7 | 122.9 | 104.9 |
| 96.7 | 108.5 | 97.2 |
| 97 | 99.8 | 95 |
| 116.5 | 108.5 | 81.2 |
| 90.3 | 97.7 | 86.8 |
| 94.3 | 100.7 | 98.7 |
| 94.9 | 114.1 | 122.2 |
| 91.4 | 98.5 | 89.4 |
| 97.3 | 94.1 | 93.4 |
| 109.9 | 93.8 | 114.8 |
| 87.3 | 91.2 | 104.3 |
| 107.3 | 103.5 | 97.9 |
| 87.6 | 100.3 | 97.2 |
| 92.5 | 99.8 | 104.6 |
| 97.1 | 83.2 | 101.8 |
| 100.3 | 101.3 | 94.2 |
| 93.1 | 64.1 | 113.3 |
| 98.7 | 94.9 | 102 |
| 101.7 | 108.9 | 139.1 |
| 76.7 | 108.3 | 115.2 |
| 92.6 | 88.4 | 108 |
| 93.3 | 90.9 | 97.7 |
| 100.1 | 110.8 | 91.7 |
| 102.8 | 87.1 | 83.5 |
| 101 | 105.4 | 88.3 |
| 98.7 | 92.9 | 98.8 |
| 89.6 | 96.1 | 132.5 |
| 80.8 | 99.1 | 96.7 |
| 108 | 56.1 | 112.2 |
| 88.4 | 90.8 | 99.9 |
| 94.3 | 112 | 98.8 |
| 85 | 94.7 | 82.3 |
| 101.3 | 101.1 | 99 |
| 96.5 | 89.7 | 87 |
| 107.5 | 114.6 | 110.2 |
| 89.7 | 88.4 | 84.8 |
| 95.4 | 102.1 | 97.6 |
| 102.4 | 115.3 | 109.1 |
| 86.1 | 102.7 | 90.2 |

| emPAI | # Razor Peptides | Score Sequest HT | # Peptides Sequest HT |
|---|---|---|---|
| 23.119 | 25 | 302.99 | 29 |
| 32.246 | 18 | 254.75 | 16 |
| 718.686 | 0 | 200.79 | 10 |
| 230.013 | 18 | 170.11 | 15 |
| 2.594 | 0 | 162.47 | 17 |
| 186.382 | 2 | 161.4 | 14 |
| 718.686 | 0 | 135.25 | 10 |
| 3.062 | 1 | 129.68 | 10 |
| 3.062 | 11 | 124.68 | 10 |
| 2.008 | 0 | 107.7 | 7 |
| 73.989 | 0 | 107.2 | 7 |
| 0.526 | 0 | 66.49 | 7 |
| 0.896 | 0 | 65.26 | 3 |
| 0.708 | 5 | 63.24 | 8 |
| 24.119 | 0 | 52.99 | 10 |
| 0.382 | 0 | 51.65 | 3 |
| 0.409 | 0 | 40.1 | 4 |
| 1.395 | 0 | 39.2 | 10 |
| 0.354 | 0 | 38.95 | 5 |
| 0.487 | 0 | 34.81 | 4 |
| 2.415 | 0 | 32.36 | 6 |
| 1.424 | 0 | 31.65 | 3 |
| 0.431 | 3 | 31.37 | 5 |
| 0.271 | 0 | 30.7 | 5 |
| 0.931 | 0 | 29.44 | 5 |
| 0.995 | 0 | 22.17 | 2 |
| 0.389 | 0 | 21.43 | 4 |
| 0.638 | 0 | 20.58 | 2 |
| 0.445 | 0 | 20.39 | 3 |
| 0.252 | 0 | 20.26 | 3 |
| 0.233 | 0 | 19.3 | 4 |
| 0.25 | 0 | 18.83 | 3 |
| 0.389 | 0 | 17.33 | 1 |
| 0.233 | 0 | 16.95 | 3 |
| 0.311 | 0 | 15.04 | 3 |
| 0.145 | 0 | 15 | 1 |
| 0.259 | 0 | 14.13 | 2 |
| 0.931 | 0 | 12.53 | 4 |
| 0.186 | 0 | 12.08 | 4 |
| 0.73 | 0 | 11.26 | 5 |
| 1.031 | 0 | 10.84 | 4 |
| 0.212 | 0 | 10.61 | 2 |
| 0.101 | 0 | 10.43 | 1 |
| 0.638 | 0 | 10.03 | 2 |
| 0.134 | 0 | 10 | 3 |
| 0.468 | 0 | 9.44 | 1 |
| 0.701 | 0 | 8.48 | 2 |

| | | | |
|---|---|---|---|
| 0.136 | 0 | 8.25 | 1 |
| 1.154 | 0 | 8.16 | 1 |
| 0.389 | 0 | 8.05 | 2 |
| 1.154 | 0 | 8 | 1 |
| 0.369 | 0 | 7.88 | 2 |
| 0.133 | 0 | 7.77 | 2 |
| 0.359 | 0 | 7.7 | 2 |
| 0.122 | 0 | 7.39 | 1 |
| 0.52 | 0 | 7.03 | 2 |
| 0.125 | 0 | 6.94 | 1 |
| 0.202 | 0 | 6.91 | 2 |
| 0.129 | 0 | 6.89 | 1 |
| 0.15 | 0 | 6.73 | 2 |
| 0.668 | 0 | 6.43 | 1 |
| 0.05 | 0 | 6.23 | 1 |
| 0.05 | 0 | 5.9 | 1 |
| 0.245 | 0 | 5.78 | 2 |
| 0.212 | 0 | 5.73 | 1 |
| 0.166 | 0 | 5.7 | 2 |
| 0.129 | 0 | 5.63 | 1 |
| 0.066 | 0 | 5.6 | 1 |
| 0.304 | 0 | 5.53 | 3 |
| 0.129 | 0 | 5.49 | 1 |
| 0.017 | 0 | 5.19 | 3 |
| 0.292 | 0 | 4.74 | 1 |
| 0.311 | 0 | 4.68 | 2 |
| 0.015 | 0 | 4.67 | 1 |
| 0.212 | 0 | 4.48 | 1 |
| 0.233 | 0 | 4.39 | 1 |
| 0.105 | 0 | 4.33 | 1 |
| 0.166 | 0 | 4.21 | 1 |
| 0.058 | 0 | 4.1 | 1 |
| 0.184 | 1 | 3.93 | 3 |
| 0.086 | 0 | 3.82 | 1 |
| 0.166 | 0 | 3.76 | 1 |
| 0.194 | 0 | 3.74 | 1 |
| 0.425 | 0 | 3.66 | 2 |
| 0.058 | 0 | 3.54 | 1 |
| 0.194 | 0 | 3.51 | 1 |
| 0.105 | 0 | 3.51 | 1 |
| 0.103 | 0 | 3.43 | 2 |
| 0.046 | 0 | 3.41 | 1 |
| 0.061 | 0 | 3.4 | 1 |
| 0.136 | 0 | 3.34 | 1 |
| 0.066 | 0 | 3.29 | 1 |
| 0.52 | 0 | 3.25 | 1 |
| 0.077 | 0 | 3.17 | 1 |
| 0.08 | 0 | 3.14 | 1 |

| | | | |
|---|---|---|---|
| 0.212 | 0 | 3.12 | 1 |
| 0.259 | 0 | 3.11 | 1 |
| 0.334 | 0 | 2.94 | 1 |
| 0.041 | 0 | 2.91 | 1 |
| 0.04 | 0 | 2.9 | 1 |
| 0.093 | 0 | 2.89 | 1 |
| 0.166 | 0 | 2.84 | 1 |
| 0.334 | 0 | 2.83 | 1 |
| 0.045 | 0 | 2.82 | 1 |
| 0.077 | 0 | 2.81 | 1 |
| 0.145 | 0 | 2.81 | 1 |
| 0.049 | 0 | 2.8 | 1 |
| 0.11 | 0 | 2.72 | 1 |
| 0.023 | 0 | 2.68 | 1 |
| 0.105 | 0 | 2.67 | 1 |
| 0.259 | 0 | 2.63 | 1 |
| 0.212 | 0 | 2.58 | 1 |
| 0.07 | 0 | 2.21 | 1 |
| 0.016 | 0 | 2.01 | 1 |
| 0.585 | 0 | 1.62 | 1 |
| 0.096 | 0 | 0 | 1 |
| 0.194 | 0 | 0 | 1 |
| 0.08 | 0 | 0 | 1 |
| 0.136 | 0 | 0 | 1 |
| 0.136 | 0 | 0 | 1 |
| 0.101 | 0 | 0 | 1 |
| 0.058 | 0 | 0 | 1 |
| 0.07 | 0 | 0 | 1 |
| 0.166 | 0 | 0 | 1 |
| 0.233 | 0 | 0 | 1 |
| 0 | 0 | 0 | 1 |
| 2.162 | 0 | 0 | 1 |
| 0.059 | 0 | 0 | 1 |
| 0 | 0 | 0 | 1 |
| 0.064 | 0 | 0 | 1 |
| 0.083 | 0 | 0 | 1 |
| 0.129 | 0 | 0 | 1 |
| 0.055 | 0 | 0 | 1 |
| 0.009 | 0 | 0 | 1 |
| 0.025 | 0 | 0 | 1 |
| 0.061 | 0 | 0 | 1 |
| 0.468 | 0 | 0 | 1 |
| 0.08 | 0 | 0 | 1 |
| 0.468 | 0 | 0 | 1 |
| 0.046 | 0 | 0 | 1 |
| 0.585 | 0 | 0 | 1 |

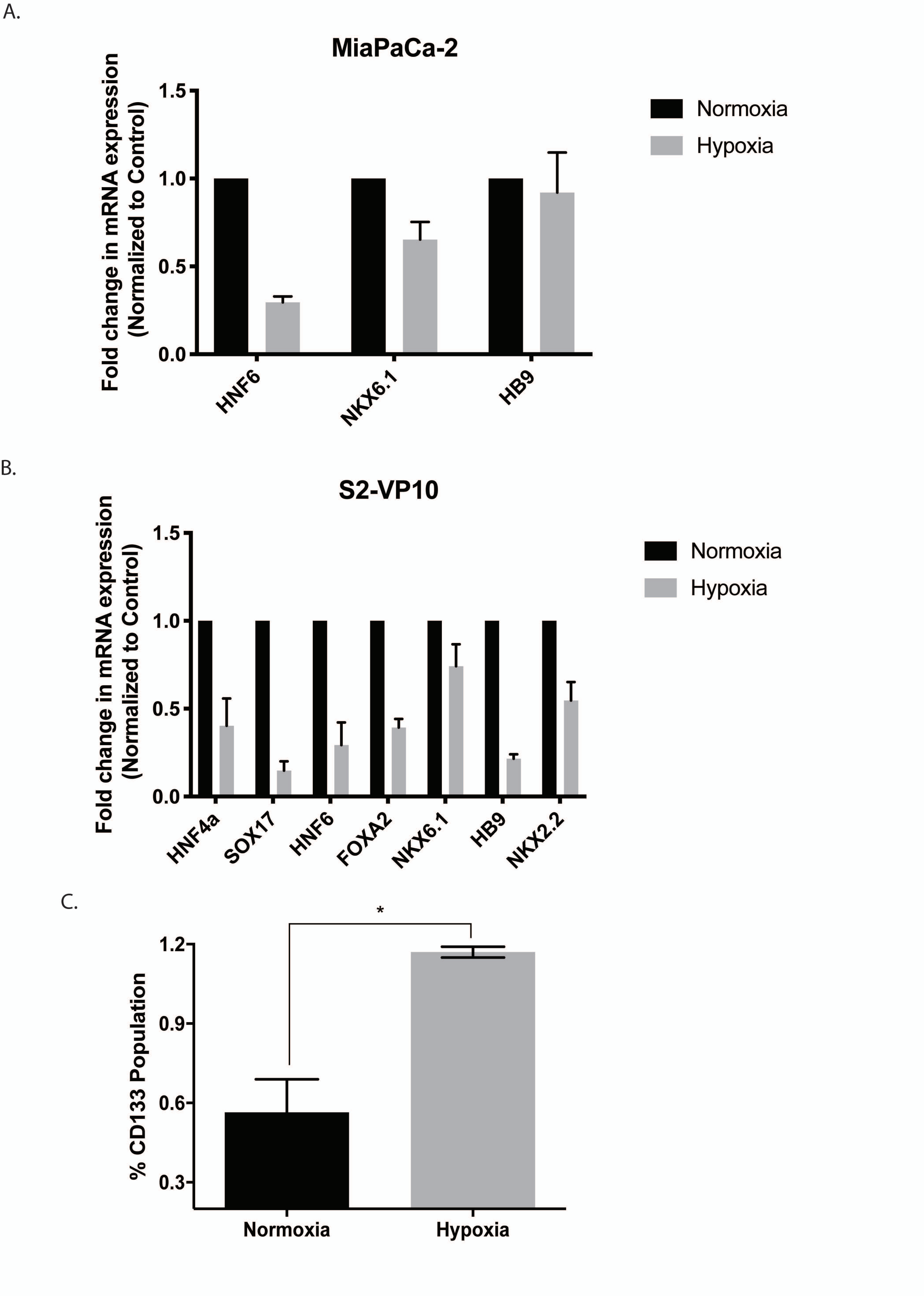

Supplementary Figure 1. Hypoxia suppressed expression of genes involved in differentiation in pancreatic cancer cell line MIA-PACA2(A) and S2VP10 (B). Further, exposure to hypoxia increased CD133+ population in MIA-PACA2 cells

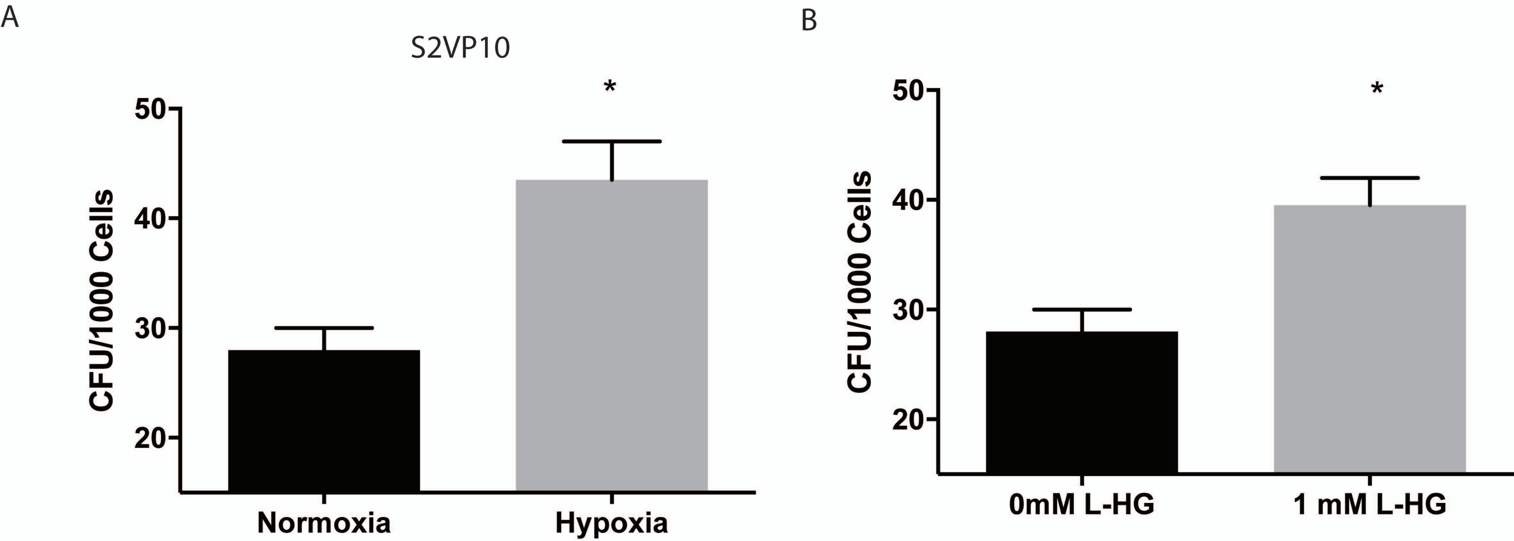

Supplementary Figure 2. Hypoxia (A) and L-2HG (B) increased colony forming units in S2VP10 pancreatic cancer cells

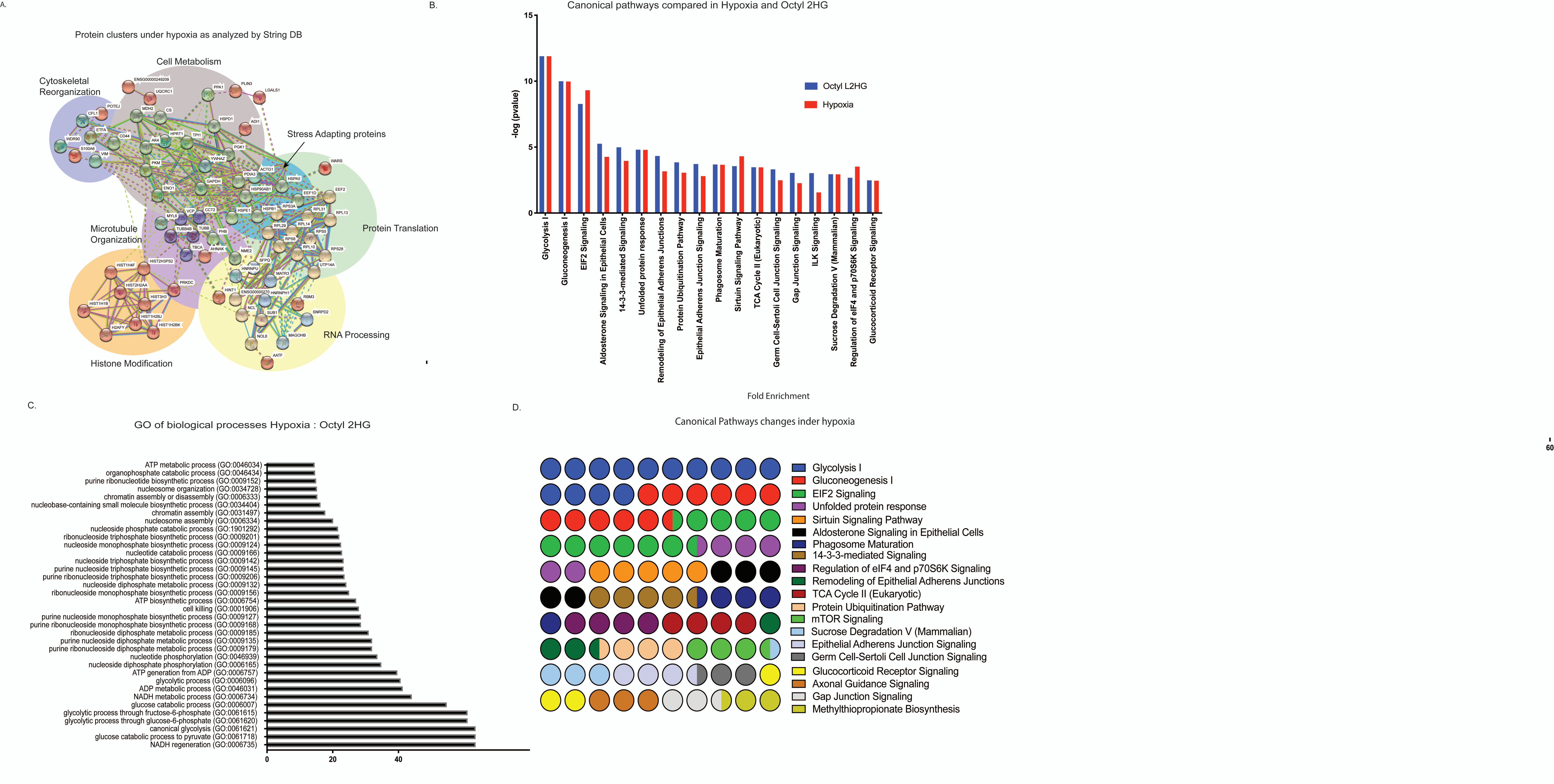

Supplementary Figure 3. ITRAQ analysis of samples under hypoxia compared to normaxia. String DB analysis of protein-protein interation (A). Canonical pathways identified in L-2HG and hypoxia samples (B) GO analysis of biological processes (C) and parts of whole analysis of pathways altered in hypoxia (D).

A.

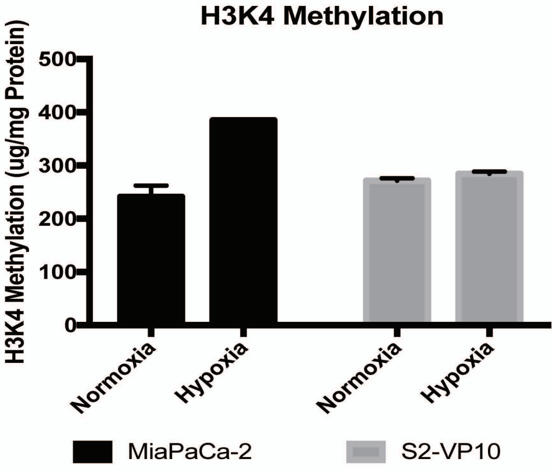

B.

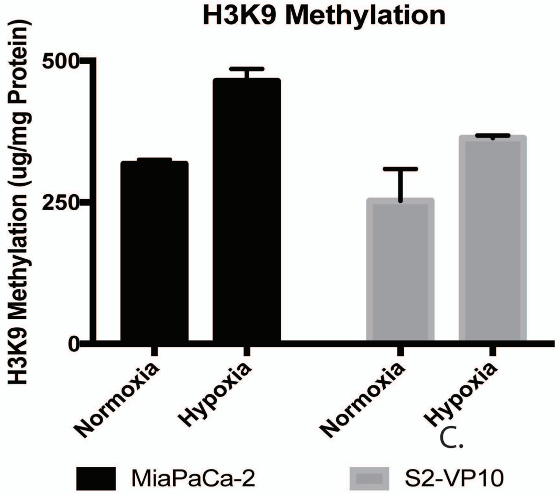

C.

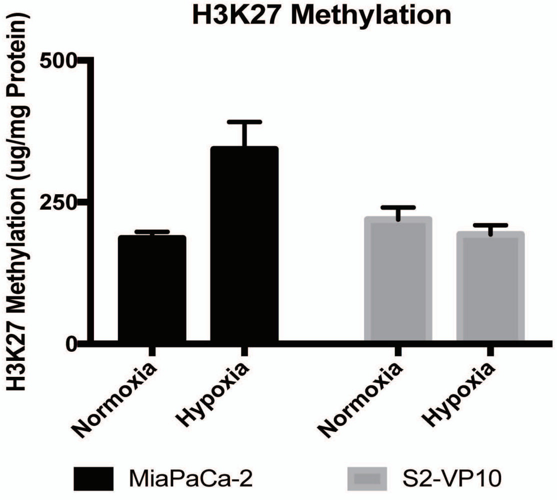

Supplementary Figure 4. methylation status of H3K4 (A), H3K9 (B) and H3K27 (C) upon hypoxia (1% $O_2$ )

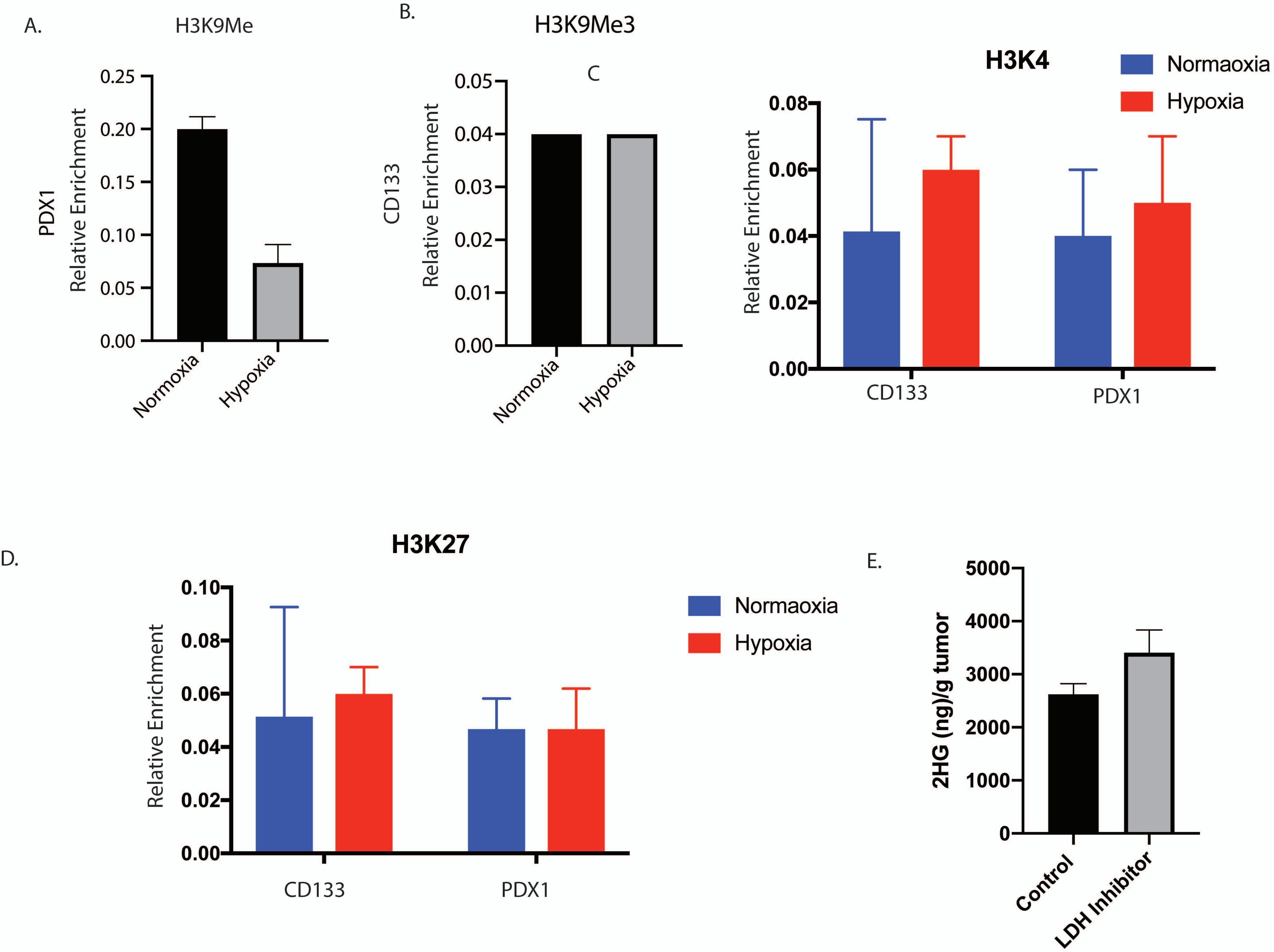

Supplementary Figure 5. PDX1 was not monomethylated during hypoxia indicating repression of gene expression (A) while CD133 not trimethylated during hypoxia indicating its activation (B). Methylation of H3K4 (C) and H3K27 (D) did not change in the promoter of CD133 and PDX1. Inhibition of LDH did not alter intratumoral L2HG (E).
